# Hypermutability of ultraconserved histone genes and its contribution to human disease

**DOI:** 10.64898/2026.06.11.731563

**Authors:** Zongjin Jiang, Jing Xue, Yingchong Teng, Minhuan Lin, Shaobin Lin, Yanmin Luo, Xionglei He, Cai Li

**Affiliations:** State Key Laboratory of Biocontrol, School of Life Sciences, Guangdong Provincial Key Laboratory for Aquatic Economic Animals, Sun Yat-sen University, Guangzhou, Guangdong, China; MOE Key Laboratory of Freshwater Fish Reproduction and Development, School of Life Sciences, Southwest University, Chongqing, China; Department of Obstetrics and Gynecology, The First Affiliated Hospital of Sun Yat-sen University, Guangzhou, Guangdong, China; Guangdong Provincial Clinical Research Center for Obstetrical and Gynecological Diseases, Guangzhou, Guangdong, China

## Abstract

Histone genes are among the most evolutionarily conserved sequences in eukaryotes, reflecting their essential role in nucleosome architecture. Intriguingly, our analysis reveals unexpectedly high germline mutation rates in replication-dependent histone genes across vertebrates. We provide multiple lines of evidence suggesting that this hypermutability is largely driven by off-target mutagenic effects of AID/APOBEC enzymes, representing an unintended fitness cost associated with vertebrate immune system innovation. To characterize the impact of these mutations, we develop HistMTR, a paralog-aware missense constraint framework that outperforms existing tools in predicting the pathogenicity of histone mutations. Applying this framework to large-scale human genomic data, we demonstrate that both coding and regulatory histone mutations contribute to a spectrum of diseases, including developmental disorders and reproductive failure. Analyses of allele transmission ratios and the distribution of fitness effects indicate intense ongoing negative selection against deleterious histone mutations in humans. Together, these findings uncover a hidden vulnerability of ultraconserved histone genes and establish HistMTR as an important framework for characterizing the underappreciated role of histone mutations in human disease.

## Introduction

Germline mutation rates across the genome are highly heterogeneous, shaped by many factors, including local sequence context, replication timing, chromatin architecture, and transcriptional activity^1–3^. While recent advances have enabled fine-scale mapping of mutation rate variation^4–6^, the biological significance of hypermutated genes—loci with mutation rates substantially exceeding genomic background—remains poorly understood. Two recent studies revealed that some localized mutational hotspots arise from clonal expansions driven by positive selection in spermatogonia^7,8^. Yet many hypermutated loci show no evidence of selection-driven expansion, suggesting that additional mechanisms, acting independently of positive selection, remain to be discovered.

In this context, our previous work revealed that histone genes exhibit markedly elevated germline mutation rates in human populations—an unexpected finding given their essential functions and exceptional evolutionary conservation^5^. This observation prompted us to examine histone genes more closely. We found that the histone genes with elevated mutation rates are those encoding replication-dependent histones (RDHs). RDHs, also referred to as canonical histones, comprise five families (H2A, H2B, H3, H4, and H1) and are synthesized during S phase to package newly replicated DNA. Their coding sequences are among the most conserved across eukaryotic lineages^9,10^. This conservation reflects their indispensable role in chromatin assembly, where even subtle amino acid changes can disrupt nucleosome stability, epigenetic regulation, and genome integrity^11–13^. Thus, the observation that RDH genes are hypermutable in humans is counter-intuitive and warrants a thorough investigation of its underlying mutagenic mechanisms and consequences.

One plausible source of RDH gene hypermutation is off-target activity of ‘activation-induced cytidine deaminase/apolipoprotein B mRNA-editing enzyme catalytic polypeptide-like’ (AID/APOBEC) protein family. These enzymes, which evolved to diversify immune repertoires in vertebrates, have been shown to deaminate cytosines in transient single-stranded DNA exposed during transcription, particularly in highly expressed genes^14,15^. Transfer RNA loci, for example, experience unusually high mutation rates that have been attributed in part to transcription-associated AID/APOBEC mutagenesis^15^. Given that RDH genes are among the most highly expressed during S phase, they may be similarly vulnerable to this off-target activity, though this hypothesis remains untested. In addition, RDH genes cluster in three genomic loci in humans (6p22, 1q21, 1q42), lack introns, and undergo unique 3′ end processing^16^—features that may influence their mutagenic susceptibility.

Histone mutations have been intensively studied in the somatic context, particularly in cancer, where they are known as ‘oncohistone’ mutations. Nacev et al.^17^ catalogued the breadth of oncohistone mutations, while mechanistic follow-up studies demonstrated that these variants could remodel chromatin landscapes and perturb cell fate decisions^12^. Yet germline mutations in histone remains comparatively underexplored. Only recently have investigators begun to characterize ‘histonopathies’, Mendelian disorders caused by germline mutations in histone genes. Knapp et al.^18^ summarized the emerging evidence for histone-driven developmental syndromes, and Lubin et al.^19^ extended this landscape through deep phenotyping of 192 individuals with germline histone mutations. A broader clinical review^20^ catalogued dozens of histone-related disorders, though only a subset are caused by direct histone gene mutations. These studies collectively underscore that many pathogenic histone mutations and their associated phenotypes remain undiscovered.

These observations highlight four critical knowledge gaps that motivate this study. First, it remains unclear whether RDH hypermutation in germline is a human-specific phenomenon or a conserved feature across many species. Second, the underlying mutational processes driving elevated mutation rates in RDH genes are unknown. Elucidating phylogenetic distribution of RDH hypermutation and associated mutational processes would shed light on whether this vulnerability arose with lineage-specific mutagenic mechanisms or reflects a deeper, conserved genomic property. Third, predicting functional effects of histone mutations is hampered by the limitations of conventional computational predictors, which often underperform in histone genes due to their short sequences and paralog redundancy, impeding accurate prioritization of pathogenic variants. Fourth, while a small number of germline histone mutations have been linked to severe Mendelian disorders (so-called histonopathies), the broader phenotypic consequences of both coding and regulatory variants remain largely uncharacterized.

In this study, we perform a thorough investigation with diverse omics datasets to address these fundamental questions. We investigate the cross-species conservation of histone hypermutation, delineate the mutational mechanisms responsible for elevated mutation rates, develop an improved framework for predicting functional impact of missense histone mutations, and characterize the disease associations of coding and noncoding histone mutations. These findings provide important insights into intragenomic mutation rate variation, evolutionary constraints on histone genes, and the broader landscape of human histonopathies.

## Results

### Replication-dependent histone genes exhibit elevated mutability in vertebrate genomes

Building upon prior evidence that histone genes were enriched among human coding genes with high mutation rates^5^, we systematically investigated the mutational landscape of all histone genes in the human genome. As relatively few *de novo* mutations (DNMs) were available and extremely rare variants are a reasonable proxy for DNMs^4,5^, we used rare variant density for mutation rate comparison. With an allele frequency (AF) cutoff of 1 × 10^-4^, we extracted 461 million rare single-nucleotide variants (SNVs) and 31 million rare short indels (1-3 bp) from the gnomAD v4.0 database^21^. A total of 19,100 autosomal protein-coding genes were categorized into three groups: RDH genes (n = 70), replication-independent histone (RIH) genes (n = 19), and other genes (n = 19,011). Genes were further restricted to those with more than 50 callable nucleotides in subsequent analyses based on coverage and mappability (Methods).

The average SNV and indel rare variant densities of human RDH genes were about two folds higher than genomic average (**Fig. 1a**), while RIH genes showed levels comparable to genomic average. All five RDH gene families (H1, H2A, H2B, H3, and H4) consistently showed elevated rare variant densities (**Fig. 1a**). Similar patterns were also observed in somatic mutations (**Supplementary Fig. 1**). By separating SNV subtypes, we found that the elevated mutation burden was mainly from mutations of non-CpG sites (**Supplementary Fig. 2**). The reduced rare-variant density at CpG sites in RDH genes was consistent with their lower germline methylation levels relative to other genes (**Supplementary Fig. 3a**), which would generate fewer deamination-derived mutations. Although the small number of DNMs was insufficient for gene-level comparison, RDH genes exhibited higher group-level *de novo* mutation rate (**Supplementary Fig. 4**), further supporting their elevated mutability.

**Fig. 1:**
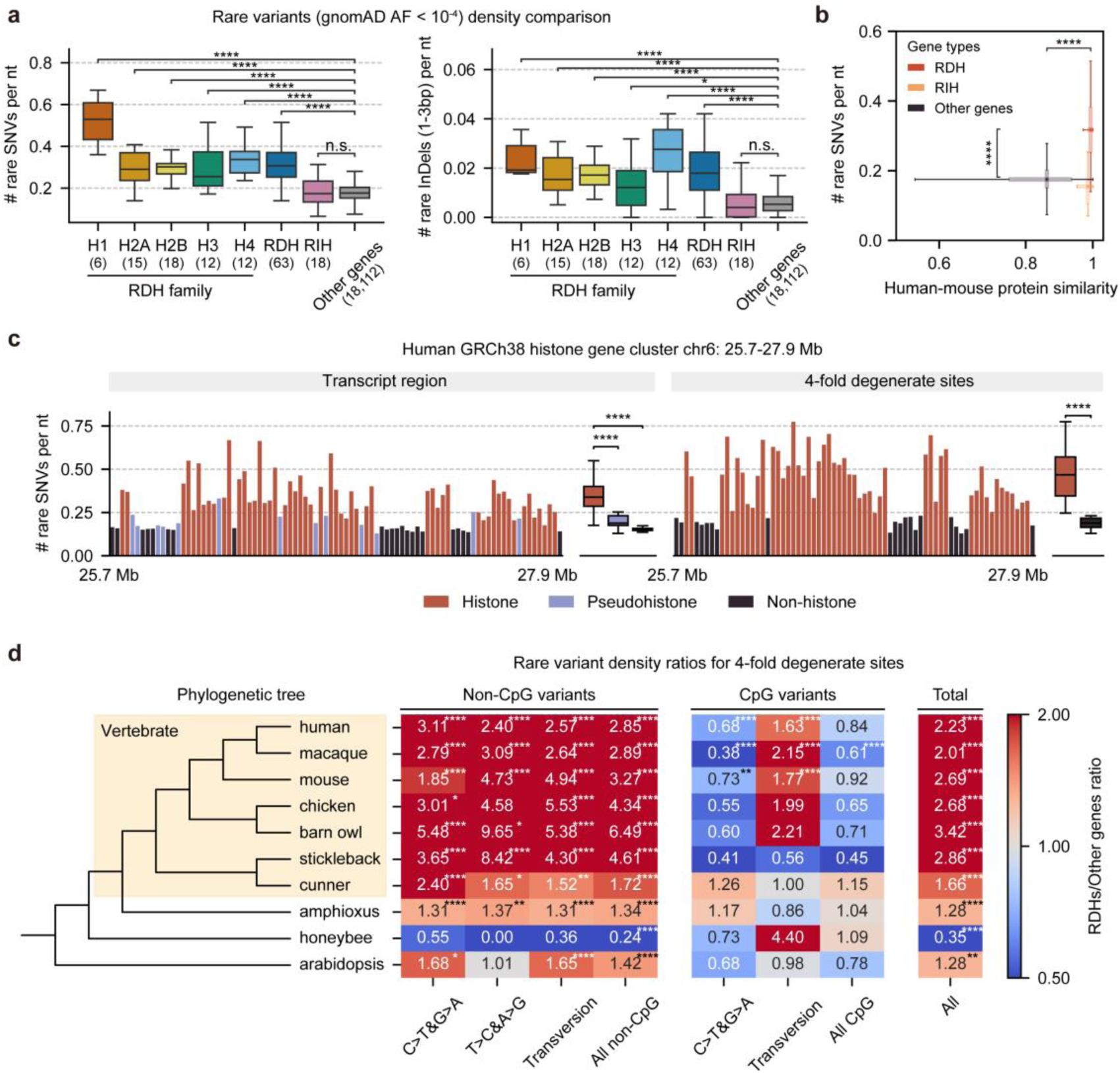
RDH genes exhibit elevated mutability. **a**, Bars represent the mean rare variant density per nucleotide in coding sequences for each histone family, the aggregated RDHs group (H1–H4), RIHs, and other genes. RDH denotes the combined set of canonical replication-dependent histone families (H1, H2A, H2B, H3, and H4). The number of genes in each set is indicated in brackets. *P* values were obtained by two-sided Mann–Whitney U tests. Box-plot elements: center line, median; box limits, first and third quartiles; whiskers, 1.5× interquartile range. **b**, Distributions of protein-sequence similarity and human rare SNV density for human–mouse one-to-one orthologs. *P* values were obtained by two-sided Mann–Whitney U tests. **c**, Rare SNV density across the largest human histone gene cluster, shown per gene for transcript regions and 4-fold degenerate sites. Genes are arranged by genomic position of their TSS. Box plot elements and statistics are as in **b**. **d**, Heatmap displaying the ratio of rare SNV density at 4-fold degenerate sites in RDH genes relative to genomic background (other genes) across ten species, summarized by substitution class (non-CpG, CpG, and all). A simplified phylogenetic tree (left) was generated with iTOL (itol.embl.de). *P* values were calculated using two-sided Fisher’s exact tests. *: 0.01 < *P* ≤ 0.05; **: 0.001 < *P* ≤ 0.01; ***: 0.0001 < *P* ≤ 0.001; ****: *P* ≤ 0.0001.

Notably, 41.3% of RDH genes ranked among the top 1% highest of all genes for rare SNV density in coding sequences. While RDH genes showed significantly higher rare SNV density in human populations, they maintained extremely high protein similarities (a median of 99.3%) and ultra-low ratios of nonsynonymous to synonymous substitution rates (dN/dS) in human-mouse orthologs (**Fig. 1b**; **Supplementary Fig. 5**). 21.8% of surveyed RDH genes had a dN/dS of 0, markedly higher than the genome-wide proportion of 1.6%. This indicates RDH genes underwent strong purifying selection against many new missense mutations derived from the elevated mutability, consistent with the extreme sequence conservation of histones^9^ (**Fig. 1b**; **Supplementary Fig. 6**). RDH genes also showed significantly higher synonymous substitution rates (dS), suggesting their inherent hypermutability over evolutionary timescales (**Supplementary Fig. 5**).

As background selection reduces neutral diversity at linked sites through the purging of deleterious alleles, the rare-variant–based metric above likely underestimated the true elevation of mutation burden in RDH genes. Indeed, the less conserved H1 family showed higher variant density than other four RDH families (**Fig. 1a**), likely due to weaker background selection in H1 genes. Furthermore, the central gene body regions of RDH genes showed lower rare variant densities than upstream and downstream regions (**Supplementary Fig. 2**), also suggesting the impact of background selection. When using rare variants at 4-fold degenerate sites to minimize the impact of selection, we observed increased difference in mutation burden between RDH genes and other genes (**Fig. 1c**) and reduced difference between H1 and other RDH gene families (**Supplementary Fig. 7**).

To assess whether RDH gene hypermutation is human-specific or a conserved across taxa, we analyzed histone mutation burden in nine other representative species using rare variants (AF < 0.01) from population data (Methods). To mitigate selection bias, we computed the RDH-to-others variant density ratios for 4-fold degenerate sites. RDH genes consistently showed elevated non-CpG variant densities relative to genomic background across all surveyed vertebrates, with fold-increases ranging from 1.72 to 6.49 (**Fig. 1d**). In many invertebrates (e.g., insects and mollusks) and some vertebrates (e.g., reptiles and some fish), RDH genes form tandem arrays^22^, complicating genome assembly and variant calling and thus limiting analysis. To circumvent this, we examined several non-vertebrate species lacking such tandem arrays. We observed a slight elevation in mutation rates in amphioxus (a close vertebrate relative) and *Arabidopsis thaliana*, but no increase in the honeybee. Taken together, these data demonstrate that RDH gene hypermutability is a highly conserved pattern across vertebrates, but not in other lineages.

### Hypermutability of RDH genes is not explained by genomic context or expression level alone

Human RDH genes are organized into three genomic clusters (55 genes at 6p22, 11 at 1q21, and 3 at 1q42), with the 6p22 cluster encompassing most of the genes. To investigate the underlying mutational mechanisms, we first assessed whether increased mutability is a general feature of histone gene cluster regions by comparing RDH genes and nearby non-histone genes. We analyzed rare SNV density at the genic regions and specifically at 4-fold degenerate sites to minimize influence of selection. Within the cluster regions, RDH genes showed significantly higher SNV density than other genes, whereas pseudohistone genes were less mutable and less expressed (**Fig. 1c**; **Supplementary Fig. 8**). This suggests a contribution from transcription-associated mutational processes^14^.

Given that gene expression and sequence context can influence mutation rates^5,14^, we performed additional controls to investigate these factors. Human non-histone protein-coding genes were subdivided into germline-expressed (n = 15,861) and germline-unexpressed (n = 3,150) categories (Methods). The 3-mer composition of each group was rescaled to match that of expressed genes for eliminating context bias (**Supplementary Fig. 9**). We further applied a calibration to correct for potential recurrent mutations, yielding calibrated variant densities (**Supplementary Fig. 10**; Methods). Even after these adjustments, RDH genes retained higher rare variant density for all non-CpG 1-mers and for most 3-mer classes at both transcript-level and 4-fold degenerate sites, compared to either expressed or unexpressed non-histone genes, whereas the mutability of the latter two was nearly the same (**Fig. 2a**; **Supplementary Fig. 11**). This underscores that RDH genes’ elevated mutability is not merely caused by their specific sequence context. Moreover, while germline expression generally exhibited a weak negative correlation with mutability in non-RDH genes (Spearman’s rho = −0.069, *P* = 2.39 × 10^-20^), it showed a significant positive correlation with rare variant density (Spearman’s rho = 0.425, *P* = 0.0006) at 4-fold degenerate sites in RDH genes (**Supplementary Fig. 12**). This reversal indicates that the relationship between expression and mutability is more complex in RDH genes, and suggests that their high expression, in conjunction with other unique features, contributes to their elevated mutation rates rather than acting alone.

**Fig. 2:**
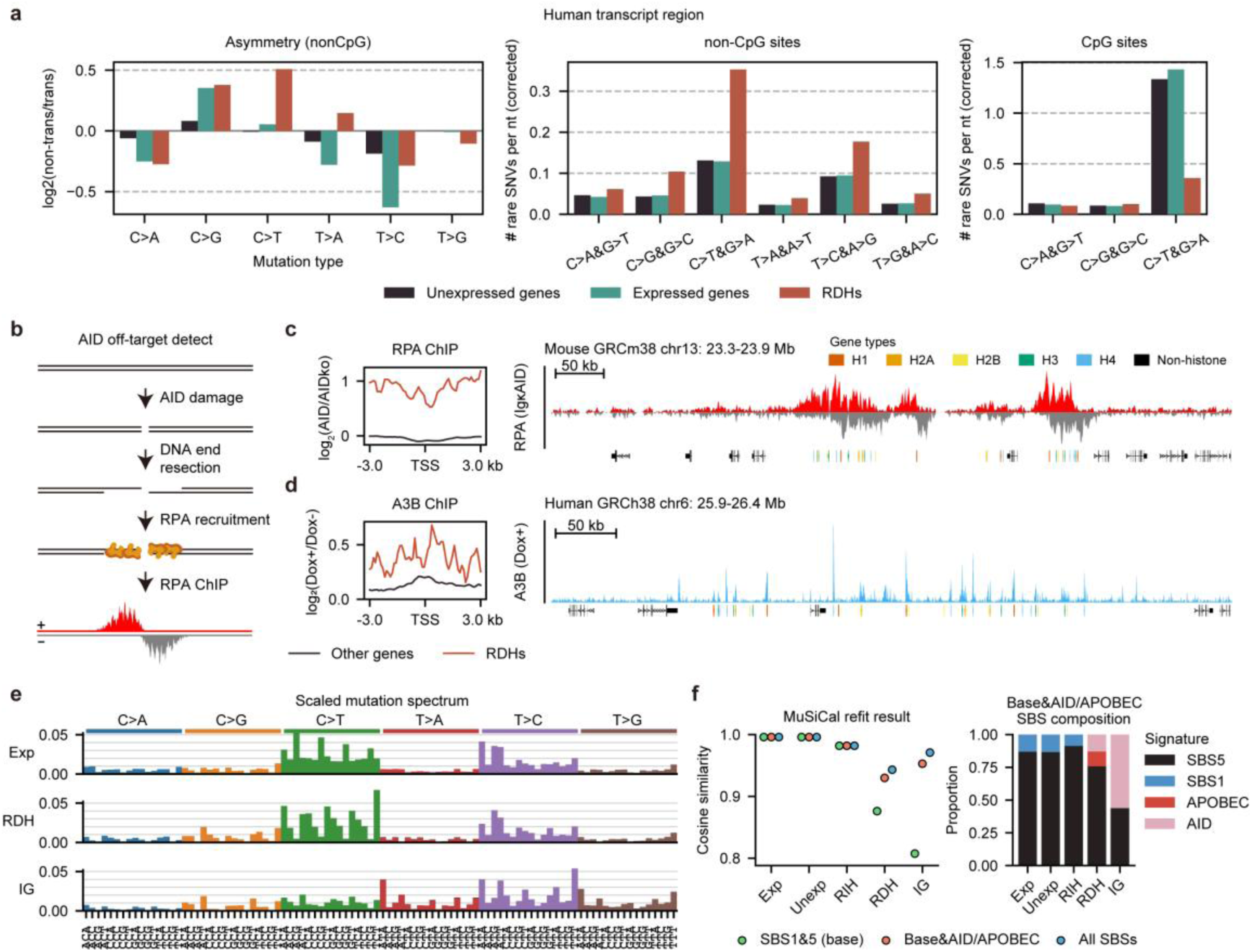
AID/APOBEC activity contributes to mutagenesis in RDH genes. **a,** Corrected SNV densities (1-mer) and asymmetry scores for each gene group. Analyses were restricted to transcript regions; CpG and non-CpG sites are shown separately. Asymmetry scores were computed using non-CpG sites. **b,** Schematic of RPA recruitment following off-target AID-mediated DNA damage (adapted from Qian et al.^24^). **c-d,** Genomic profiles of AID/APOBEC activity within the largest RDH gene cluster in mouse (**c**) and human (**d**), respectively. **c**, Genomic profiles of strand-specific RPA-ChIP-seq RPKM values in mouse B cells with the IgκAID transgene. **d**, Genomic profiles of APOBEC3B (A3B) activity MCF10A cells. The left panel shows the SES-scaled log_2_(treatment / control) coverage ratio; the right panel shows – log_10_(*P* value) for the doxycycline (Dox)-induced sample. **e,** Scaled mutation spectra for expressed genes, RDH genes, and IG genes. The y-axis indicates the proportion of mutations within each 3-mer context. **f,** Refitting of mutational signatures using MuSiCal under three signature combinations (left), and decomposition plot showing the contribution of each SBS category under the ‘clock-like + AID/APOBEC’ model (right).

### Mutational patterns and functional genomic assays indicate the involvement of AID/APOBEC activity

To investigate the potential drivers of elevated mutability in RDH genes, we analyzed strand asymmetry and the 1-mer mutation spectrum (Methods), as these might provide signals for specific mutagenic processes. RDH genes displayed a pronounced enrichment of C>T substitutions relative to expressed genes, with asymmetry scores indicating preferential accumulation on the non-transcribed strand (**Fig. 2a**). This mutational pattern is consistent with the activity of the AID/APOBEC cytidine deaminase family, which is known to act in transcriptional contexts and produce excess C>T transitions in other highly transcribed loci such as transfer RNA genes^15^.

To assess the role of AID/APOBECs in RDH gene hypermutation, we further analyzed related functional genomic data. Because no relevant data profiling AID binding patterns in the human genome was available, we utilized RPA-ChIP data from mouse activated B cells to investigate AID activities^23^. In the RPA-ChIP experiment, 53BP1 knockout enhances RPA recruitment in the presence of the Igκ-AID transgene, and the resulting strand-asymmetrical accumulation of RPA serves as a proxy for AID-mediated damage^23^ (**Fig. 2b**). Visualization of strand-specific coverage confirmed extensive AID targeting in RDH genes which was previously reported by Qian et al.^24^ (**Fig. 2c**; **Supplementary Fig. 13a**). This signal was absent in AID-knockout B cells, confirming its AID dependency (**Fig. 2c**; **Supplementary Fig. 14**). Interrogation of APOBEC3B (A3B) ChIP-seq data from human MCF10A cells^25^ also revealed a significantly higher log2 coverage ratio (doxycycline-induced vs. uninduced control) in RDH genes than in other genes, suggesting enriched binding of A3B at these regions (**Fig. 2d**). Furthermore, genome-wide A3B signals revealed strong colocalization with RDH genes (**Fig. 2d; Supplementary Fig. 15 and Supplementary Fig. 13b**). Although APOBEC3A (A3A) is considered the primary source of endogenous APOBEC-signature mutations, A3B shares a similar preferred motif and has been shown to contribute to mutagenesis and correlate with activity of A3A^26,27^. Furthermore, ELOF1—an essential factor for transcription-coupled AID deamination^28^—also showed increased signal within RDH genes (**Supplementary Fig. 16a**), providing additional supportive evidence.

Because AID/APOBEC activity can induce DNA double-strand breaks (DSBs) and activate downstream damage responses^24,29,30^, we reanalyzed published ChIP-seq data for the DSB-associated marker γH2AX. RDH genes indeed exhibited a marked enrichment of γH2AX signal relative to other genes in both human and mouse (**Supplementary Fig. 16b**). This enrichment is consistent with off-target AID/APOBEC activity, although alternative sources of DSBs cannot be excluded.

### Quantifying the contribution of AID/APOBECs by mutational signature analysis

Since AID/APOBEC enzymes deaminate cytosines within preferred sequence motifs, we next analyzed the mutation spectra to infer underlying mechanisms. We scaled mutation spectrum of each gene group to match the whole-genome 3-mer context (**Fig. 2e**) and used MuSiCal^31^ to fit three nested combinations of COSMIC signatures: a baseline ‘clock-like’ set (SBS1+SBS5, comprising the majority of mutations across different tissues^32^), an expanded set including AID/APOBEC-related signatures, and a global set of all SBS signatures (**Fig. 2f**; Methods). Immunoglobulin (IG) genes, known AID targets, were included as a positive control (n = 214 genes). Gene groups other than RDHs and IGs were well explained by clock-like signatures alone (**Fig. 2f**; cosine similarity > 0.98). In contrast, both RDHs and IGs showed poorer fits (RDHs: 0.88; IGs: 0.81). The inclusion of five AID/APOBEC-associated SBSs substantially improved the cosine similarity (RDHs: 0.93; IGs: 0.95), bringing it near the fit achieved using all signatures (RDHs: 0.94; IGs: 0.97). Decomposition analysis indicated that AID/APOBEC-related signatures contributed approximately 24.2% of mutations in RDH genes (**Fig. 2f**; **Supplementary Table 1a**). This proportion might represent an underestimate, as some AID/APOBEC-induced lesions could be classified as SBS5 due to repair errors^33,34^. A permutation test confirmed that adding AID/APOBEC-related SBSs significantly improved the fit for RDHs and IGs, but not for other gene groups (**Supplementary Table 1a**; Methods). Results were consistent across different variant datasets and genomic regions, mitigating concerns of somatic artefacts, dataset bias, gene length and codon selection (**Supplementary Table 1b-f**).

### Quantifying pathogenicity of protein-altering mutations in RDH genes

Given the extreme conservation of RDH sequences, many protein-altering variants in RDH genes are likely pathogenic. Although mutations in RDH genes have been implicated in human diseases, including cancer and developmental disorders^35–38^, RDH genes are generally under-represented in disease-association studies. One plausible explanation is that similar pathogenic phenotypes can arise from mutations in different RDH paralogs^37–39^, whereas most analytical frameworks focus on signals from individual genes. Furthermore, the short coding sequences of RDH genes limit the statistical power of common gene-based approaches^40^, necessitating a more effective framework for quantifying their pathogenic potential.

Given the strong sequence and functional conservation within each RDH family, we developed a missense constraint metric—termed HistMTR—based on alignments of paralogs and polymorphism data from each RDH family. Missense tolerance ratio (MTR) is analogous to the ratio of nonsynonymous to synonymous polymorphism rate (pN/pS) but is more robust when variant counts are low^41^. For HistMTR calculation, we aligned all paralogs within each RDH family and computed site-level MTRs using all variants at each aligned codon (**Fig. 3a**; Methods). Variant counts were weighted by gene-specific pN/pS ratios, with greater weight assigned to genes under stronger constraint (lower pN/pS). We further computed a gene-level MTR for each RDH gene using its own polymorphic sites. Site-level and gene-level MTRs were then integrated to generate a HistMTR value for each codon position in each RDH gene (**Supplementary Table 2**). This framework leveraged million-exome coding variation from RGC-ME^42^ together with context-aware mutation rates predicted by MuRaL^5^ (**Fig. 3a**; Methods). Analyses were restricted to core RDH families (H2A, H2B, H3, and H4), as the greater sequence divergence of H1 genes limited alignment accuracy.

**Fig. 3:**
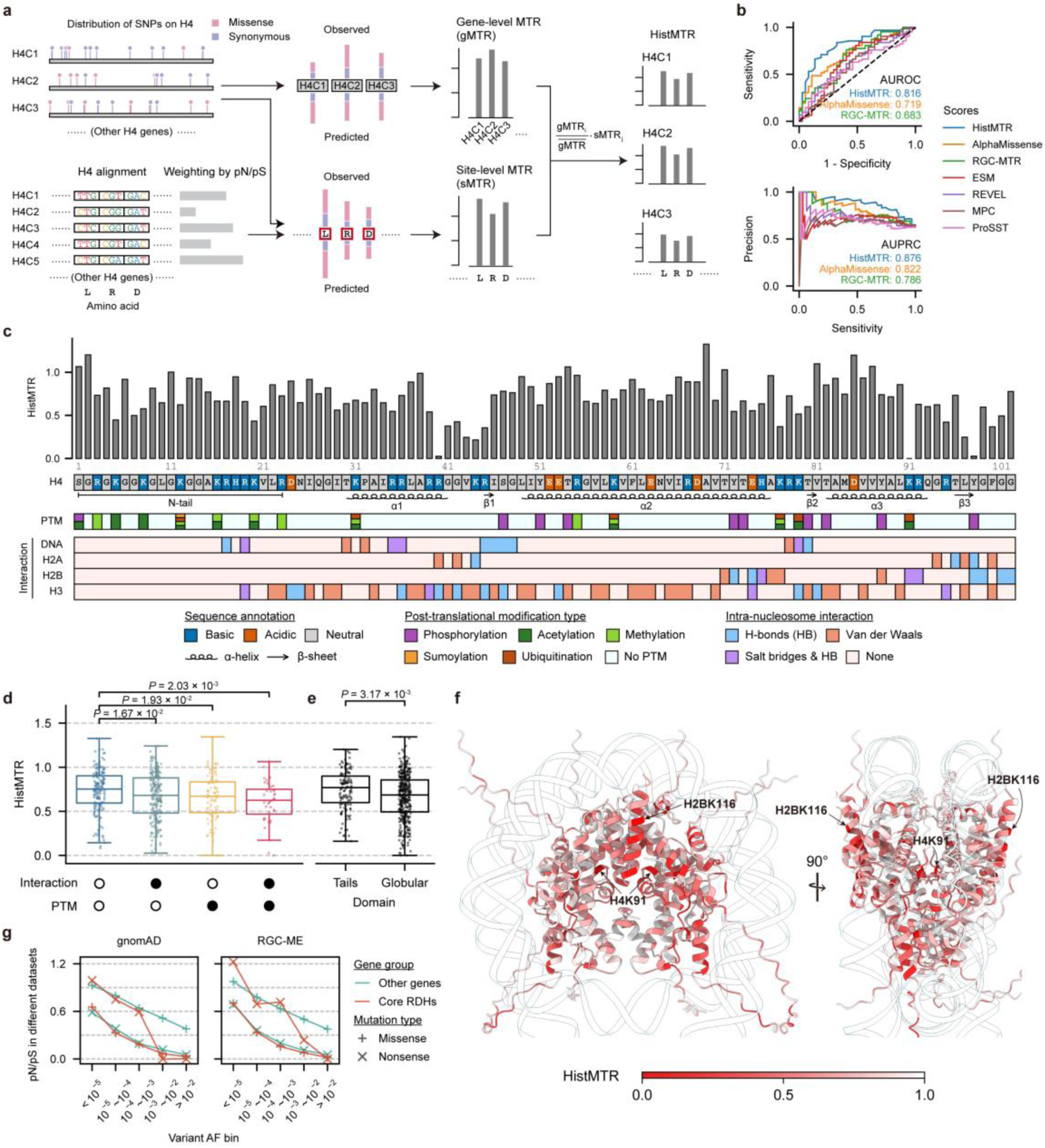
Pathogenicity prediction of amino acid mutations in core RDHs. **a,** Schematic of the HistMTR framework. HistMTR integrates site-level and gene-level missense tolerance ratios (MTRs) to assess constraint at each amino acid position in RDH genes: site-level MTRs are derived from variants across all aligned paralogs, weighted by gene-specific pN/pS ratios (lower pN/pS = higher weight), while gene-level MTRs are based on polymorphic sites within the individual gene (Methods). **b,** Benchmarking of deleteriousness predictors on curated RDH mutations (n = 99) from Bagert et al.^12^. AUROC/AUPRC for the three top-performing predictors are shown. **c,** Site-specific mean HistMTR values for the H4 protein, with annotations indicating consensus amino acids, post-translational modifications (PTMs), and intra-nucleosome contacts. Results for other core RDH families are shown in **Supplementary Fig. 18**. **d**, Comparison of HistMTR values between sites grouped by presence (filled circles) or absence (open circles) of PTMs or intra-nucleosome contacts. *P* values were calculated using two-sided Mann–Whitney U tests. **e,** Distribution of HistMTR for residues in histone tails versus globular domains. *P* value was calculated using two-sided Mann–Whitney U test. **f**, Nucleosome structures colored by HistMTR (red = lower missense tolerance); H1 is omitted and DNA is shown in transparent mode. Two sites with zero HistMTR (H2BK116 & H4K91) are marked. **g,** Missense and nonsense pN/pS for core RDH genes versus other genes across AF bins, estimated from gnomAD whole-genome and RGC-ME exome datasets.

We evaluated HistMTR’s performance in distinguishing putative pathogenic histone mutations derived from an independent functional assay dataset^12^, against six established metrics: AlphaMissense^43^, RGC-MTR^42^ (based on sliding genomic windows instead of homolog alignments), ESM-1v^44^, REVEL^45^, MPC^46^, and ProSST^47^ (Methods; mutant data detailed in **Supplementary Table 3**). HistMTR demonstrated superior performance, achieving an area under the receiver operator curve (AUROC) of 0.816 and an area under the precision-recall curve (AUPRC) of 0.876, substantially outperforming the second best tool, AlphaMissense (AUROC = 0.719; AUPRC = 0.822) (**Fig. 3b**; **Supplementary Table 4**). Moreover, HistMTR was uniquely effective at discriminating both moderately and strongly deleterious mutations, whereas other metrics showed limited capacity to identify variants with intermediate effects (**Supplementary Fig. 17**). These results demonstrate that integrating information across paralogs significantly enhances pathogenicity prediction, establishing HistMTR as an improved measure of functional constraint for residues in core histones.

HistMTR values exhibited marked variation across adjacent sites within RDHs (**Fig. 3c** for H4; **Supplementary Fig. 18** for other families; **Supplementary Table 5**), highlighting the framework’s capacity to quantify functional importance at single-residue resolution. By integrating structural and post-translational modification (PTM) information, we compared the functional constraints of individual residues (**Fig. 3d, e**). Overall, sites with PTMs and intra-nucleosome contacts exhibited significantly lower HistMTR values (i.e., lower missense tolerance) than those without such features (**Fig. 3d**), consistent with the expectation that mutations at residues directly involved in nucleosome structure or PTMs tend to be more deleterious. Furthermore, residues within the structured globular domains were significantly less mutation-tolerant than those in the unstructured tails (**Fig. 3e**). Surface-exposed helical residues displayed generally low HistMTR values (**Fig. 3f and Supplementary Fig. 19**), likely due to their potential roles in inter-nucleosome interactions or regulatory protein binding.

Among all RDH sites, two residues—H2BK116 and H4K91—exhibited an HistMTR of 0 (**Fig. 3f**), indicating complete absence of missense variants across paralogs in the surveyed population. H4K91, located on the third helix of H4 and a site for two PTMs and contact with H2B (**Fig. 3c, f**), is a known pathogenic site. Distinct amino acid substitutions at this residue have been identified in three patients with severe developmental disorders, and functional assays in zebrafish confirmed that these variants destabilize chromatin and cause genome instability^39^. H2BK116, positioned within the C-terminal helix of histone H2B, carries three known PTMs (**Supplementary Fig. 18b**). Although no germline or somatic pathogenic variants have yet been reported at this position, prior work demonstrated that BARD1—a critical mediator of DNA damage repair—interacts with H2BK116^48^, suggesting an essential role in nucleosome recognition by repair machinery.

We extended our investigation to nonsense variants, typically classified as loss-of-function (LoF) alleles. Due to their extreme rarity, we aggregated these variants to calculate a grouped nonsense pN/pS, contrasted with a missense pN/pS (Methods). As expected, both ratios decreased with increasing AFs, consistent with longer exposure to purifying selection in high-AF variants (**Fig. 3g**). Strikingly, in core RDH genes, the pN/pS ratio was lower for missense changes than for nonsense changes—a pattern opposite to that seen in other genes. This trend was consistent across independent datasets and various histone families (**Fig. 3g**; **Supplementary Fig. 20**). These findings suggest that core RDH genes are more intolerant of amino acid substitutions than of truncations. This may be explained by a dominant-negative mechanism, wherein missense-mutated histones are incorporated into the nucleosome to ‘poison’ chromatin integrity and drive genome instability, whereas truncated products are likely degraded or excluded from assembly^17^.

### Roles of coding mutations of RDH genes in human disease

Since previous research has cataloged and systematically investigated recurrent coding mutations in histone genes across various cancer types^17^, in this study we concentrated our analysis on the roles of histone mutations in non-cancer diseases. We first reanalyzed variant datasets from large developmental disorder (DD) and autism spectrum disorder (ASD) cohorts to assess whether mutations in core RDH genes are enriched in affected individuals, and then conducted comprehensive association analyses using UK Biobank (UKB) data.

Prior gene-based enrichment analysis by DeNovoWEST^49^ with a large DD cohort did not identify a significant association between DNMs in core RDH genes and DD. However, when comparing DNMs in DD patients against population-derived variants (Methods), we observed a significant enrichment of missense DNMs within the core RDH gene families (**Fig. 4a, top**; **Supplementary Table 6**). Although the magnitude of enrichment in core RDH genes (odds ratio [OR] = 2.72) was lower than that observed for DeNovoWEST-defined DD-related genes (OR = 7.37), it exceeded the genome-wide background (OR = 1.37), suggesting that mutations in core RDH genes confer elevated risk for DD relative to the genomic baseline. Notably, stratifying RDH missense sites into five bins based on HistMTR values revealed markedly stronger enrichment in the two lowest bins (first bin: OR = 19.26; second bin: OR = 4.52; **Fig. 4a**), highlighting the substantial contribution of missense-intolerant RDH sites to DD risk. Similar but less pronounced patterns were observed when stratifying RDH missense sites using other variant effect predictors (**Supplementary Fig. 21**).

**Fig. 4:**
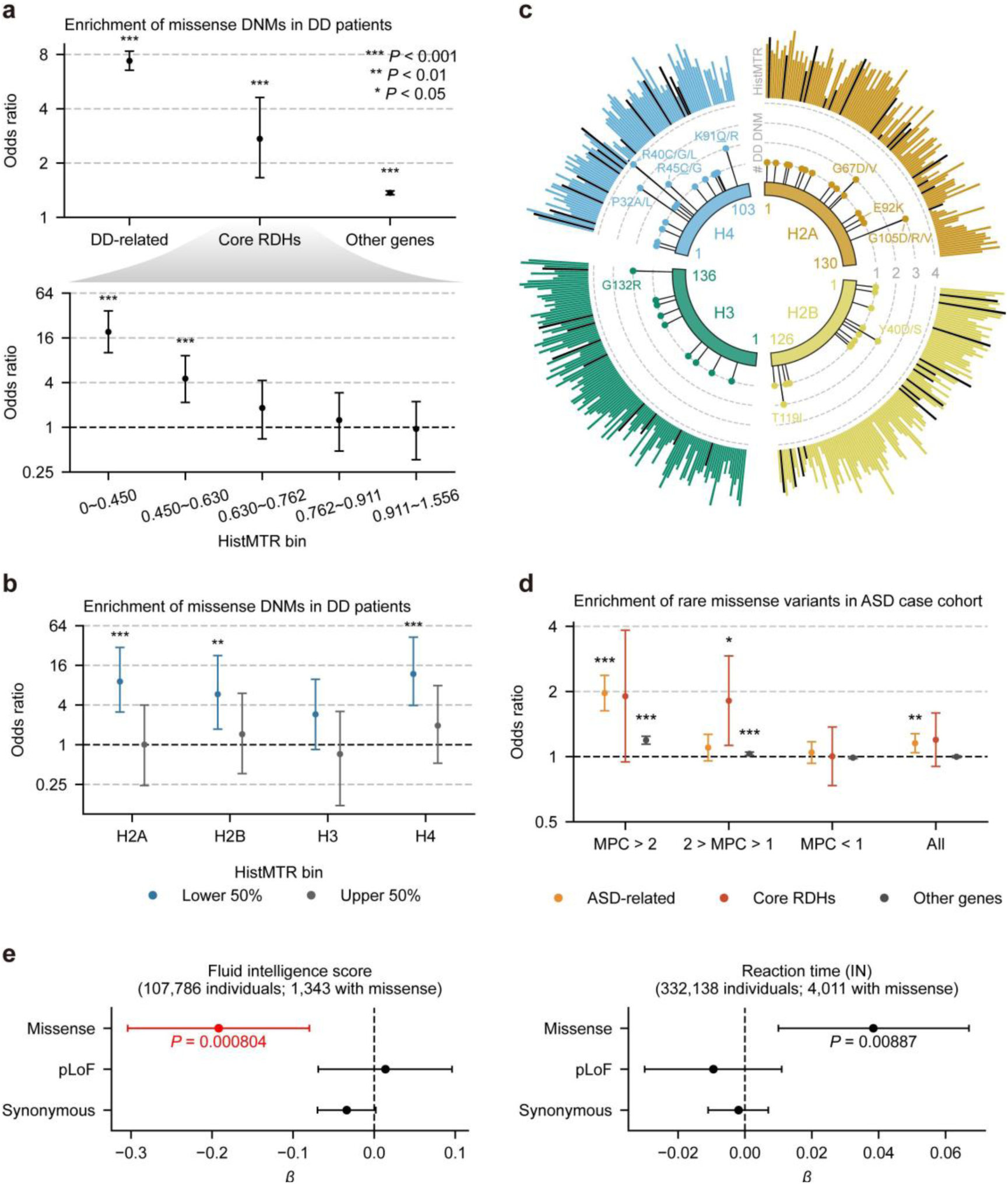
Roles of RDH coding mutations in human disease. **a,** Enrichment of missense DNMs in DD patients for DenovoWEST-defined DD genes, core RDH genes, and other genes. Odds ratios and *P* values were calculated using Fisher’s exact test. For core RDH genes, amino acid sites were further stratified into five bins based on HistMTR values. **b,** Enrichment of missense DNMs in DD patients for core RDH families (H2A, H2B, H3, and H4). Amino acid sites in each RDH family were stratified into two bins based on HistMTR values. Odds ratios and *P* values were calculated using Fisher’s exact test. **c,** Distribution of missense DNMs in DD patients across core RDH families. Amino acid substitutions at sites with recurrent DNMs are shown with labels; substitutions that match DECIPHER ‘Full’ variants are denoted by underlining. HistMTR bars are shown in black where the corresponding sites harbour missense DNMs in DD patients. See **Supplementary Table 8** for details. **d,** Enrichment of rare missense variants in ASD cases, stratified by MPC bins, for TADA-defined ASD genes, core RDH genes, and other genes. Odds ratios and *P* values were calculated using Fisher’s exact test. **e,** Association tests between cognitive phenotypes and three coding variant classes in core RDH genes: putative pathogenic missense, putative LoF, and synonymous variants. *β* values for fluid intelligence represent effect sizes on standardized fluid-intelligence scores (range: 1–13), whereas *β* values for reaction time (inverse-normal transformed) represent effect sizes in standard-deviation units. Error bars represent 95% confidence intervals; bars are colored red if the association remained significant after Bonferroni correction (adjusted *P* < 0.05). Raw *P* values < 0.05 are shown. *: 0.01 ≤ *P* < 0.05; **: 0.001 ≤ *P* < 0.01; ***: *P* < 0.001.

Previous studies have reported specific associations between core RDH genes and DD (compiled in **Supplementary Table 7**), with most associations involving H4 genes^19,37^. Consistent with this, our enrichment analysis revealed that the H4 family possesses the strongest signal among all core RDH families (**Fig. 4b**). However, we also observed significant enrichment for H2A and H2B (**Fig. 4b**), suggesting an underappreciated role for these histone families in the landscape of DD pathology. To further delineate the contribution of individual residues, we mapped these DNMs onto core RDH families (**Fig. 4c**).

Consistent with prior findings, three H4 residues—H4R40 (mean HistMTR = 0.025; hereafter, HistMTR refers to the mean across all paralogs when discussed at the family level), H4R45 (HistMTR = 0.356), and H4K91 (HistMTR = 0)—each associated with ‘Full’ explanatory variants in the DECIPHER^50^ database (**Supplementary Table 7**), exhibited recurrent DNMs in DD patients (**Fig. 4c**; **Supplementary Table 8**). Interestingly, the specific amino acid changes resulting from these DNMs were not always identical to those documented in DECIPHER (**Supplementary Table 8**), suggesting that distinct alterations at the same residue can converge on similar phenotypic outcomes. A similar pattern was observed at H2A E92 (HistMTR = 0.210): while the E92A substitution is classified as ‘Full’ in DECIPHER, we identified a different alteration, E92K (**Fig. 4c**), among DNMs in DD patients. Although this amino acid change can result from 14 different SNVs in human genome, its complete absence from the RGC-ME database (which covers 32.2% of all possible missense variants^42^) further supported its potential contribution to DD pathogenesis (**Supplementary Table 8**).

Other residues with recurrent DNMs in DD patients, despite lacking prior disease annotations, exhibited amino acid changes that were largely absent from general populations (**Supplementary Table 8**) and generally low HistMTR values, implicating potential pathogenicity. Many of these sites are located in histone families other than H4. For the H2A family, G67 (HistMTR = 0.582) and G105 (HistMTR = 0.260) carried two and three DNMs in DD patients, respectively. These five mutations gave rise to five distinct H2A variants (**Fig. 4c**), four of which are absent from the RGC-ME database despite each being theoretically generated by 13-15 possible SNVs. For H2B, Y40 (HistMTR = 0.377) and T119 (HistMTR = 0.660) each harbored two DNMs in DD patients, resulting in three distinct H2B variants (**Fig. 4c**). All three were undetected in RGC-ME, though they could arise from 15-17 potential SNVs in the human genome.

Although one recurrent mutation site was identified in H3 (**Fig. 4c**), its presence in the RGC-ME database and high HistMTR value (0.784) suggest that this site is relatively tolerant of missense changes. Consequently, this variant is less likely to represent a high-impact pathogenic mutation. However, several H3 variants observed as DNMs in DD patients—such as H3R49S and H3S57P—were absent from RGC-ME, implicating their involvement in DD. Together, these findings broaden the spectrum of pathogenic variation linked to DD and underscore that non-H4 RDHs represent a substantial and previously underrecognized source of disease-associated variation.

For the ASD cohort, we assessed the contribution of core RDH genes using rare variant counts stratified by MPC pathogenicity bins from Fu et al.^51^. HistMTR stratification was not performed, as the genomic coordinates of these variants were unavailable. Analysis of highly deleterious variants (MPC > 2, ‘MisA’) revealed strong enrichment in both TADA-defined ASD-related genes^51^ (OR = 1.97, *P* = 5.3 × 10^-13^) and marginally significant enrichment in core RDH genes (OR = 1.90, *P* = 0.061; **Fig. 4d**). Notably, in the intermediate pathogenicity bin (1 < MPC ≤ 2, ‘MisB’), core RDH genes showed the strongest enrichment (OR = 1.81, *P* = 0.010), exceeding ASD-related genes. The enhanced signal for core RDH genes in the intermediate MPC category likely reflects the generally lower MPC estimates characteristic of RDH gene mutations (**Supplementary Fig. 17b**). Collectively, these findings support the contribution of RDH gene mutations to the genetic architecture of ASD.

To evaluate the contribution of core RDH gene variants to other human disease-related phenotypes, we performed rare variant association tests with the UKB cohort. Previous work found that genes under strong selective constraint are significantly associated with reduced reproductive success, a relationship that is probably mediated by genetically associated cognitive and behavioral traits^52^. Therefore, we first conducted association tests for reproductive and cognitive phenotypes (Methods). Associations were tested for three classes of core RDH gene variants: putative pathogenic missense, putative loss-of-function (pLoF), and synonymous variants (Methods).

We examined three reproductive phenotypes—spontaneous abortion, male infertility, and female infertility—and found no significant associations with core RDH coding variants (**Supplementary Table 9**). This lack of association may be due to the limited number of affected cases in UKB and consequently low statistical power. In contrast, analyses of cognitive phenotypes revealed a significant association between the core RDH missense burden and fluid intelligence scores (**Fig. 4e**; **Supplementary Table 9**). A nominally significant association was also observed with reaction time (*P* = 0.009, uncorrected; **Supplementary Table 9**). These signals were absent when all missense variants were analyzed without HistMTR filtering (**Supplementary Fig. 22**), indicating improved specificity. These results suggest that, despite the ‘healthy volunteer’ bias inherent to the UKB cohort^53–55^, rare deleterious RDH variants may influence cognitive performance within a generally healthy population, potentially through effects on neurodevelopmental functions.

To explore broader disease associations, we performed a phenome-wide association study (PheWAS) of core RDH missense variants. No phenotype achieved the significance level after Bonferroni correction (**Supplementary Table 10**). We attribute this null result to two plausible factors. First, as described above, the underrepresentation of severe early-onset conditions in UKB limits power to detect highly penetrant mutations^53–55^. Second, HistMTR does not discriminate the pathogenicity of alternative amino acid substitutions at the same residue (**Supplementary Table 3**); as a result, some benign missense variants at low HistMTR sites might be misclassified as putatively pathogenic, introducing noise that could attenuate association signals.

### Roles of regulatory mutations of RDH genes in human disease

Unlike most protein-coding genes, RDH genes in metazoans produce mRNAs that terminate in a conserved SLBP-binding stem-loop structure rather than a poly(A) tail (**Fig. 5a**). This stem-loop element, along with the adjacent histone downstream element (HDE), facilitates cleavage via recruitment of the U7 small nuclear ribonucleoprotein (snRNP). Because of essential regulatory roles, these stem-loop sequences themselves exhibit strong conservation across vertebrate species (**Fig. 5b**). RDH stem-loops showed significantly higher rare SNV density compared to other 3′UTR regions in the human genome (*P* = 1.6 × 10^-5^, *t*-test; **Fig. 5b**), indicating they are also mutation hotspots in the genome.

**Fig. 5:**
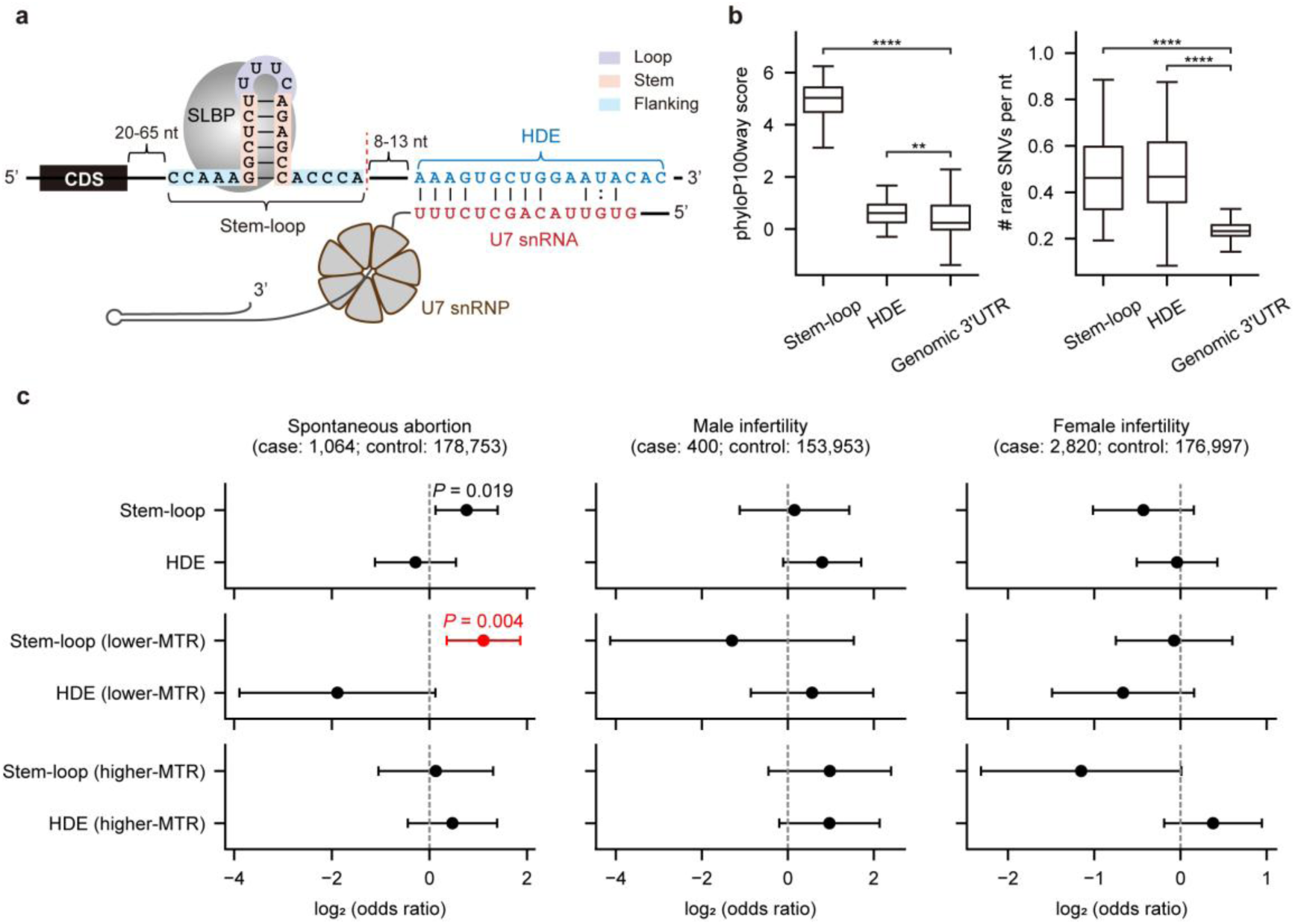
Conserved regulatory elements in RDH genes show elevated mutability and contribute to disease. **a,** Schematic of the important regulatory elements in 3′ regions of human RDH pre-mRNAs, highlighting the conserved stem-loop and HDE regions. The stem-loop is bound by the SLBP, a critical regulator of histone metabolism. The HDE, positioned 8–13 bp further downstream, binds the U7 snRNP and is required for 3′-end processing. **b,** Evolutionary conservation and mutability across regulatory elements. Left: phyloP100way conservation scores for stem-loops, HDEs and genomic 3′ UTR regions. Right: rare SNV density for the same loci. *P* values were obtained by two-sided Mann–Whitney U tests (*: 0.01 < *P* ≤ 0.05; **: 0.001 < *P* ≤ 0.01; ***: 0.0001 < *P* ≤ 0.001; ****: *P* ≤ 0.0001). **c,** Burden associations of rare regulatory variants in RDH genes with reproductive phenotypes. Odds ratios and 95% confidence intervals are shown. Bars are colored red if the association remained significant after Bonferroni correction (adjusted *P* < 0.05). Raw *P* values < 0.05 are shown.

To assess the functional impact of these variants, we performed a rare variant burden analysis after excluding weakly conserved segments—specifically, the flanking regions of stem-loops and unpaired sites within HDEs (Methods; **Supplementary Fig. 23**). We observed a negative correlation between RDH gene expression (germline and somatic) and gene-level MTR, indicating that highly expressed genes are under stronger constraint (**Supplementary Fig. 24**). We therefore hypothesized that regulatory variants in RDH genes with stronger constraint (also higher expression) would be more deleterious, and stratified genes by MTR to re-run the association analysis (Methods).

An initial analysis considering all RDH genes revealed a nominally significant association between stem-loop variant burden and spontaneous abortion (*P* = 0.02). After stratifying RDH genes by gene-level MTR, the burden of rare stem-loop variants in lower-MTR genes showed a stronger association with spontaneous abortion (OR = 2.15, 95% CI: 1.28-3.63, *P* = 0.024 after Bonferroni correction; **Fig. 5c**), whereas variants in higher-MTR RDH genes showed no signal. This suggests that regulatory mutations in lower-MTR RDH genes may specifically impair female fecundity or early embryonic viability. We found no significant association of rare stem-loop variants with clinical infertility (**Fig. 5c**).

No significant associations were detected for rare HDE variants across the tested reproductive phenotypes (**Fig. 5c**; **Supplementary Table 11**), possibly reflecting a more subtle functional impact of mutations in this element. However, among higher-frequency variants, several HDE variants showed nominally significant associations with infertility prior to multiple-testing correction (**Supplementary Fig. 25**; **Supplementary Table 12**), implicating their effects on gametogenesis or fertilization. Interestingly, one HDE variant in *H2BC5* was associated with both better cognitive performance and poorer reproductive outcomes (**Supplementary Fig. 25**), a pattern consistent with antagonistic pleiotropy.

By performing PheWAS and controlling for linkage disequilibrium (LD) (Methods), we identified multiple significant associations between RDH regulatory variants and diverse disease categories (**Supplementary Fig. 26**; **Supplementary Table 13**), such as hyperthyroidism and systemic lupus erythematosus, further implicating a role for non-coding variation in RDH genes in human disease.

### Strong ongoing negative selection acting on core RDH genes

RDH genes exhibited extreme long-term constraint, reflected in exceptional sequence conservation and elevated phyloP^56^, phastCons^57^ and GERP^58^ scores (**Supplementary Fig. 6**). Beyond these deep-time signals, and given links to developmental and reproductive disorders, it is important to determine when deleterious RDH variants are purged during the life cycle and the strength of ongoing purifying selection in humans. To address the former, we organized our findings according to Endler’s classification of non-sexual selection^59^, distinguishing fecundity, gamete, and viability selection (**Fig. 6a**); to address the latter, we inferred the distribution of fitness effects (DFE) of RDH gene variants.

**Fig. 6:**
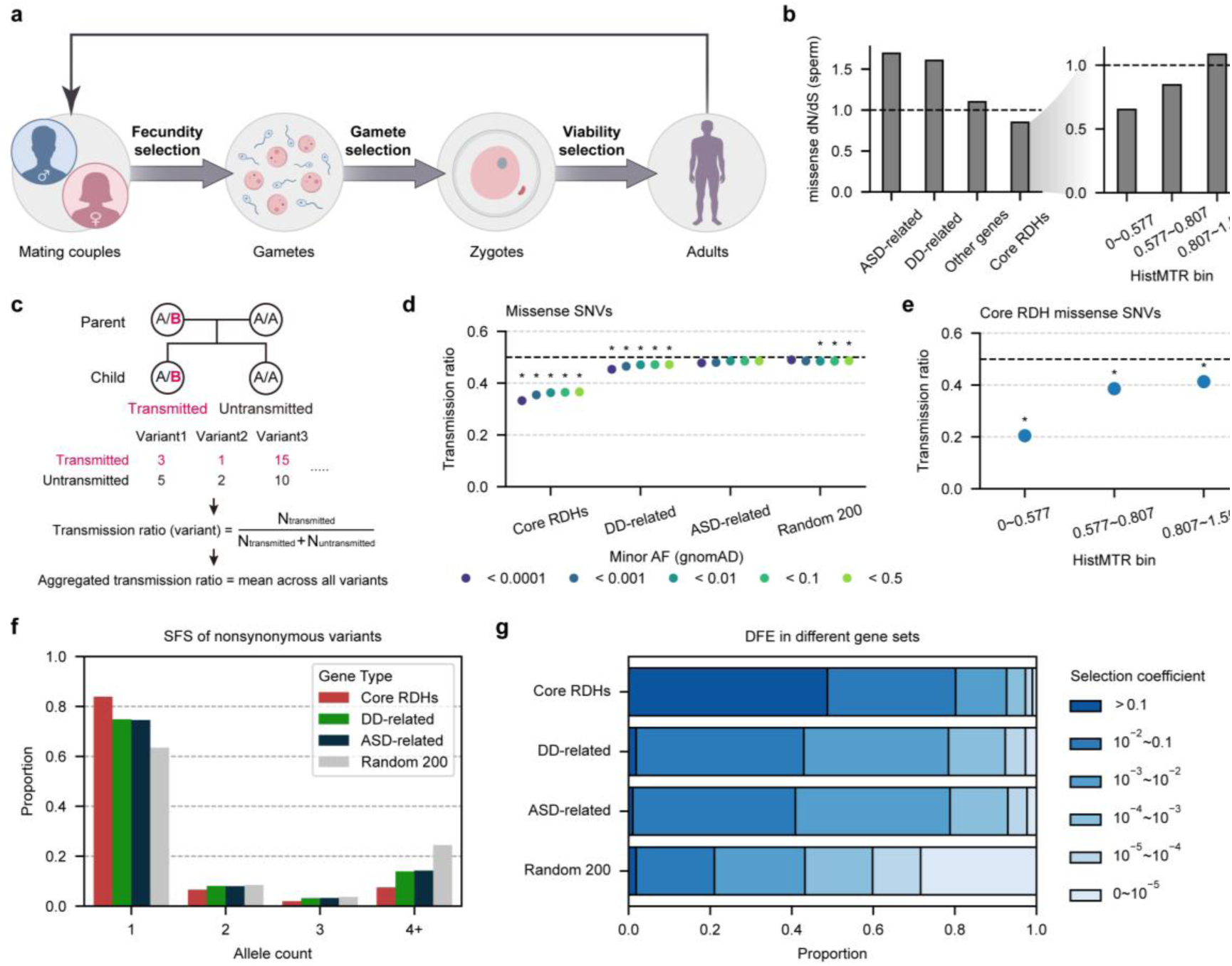
Strong ongoing selection on deleterious variants in RDH genes. **a,** Schematic presentation of Endler’s classification of non-sexual selection. Fecundity selection pertains to the production rate of functional gametes and viable zygotes derived from mating couples. Gamete selection acts on gametes before and during fertilization. Viability selection operates throughout the development from zygotes to reproductive adults. Graphical elements were adapted from BioGDP^60^. **b**, Comparison of dN/dS ratios of coding mutations in sperm sequencing data among gene groups. For core RDH genes, missense mutations were further stratified into three bins by HistMTR, and dN/dS was calculated for each bin. **c,** Method for calculating transmission ratios of derived alleles for a given gene group. **d,** Transmission ratios of missense variants across gnomAD AF bins for different gene groups. The thick dashed line (ratio = 0.5) indicates the neutral expectation. Asterisks denote significance after Benjamini–Hochberg correction (*P* < 0.05) based on simulation analysis (Methods). **e,** Transmission ratios of missense variants across HistMTR bins for core RDH genes. **f,** Site frequency spectra of nonsynonymous variants after dadi downsampling (Methods). Variants with allele count ≥ 4 were merged into ‘4+’. **g,** Estimated DFEs of nonsynonymous variants for different gene categories. Proportions of variants are shown for different selection coefficient (|s|) bins.

Fecundity selection acts through variation in reproductive success, including the ability to produce functional gametes and viable zygotes. Evidence for such selection on RDH genes emerged from their association with infertility phenotypes, described in the previous section. Rare damaging RDH gene variants appeared to be enriched in individuals with infertility or spontaneous abortion, suggesting that these variants reduce reproductive success. Because fertility-related traits directly influence the number of offspring produced, deleterious RDH gene variants affecting gonad development, germ cell development or other reproductive functions are likely to be eliminated by selection before contributing substantially to the next generation. These disease associations therefore pointed to fecundity selection as an important component of purifying selection acting on RDH genes.

Selection may also act at the level of gametes themselves, favoring functionally competent sperm or oocytes. Two independent lines of evidence supported strong gamete-level selection against deleterious RDH gene variants, particularly in the male germline. First, analysis of coding DNMs identified from recently published sperm sequencing data^7^ revealed a markedly reduced missense dN/dS ratio (0.848) for RDH genes compared with other genes (dN/dS = 1.100) (**Fig. 6b**, left). Stratification of RDH sites by HistMTR further showed progressively lower dN/dS values in lower-HistMTR bins (**Fig. 6b**, right). This depletion of missense RDH DNMs in sperm suggests that many new deleterious mutations are immediately purged during spermatogenesis. Second, transmission ratio distortion (TRD) analysis in parent–offspring trio sequencing data provided complementary evidence (**Fig. 6c**). Analysis with 599 trios in 1000 Genomes (1KG) project revealed that missense variants in RDH genes had lower transmission ratios than those in other gene groups and were significantly under-transmitted relative to the Mendelian expectation of 0.5 (**Fig. 6d**; **Supplementary Table 14**). Stratification by HistMTR further showed lower transmission ratios for missense variants in lower- HistMTR bins (**Fig. 6e**), whereas synonymous variants showed no such distortion (**Supplementary Fig. 27**). Similar TRD patterns were also observed in two independent trio exome sequencing datasets derived from 948 youngsters and fetuses, indicating the robustness of the results (**Supplementary Fig. 28**). The weaker TRD patterns observed in the two exome datasets likely reflected a shorter duration of exposure to selection and/or ascertainment bias toward individuals with disease phenotypes. Moreover, essential regulatory elements in RDH genes also exhibited lower transmission ratios than 3′ UTRs in other genes (**Supplementary Fig. 29**; **Supplementary Table 15**). The TRD patterns could result partly from selective depletion of deleterious alleles during gamete formation or fertilization. Together, reduced dN/dS in sperm and under-transmission of missense variants in trios implicated gamete selection as an important selective force shaping standing variation in RDH genes.

Viability selection acts after zygote formation, through effects on embryonic development, postnatal survival, or functional performance. Multiple results from earlier sections pointed to this mode of selection operating on RDH genes. Rare damaging RDH missense variants were significantly enriched in DD and ASD cohorts, and burden analyses in the UK Biobank further demonstrated the association between RDH missense variants and reduced fluid intelligence. Moreover, multiple mutations have been reported to cause Mendelian genetic diseases, especially neurodevelopmental disorders^18^. These findings indicate that RDH gene variants can impair developmental and cognitive functions, reducing overall fitness even in individuals who survive to adulthood. Transmission ratio analysis also supported viability selection. Under-transmission of RDH missense variants could partly reflect reduced survival of embryos or fetuses carrying these alleles, in addition to gamete-level selection. Using the UK Biobank data, we found that missense pN/pS estimates stratified by age remained nearly constant across adult age bins ranging from 39 to 72 years (**Supplementary Fig. 30**). This lack of age dependence in relatively old adults suggests that most viability selection occurs before birth or early in life.

To synthesize these stage-specific signals, we further quantified the DFE for coding variants in RDH genes. The site frequency spectrum of nonsynonymous variants in RDH genes was heavily skewed toward ultra-rare variants, with the fraction of singletons substantially exceeding those of DD-related genes, ASD-related genes, or randomly selected genes (**Fig. 6f**). Correspondingly, DFE inference indicated that 80.2% of nonsynonymous variants in RDH genes falled into the strongly deleterious class (selection coefficient *s* ≥ 0.01)—roughly double the fraction observed for DD- or ASD-related genes and nearly fourfold higher than genomic background (**Fig. 6g**; **Supplementary Table 16**). Based on the DFE, we estimated that although core RDH genes comprise only 0.8‰ of the total coding region, they contribute 6.5‰ (95% CI: 4.1‰-9.8‰) of all newly arising strongly deleterious nonsynonymous variants genome-wide. More strikingly, ∼49% of nonsynonymous variants in RDH genes exhibited a selection coefficient of ≥ 0.1, compared with fewer than 2% in other gene groups (**Fig. 6g**). Given such strong selection, majority of newly arising nonsynonymous variants in RDH genes may stay in a population for fewer than five generations^61^. Similar trends were observed in both the UKB and 1KG datasets (**Supplementary Fig. 31**; **Supplementary Table 16**).

These results demonstrated that RDH genes experienced some of the strongest ongoing purifying selection in the human genome. Integrating evidence across fecundity, gamete, and viability selection revealed a consistent picture: although RDH genes are hypermutable, deleterious coding variants are eliminated at multiple stages of the life cycle, preventing their accumulation in human populations.

## Discussion

Through the in-depth comparative analysis, we revealed that RDH genes, whose protein sequences are among the most evolutionarily conserved, harbor a markedly elevated burden of germline mutation in vertebrates. This elevated mutation burden necessitates cautious interpretation of variation in these loci. As many mutations in H4 genes were observed in gnomAD, previous work has suggested that H4 genes are tolerant to both loss-of-function and missense variation^37^, but our findings indicated that this perceived tolerance more likely reflects their intrinsically high mutation rate rather than genuine genetic resilience, as evidenced by the very low population frequencies of most variants in histone genes.

Regarding the underlying mutational mechanisms, our analyses implicated AID/APOBEC cytidine deaminases as a major contributor to the elevated mutability of RDH genes. Functional genomic data from human and mouse revealed clear AID/APOBEC activity in RDH genes, likely due to off-target effects. Although these datasets were not derived directly from germline cells—a limitation that warrants caution—several lines of evidence support the biological plausibility of germline activity. APOBEC enzymes are known to be expressed to restrict retrotransposons^62^ and are proved to generate heritable genomic mutations^27^. While direct evidence of AID-mediated germline mutagenesis remains unreported, AID is expressed in germ cells and has been implicated in epigenetic regulation^63,64^, with demonstrated mutagenic potential in non-B cell lineages^65^. Furthermore, our mutational signature analysis in population genomic data provided additional support for AID/APOBEC activity in the germline. Future studies are needed to determine the precise stage(s) of germline development at which elevated mutagenesis in RDH genes occurs.

The high mutation rates observed in RDH genes likely reflect an evolutionary trade-off following the emergence of AID/APOBEC enzymes^27^, which confer enhanced immune functions in vertebrates. Although the elevated mutation burden imposed by AID/APOBEC activity on RDH genes incurs a fitness cost, the immune benefits provided by these enzymes are likely much more indispensable. It is plausible that off-target mutagenesis of histone genes was even more pronounced early in vertebrate evolution, with subsequent selection acting to partially mitigate this burden. One potential mitigating strategy may be the evolutionary transition from tandemly repeated histone arrays, common in invertebrates, to more dispersed and non-repetitive RDH gene organization in many vertebrates (e.g. birds and mammals), which could limit sustained exposure to AID/APOBEC activity. However, directly testing this evolutionary scenario remains challenging.

The pronounced mutability of RDH genes necessitates accurate quantification for variant deleteriousness in these genes. When evaluated against existing variant effect predictors, even the top-performing tools showed relatively poor performance for RDH gene mutations. To address this, we developed HistMTR, a novel missense constraint metric that integrates evolutionary information across paralogs within the same histone family. HistMTR outperformed existing predictors and identified novel candidate variants associated with disease risk, demonstrating its high potential for prioritizing pathogenic variants in clinical settings. These results underscore the value of analyzing functionally related paralogs collectively rather than in isolation, substantially enhancing the power to uncover pathogenic variants in multicopy gene families. We note that, due to few or no observed variants for certain amino acid substitutions in RDH genes, HistMTR currently assigns the same score to all substitutions at a given residue; incorporating data from a larger cohort in the future could improve the resolution of deleteriousness predictions. In addition, our use of synonymous variants as a neutral baseline may introduce bias, given emerging evidence that some synonymous mutations are subject to selection^66^.

By integrating HistMTR with large-scale genomic and phenotypic data, we identified associations between RDH genes and multiple human diseases, particularly developmental disorders. Our analysis extended beyond coding regions to include conserved non-coding regulatory elements of RDH genes, revealing that these regulatory sequences also contribute to disease risk. Given the emerging roles of both coding and non-coding variants of RDH genes in developmental disorders, spontaneous abortions and other diseases, incorporating these potentially highly deleterious variants in prenatal genetic screening and preimplantation genetic testing for monogenic disorders (PGT-M) in assisted reproductive technologies may be warranted. Despite the prevalence of infertility, its genetic architecture remains poorly defined; few loci have achieved genome-wide significance due to high genetic heterogeneity, modest individual effect sizes, and substantial phenotypic complexity^67^. This limits standard GWAS discovery, making rare coding variants (like those in RDH genes) and non-GWAS methods important for understanding the genetic architecture of infertility.

The multicopy nature of RDH genes implies a dose-dependent threshold for clinical pathogenesis^68^. Unlike single-copy genes where a 50% reduction typically triggers haploinsufficiency, the structural redundancy of identical histone paralogs likely buffers the genome against individual variants. However, missense mutations—particularly those exerting dominant-negative effects^17^—can exceed a critical stoichiometric threshold to ‘poison’ the global nucleosome pool. Moreover, for the same damaging RDH mutation, the chromosomal positions of affected nucleosomes are likely stochastic across individuals, potentially leading to substantial phenotypic heterogeneity and incomplete penetrance. Consequently, deciphering the functional translation of specific RDH variants into diverse clinical outcomes remains a significant challenge. Recent modeling suggested that over 1,000 developmental disorder-associated genes remain undescribed, often characterized by lower penetrance^49^. The RDH genes—with their unique combination of high mutational influx and multicopy buffering—likely constitute a critical component of this previously hidden genetic landscape, requiring integrated mechanistic and population-scale inquiry to fully resolve.

Our evolutionary analyses uncovered distinct selection patterns in core RDH genes. First, we found that loss-of-function variants in core RDH genes are relatively more tolerated than missense variants—a pattern opposite to that observed in most other genes. This may reflect their multicopy genomic organization, wherein reducing the dosage of functional histone proteins may be less deleterious than incorporating structurally aberrant variants into chromatin. Second, sperm mutation data and transmission ratio distortion analyses indicate that core RDH missense variants are selectively filtered out during gametogenesis, embryonic development, or early life stages. Combined with the inferred DFE with population genomic data, these results demonstrate that RDH mutations contribute disproportionately to the overall mutational load and are subject to exceptionally strong purifying selection in human populations.

Together, we uncovered an unexpected vulnerability of ultraconserved histone genes and elucidated the evolutionary mechanisms and functional consequences of their elevated mutation burden. The paralog-aware constraint metric HistMTR enabled more accurate prioritization of pathogenic variants in highly conserved, multicopy gene families. Our analytical approach is broadly applicable to other important multicopy gene families in the human genome, thereby accelerating functional and disease-related studies of these complex regions.

## Methods

### Data preparation

#### Representative transcript selection

For humans and macaques, representative transcripts for each gene were selected based on the ‘Ensembl_canonical’ tag in the GTF files from GENCODE^69^. For other species, the transcript with the longest coding sequence was chosen. For mouse RDH genes, we manually selected transcripts lacking introns. Only autosomal protein-coding genes were retained for downstream analyses. Annotation sources for each species are provided in **Supplementary Table 17**.

#### Extraction of 4-fold degenerate sites

4-fold degenerate sites were defined from representative transcripts. Here, only 4-fold degenerate sites without conflict across transcripts were considered. Sites that were 4-fold degenerate in one transcript but not in another were excluded to preclude potential effects of selection.

#### Identification of RDH and RIH genes

For humans and mice, RDH and RIH genes were annotated following the definitions of Seal et al.^70^. For other species, we employed a three-step procedure:

1. Protein sequences for all genes, extracted from the GTF annotation, were searched against human histone proteins using NCBI BLASTp (v2.16.0)^71^. Genes with e-value < 1 × 10^−5^ and sequence similarity > 50% were retained as putative histone genes.
2. Histone stem-loop structures were identified using Infernal (v1.1.4) cmscan^72^ with parameters ‘--cut_ga --rfam --nohmmonly’.
3. A putative histone gene was classified as RDH gene if other histone genes were located within ± 100 kb of its locus and a stem-loop was detected within the gene body or within 70 bp downstream of the 3′ end; genes not meeting both criteria were designated RIH gene.

In mice, this procedure missed only one RDH gene (*H2bc1*) due to the absence of a detectable RDH-specific stem-loop, and no RIH gene was misclassified as RDH gene. Four additional mouse genes (*H2bu2*, *H2aw*, *H4bc24* and *H4f16*) met RDH gene criteria but appear to be previously unreported RDH genes not recorded by Seal et al. In humans, no RIH gene was misclassified, and five replication-dependent H4 genes (*H4C2*, *H4C7*, *H4C12*, *H4C14* and *H4C15*) were missed owing to the absence of a cmscan-detectable stem-loop (identification of stem-loops to be described below). Above results demonstrated the high recall and low false positive rate of this procedure for RDH gene identification.

#### Identification of regulatory stem-loops and HDEs in 3′ UTRs of human RDH genes

Stem-loop structures of histone genes were initially identified in 65 loci using cmscan from Infernal (v1.1.5)^72^ to search for the Rfam motif RF00032. These hits were aligned with MAFFT (v7.525)^73^ and manually refined based on highly conserved sites. For RDH genes lacking a detectable stem-loop, we extracted the 3′ UTR plus 100 bp of downstream sequence and aligned it to the consensus motif. This procedure enabled stem-loop annotation in five additional H4 genes. Genomic coordinates of all stem-loops of RDH genes (hg38) are provided in **Supplementary Table 18**.

To identify HDEs, we extracted each stem-loop sequence along with 36-bp downstream sequence as candidate regions. Using nucleotides 3-18 of the human U7 snRNA gene (GTGTTACAGCTCTTTT; obtained from Rfam) as the query, we predicted RNA–RNA interactions with IntaRNA (v3.4.1)^74^. Predictions were restricted to candidate positions 30–55 (‘--tRegion 30–55’) of input candidate sequences. We evaluated seed lengths of 3–10 bp (‘--seedBP’) while permitting 0–3 unpaired bases within the seed (‘--seedMaxUP’). For each candidate, the interaction with the lowest predicted free energy was selected as the most likely HDE site. Downstream regions were subsequently classified as HDE or non-HDE; HDE regions were further subdivided into unpaired and paired categories based on their base-pairing pattern with U7 snRNA. Full annotation results are provided in **Supplementary Table 19**.

#### Excluded genomic regions

To minimize potential biases in downstream analyses, we excluded genomic regions with low mappability or abnormal sequencing coverage. Genome mappability for each species was calculated using GenMap (v1.3.0)^75^ with parameters ‘-K 150 -E 1’. Per-base coverage of human genome was obtained from gnomAD; for other species, coverage was estimated from a set of randomly selected sequenced individuals using deepTools bamCoverage (v3.5.6)^76^. Regions with mappability < 1 or coverage outside 0.5-1.5× the mean sequencing depth were removed from variant density calculation and other related analyses.

#### Gene expression data

For germline expression, raw RNA-seq data of human fetal testis (n = 24) and ovaries (n = 24) were obtained from Lecluze et al.^77^ and processed uniformly. Reads were adapter- and quality-trimmed with Trim Galore (v0.6.10, https://github.com/FelixKrueger/TrimGalore). Transcript abundance was quantified with Salmon (v1.10.2)^78^ in quant mode while enabling sequence- and GC-bias correction (‘--seqBias’ and ‘--gcBias’). Gene-level estimates were produced by supplying a transcript-gene map to Salmon (‘-g’). Gene-level counts were then combined across samples and filtered with edgeR (v4.0.16)^79^ ‘filterByExpr’ using default settings; genes retained by this procedure were designated ‘expressed genes’ and the remainder ‘unexpressed genes’.

For somatic expression, RNA-seq experiments were retrieved from the ENCODE portal^80^ using the following filters: released experiments; human samples; tissue-derived biosamples; total RNA-seq assays; non-perturbed conditions; embryonic or child life stages; and exclusion of experiments flagged for low replicate concordance. For biosamples sharing the same term name, one sample was randomly selected. Expression matrices were downloaded directly from ENCODE. Owing to annotation version differences, several RDH genes were remapped as follows: {H3C2, ENSG00000274267; H3C3, ENSG00000278272; H2AC18, ENSG00000203812; H2AC19, ENSG00000272196}.

### Variant processing

#### Population variant data

For wild mouse (*Mus musculus*), barn owl (*Tyto alba*), cunner (*Tautogolabrus adspersus*), and ninespine stickleback (*Pungitius pungitius*), no suitable, precomputed population-variant callsets were available. We therefore compiled published population whole-genome sequencing (WGS) data for these species and performed variant calling and filtering pipeline (detailed metadata provided in the **Supplementary Table 17**).

Raw data were downloaded and processed with fastp (v0.24.0)^81^ to remove adapter sequences and trim low-quality bases, producing clean reads for downstream analyses. Reference genomes for each species are listed in the **Supplementary Table 17**. Clean reads were aligned to the corresponding reference genome using BWA-MEM2 (v2.2.1)^82^. PCR duplicates were marked using MarkDuplicates in GATK (v4.6.1.0)^83^, and variants were called per sample using GATK HaplotypeCaller in GVCF mode to generate single-sample gVCF files. Base-wise depths across the whole reference genome and individual-level average sequencing depth metrics were subsequently computed using mosdepth (v0.3.10)^84^ with default parameters. The sequence alignment and variant detection workflows were integrated into an automated nf-core/sarek pipeline (v3.7.0)^85^ with parameters fine-tuned (--max_cpus, --max_memory), thereby maximizing the utilization of parallel computing resources. Per-sample gVCFs were subsequently jointly genotyped and merged using GATK to produce a multi-sample VCF for each species.

Cryptic relatedness can reduce power to detect rare variants. After filtering and pruning variants based on LD, we estimated within-species pairwise related score using Hail’s pc_relate module^86^. Pairs with related score > 0.05 were considered related. We then used Hail’s maximal_independent_set module to release a maximal dataset with completely unrelated individuals. For variant sites, we performed hard filtering using GATK4 VariantFiltration on SNPs and INDELs independently, excluding SNPs with ‘QUAL<30, QD<2, FS>60, MQ<40, MQRankSum<-12.5, Read PosRankSum<-8 or SOR>3’. To ensure accurate estimation of allele frequency (AF), genotypes with genotype quality (GQ) < 20 were masked as ‘./.’ and excluded from AF calculations. Rare variants were defined as those with 1 × 10^−4^ in humans and 1 × 10^−2^ in other species, because of the smaller sample sizes available for the latter.

For human, macaque (*Macaca mulatta*), chicken (*Gallus gallus*), amphioxus (*Branchiostoma japonicum*), honey bee (*Apis mellifera*) and *Arabidopsis thaliana*, we directly used publicly available, downloaded SNP datasets (**Supplementary Table 17**). These datasets were processed using the same individual-level and site-level filtering strategy described above to ensure comparability across species.

#### Human de novo mutation data

DNMs from 9,652 trios were obtained from Palsson et al.^87^. We restricted analyses to 618,192 SNVs and used these variants to estimate the genome-wide mutation spectrum and *de novo* mutation rates across different gene groups. For each gene group, the total number of callable nucleotides was calculated as the sum of filtered region lengths across genes, multiplied by the number of trios and by two to account for diploidy. The mutation rate (per site per generation) was then estimated as the number of DNMs divided by the total callable nucleotide count.

Sperm mutations were obtained from Neville et al.^7^. Germline mutations identified by deep targeted sequencing and exome NanoSeq were annotated using ANNOVAR^88^ based on GENCODE v44. Coding mutations (n = 35,186) were used for gene-group dN/dS calculations, as described in the section ‘Calculation of group-level pN/pS’.

#### Human somatic mutation data

Somatic mutations detected from normal tissues by WGS or whole-exome sequencing (WES) were obtained from SomaMutDB^89^. We included samples generated by the ‘Clone’ (n = 5,319) and ‘LCM’ (Laser Capture Microdissection, n = 4,467) methods. Mutations were first aggregated at the tissue level, and recurrent mutations from the same donor were deduplicated. After merging all tissues, 7,091,705 SNVs used to calculate exon-level mutation burden for each gene.

Somatic mutations from tumor tissues were obtained from The Cancer Genome Atlas (TCGA, https://www.cancer.gov/tcga, <u>2026-02-23</u>). We analyzed open-access TCGA mutation annotation format (MAF) files derived from WES of tumor samples. Recurrent mutations within the same patient were removed using the same preprocessing strategy, yielding 3,654,067 SNVs for downstream analyses. Exon-level mutation burden was subsequently estimated.

### Mutability analysis

#### Ortholog inference and dN/dS calculation

Orthologous relationships between human and mouse genes were inferred using OrthoFinder (v3.0.1b1)^90^ with protein sequences from all species as input. For non-RDH genes, one-to-one orthologs were taken directly from OrthoFinder results. RDH orthologs were identified based on standardized histone gene nomenclature^70^, which is conserved across mammalian species. Protein sequence alignments were performed using ParaAT (v2.0)^91^ with MAFFT specified as the aligner (‘-m mafft’). dN/dS was calculated using KaKs_Calculator (v2.0)^92^ with the Yang-Nielsen model (‘-m YN’). Orthologs with dN > 2, dS > 5, dS < 0.01 or dN/dS > 10 were discarded, as these results might be due to incorrect orthology inference or inaccurate estimation.

#### Variant density calculation

To quantify the mutation rate at the gene level, we calculated the raw variant density as the number of rare variants divided by the effective gene length (after coverage and mappability filters). However, this raw metric is biased for relative comparisons because saturation at highly mutable sites leads to an underestimation of the true mutation rate. To mitigate this bias, we implemented a two-step correction procedure.

First, we applied mutation masking. Variants that were not classified as rare (i.e., those exceeding the allele frequency threshold or failing quality control filters) were excluded from the analysis. These sites were removed from both the numerator and the denominator of the density calculation, as their evolutionary status could not be reliably ascertained.

Second, we performed a Poisson-based correction to account for the possibility of recurrent mutation at the same site, which becomes non-negligible with high mutability. For each mutation type stratum *i* (e.g., a specific 3-mer context such as A[A>T]A), we defined:

*L*_effective,*i*_ as the effective number of sites after genomic region filtering.
*N*_mask,*i*_ as the count of sites additionally masked in the previous step.
*m*_*i*_ = *L*_effective,*i*_ − *N*_mask,*i*_ as the final number of eligible sites.

We modeled the mutation count per site as a Poisson process with the rate *λ*_*i*_. The probability that a site is mutated at least once is therefore 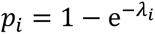. Given the observed number of sites with one or more rare variants, *N*_rare,*i*_, and assuming a binomial sampling model *N*_rare,*i*_ ∼ Binomial(*m*_*i*_, *p*_*i*_), the maximum likelihood estimator for the corrected mutation rate is derived in closed form:

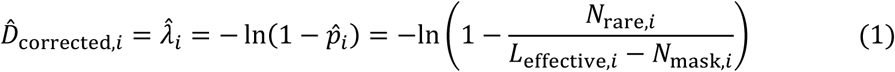

When a single summary statistic was required, we computed the site-weighted mean corrected density across all strata: *D̂*_corrected_ = (∑_*i*_ *L*_effective,*i*_*D̂*_corrected,*i*_)/ (∑_*i*_ *L*_effective,*i*_). For human-specific analyses, these calculations were also performed at the strand-aware 3-mer level. For cross-species comparisons of RDH genes versus other genes at 4-fold degenerate sites, we aggregated contexts into 1-mer strata (comprising the six basic substitution types and three CpG-specific types) to avoid issues arising from data scarcity in certain 3-mer contexts. The validity of this correction is supported by a substantial increase in the cosine similarity between the spectrum of rare variants and the spectrum of DNMs, indicating a lower bias in the corrected density estimate (**Supplementary Fig. 10**).

#### Transcriptional strand asymmetry score calculation

Transcription could introduce strand asymmetry in DNA mutagenesis. The non-transcribed strand, maintained in a single-stranded state during elongation, is more exposed to mutagens like deamination^14^. Conversely, the transcribed strand is the preferential target for transcription-coupled nucleotide excision repair (TC-NER)^93^. The interplay of these processes results in a characteristic mutational strand bias across the genome. To quantify this phenomenon across different gene groups, we calculated a mutational asymmetry score.

Prior to analysis, we excluded transcribed regions where genes are located on both strands to prevent confounding signals. For each gene group, variants were stratified based on the transcript direction. Using the corrected variant densities (*D*) described in the previous section, we computed strand asymmetry scores for the six 1-mer mutation types (C>T, C>A, C>G, T>A, T>C and T>G).

For a given mutation type in gene group *g*, the asymmetry score was defined as the log2 ratio of the variant density on non-transcribed strand (NTS) to the density on transcribed strand (TS). For example, the C>T asymmetry score was calculated as:

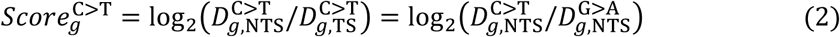

#### Mutation signature analysis

Mutational processes driven by specific etiological factors imprint characteristic patterns, or signatures, across the genome. We leveraged known single-base substitution (SBS) signatures from the COSMIC catalog^94^ to deconvolute the contributions of various mutagenic processes in RDH genes. Prior to signature decomposition, we computed the 3-mer frequencies for all usable genomic regions to establish a genome-wide background. The mutational catalog of each gene group was normalized to this background composition to account for base composition biases. To more accurately capture APOBEC-driven cytosine deamination, which occurs preferentially on single-stranded DNA, we replaced the standard SBS2 signature with its single-strand-specific counterpart, SBS2ss*, as defined by Liu et al.^95^ through single-molecule profiling.

To correct for potential saturation effects from highly mutable sites, we applied the same Poisson-based adjustment detailed in the ‘Variant density calculation’ section to the mutational counts before signature refitting. Deconvolution was performed using MuSiCal (v1.0.0)^31^, which employs a likelihood-based sparse non-negative least squares algorithm. We set a likelihood threshold of 0.005 to ensure robust signature assignment and limit false-positive identifications. Signatures were grouped etiologically based on COSMIC annotations: SBS2ss* and SBS13 were classified as APOBEC-associated; SBS9, SBS84 and SBS85 reflected AID activity. To statistically evaluate the enrichment of these specific processes, we performed a permutation test: 1) first, the clock-like signatures SBS1 and SBS5 were held constant as a universal baseline; 2) we then constructed a null distribution by iteratively refitting the spectra with five randomly selected COSMIC signatures, excluding all APOBEC- and AID-associated signatures; and 3) the empirical *P* value for the observed cosine similarity of a specific category of processes (e.g., AID/APOBEC) was derived from this null distribution.

Since proportion of variants with a CpG context were still slightly underestimated after correction, we conducted mutation spectrum analysis on a downsampled gnomAD v3.1.2 (HGDP+1KG) cohort of 1,500 unrelated individuals and used variants with AF < 5 × 10^-3^ in transcribed regions. To ensure the robustness of results, we also evaluated the following alternative processing strategies:

1. Singleton removal to mitigate potential contamination by somatic artefacts^96^.
2. UTR analysis to circumvent selection bias inherent to coding sequences.
3. TSS-focused analysis using a truncated 500-bp region downstream of the TSS to minimize signal dilution from long gene bodies, with balanced sampling of expressed and unexpressed genes to control for region length.
4. Analysis without downsampling and using variants with AF < 10^-4^ (i.e., consistent with the previous mutability analysis in humans).
5. Independent cohort validation using a downsampled UKB dataset of 1,500 unrelated White British individuals.

As detailed in **Supplementary Table 1**, permutation tests across all these strategies consistently demonstrated a significant improvement on refitting brought by AID/APOBEC-related signatures in RDH genes, confirming the robustness of our findings.

### ChIP-seq data processing

Multiple published ChIP-seq datasets were used to investigate AID/APOBEC activities at RDH gene regions. Raw reads of ChIP-seq data were quality-trimmed using Trim Galore (v0.6.10) and aligned to the reference genome with Bowtie2 (v2.5.3)^97^. PCR duplicates were marked and filtered using Sambamba (v1.0.0)^98^ with ‘[XS] == null and not unmapped and not duplicate’. Coverage tracks were generated using deepTools bamCoverage (v3.5.6)^76^. Peak calling was performed using MACS3 (v3.0.1)^99^. To generate comparative signal tracks, log10 *P* value and fold-change bedGraph files were generated using MACS3 bdgcmp. For direct visualization of enrichment, log2 ratio tracks of treatment over control were created using deepTools bigwigCompare (v3.5.6)^76^. These tracks were normalized using the SES method (‘--scaleFactorsMethod SES’), which provides robust library size adjustment based on background read distribution. A complete list of data sources and detailed software parameters are provided in **Supplementary Table 20**.

### Developing the HistMTR metric for assessing missense variant deleteriousness

Existing variant effect predictors exhibit limited predictive performance for RDH variants (**Fig. 3b**), likely reflecting challenges posed by the structural complexity of these genes (for structure-based models), extreme inter-species conservation (for sequence-based models), and short gene length (for variation-based models). Although some prior predictors have incorporated population-level genetic variation, none have leveraged information across multiple paralogous gene copies to enhance prediction accuracy. To address this gap, we aggregated variation across paralogs within each core RDH family to derive an improved metric, HistMTR, for assessing missense variant deleteriousness in RDH genes. The HistMTR framework extends the concept of the missense tolerance ratio (MTR)—a metric analogous to pN/pS but more robust when variant counts are low—and tailors it to the multicopy histone loci.

First, variants from the RGC-ME cohort were functionally annotated with ANNOVAR. Observed missense and synonymous variant counts (*O*_mis_ and *O*_syn_) were calculated for each gene, and expected counts (*E*_mis_ and *E*_syn_) were estimated by summing mutation rates predicted by MuRaL, which models single-nucleotide mutation rates using 2-kb local sequence context. Predicted mutation rates were lifted over from GRCh37 to GRCh38 using CrossMap (v0.7.0)^100^. Gene-level MTR and pN/pS values for RDH gene *g* were then calculated as:

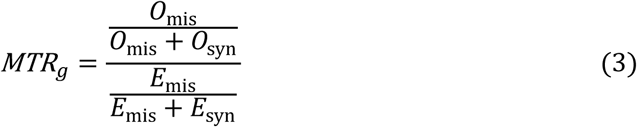

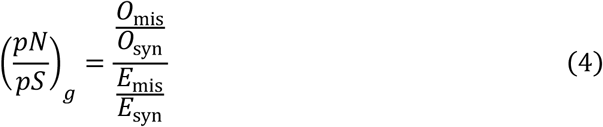

Next, multiple protein sequence alignments of paralogs within each core RDH family were aligned with MAFFT, and a family consensus was defined at each alignment column as the most frequent amino acid (gaps ignored). For each aligned site, observed and expected variant counts were aggregated across paralogs using pN/pS-based weights to obtain per-site counts of missense and synonymous variants. H2BC1 and H4C7 were excluded from analysis owing to <90% pairwise protein identity with other family members. For site *i* in family *f*, we defined an pN/pS-weighted paralog-based site-level MTR as:

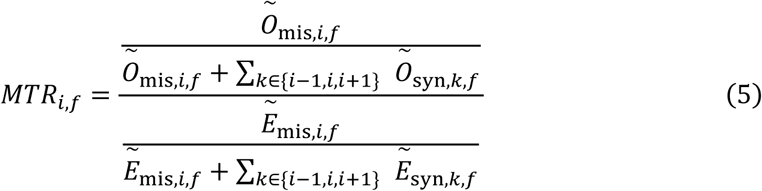

where the pN/pS-weighted observed and expected counts are defined as:

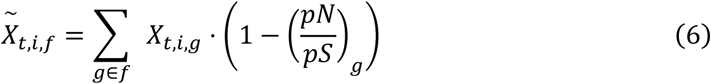

where *X* ∈ {O, E} denotes observed or expected counts, and *t* ∈ {mis, syn} indicates missense or synonymous variants. Since observed synonymous variants of individual codons are sparse and certain codons (e.g., codons of Met) lack synonymous changes, we used single-site missense counts but pooled synonymous counts over the (*i* − 1, *i*, *i* + 1) window; for terminal sites, the window used the next two internal positions.

Finally, site-level and gene-level MTR values were combined to obtain the HistMTR value for site *i* in a specific RDH gene *g* of family *f*:

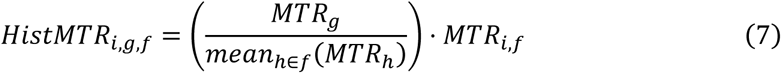

The rationale is that a given amino acid position in an RDH with a lower gene-level MTR should exhibit a correspondingly lower site-specific HistMTR. From the above equation, it can be inferred that the mean HistMTR across all paralogs at a given amino acid site equals the corresponding site-level MTR.

### Calculation of group-level pN/pS

For group-level pN/pS, we pooled counts across genes to estimate group-level constraint. For a gene set *g* and a variant class *c* ∈ {missense, nonsense}, the group-level ratio is

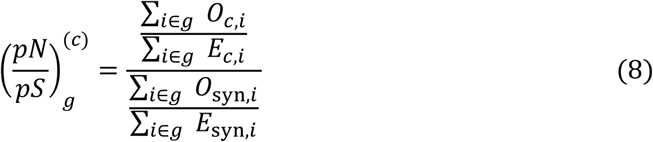

Here, *O*_*c*,*i*_ and *O*_syn,*i*_ are the observed numbers of class-*c* and synonymous variants in gene *i*; *E*_*c*,*i*_ and *E*_syn,*i*_ are the corresponding expected counts obtained by summing context-dependent per-site mutation probabilities derived from MuRaL over coding positions where that class is possible. For AF-stratified or age-stratified analyses, the same formula was applied within each bin.

Group-level dN/dS values were calculated analogously using the same formulation, except that observed mutation (or substitution) counts were used instead of variant (or polymorphism) counts.

### Nucleosome structure and histone modifications

Since many deposited nucleosome structures lack core histone N-terminal tails and/or the linker histone H1 which may lead to an incomplete histone interaction map, we generated a complete nucleosome model with AlphaFold3^101^ (server mode; random seed = 42). The histone octamer was specified by family consensus sequences; the DNA and H1 sequences were taken from PDB 7PFV^102^. The top-ranked prediction was used for subsequent interaction analyses and ChimeraX^103^ visualization.

Interatomic contacts were computed with PDBe Arpeggio^104^. We applied the following calling rules: (i) atom pairs from different chains within 3.9 Å were labeled as van der Waals contacts; (ii) pairs’ contacts annotated by Arpeggio contained ‘polar’ or ‘hbond’ were classified as hydrogen bonds; and (iii) pairs’ contacts annotated contain ‘ionic’ were classified as salt bridges. Atom pairs of set (ii) and (iii) were a subset of (i), and an atom pair could have multiple interaction types simultaneously.

α-helix and β-sheet annotations were extracted from the 7PFV mmCIF file; for each histone family, regions on different chains were merged to define final annotations. Additional secondary structure features were derived from figure 1e in Armeev et al.^105^. Residue exposure states were determined by calculating the relative solvent accessibility (RSA) using FreeSASA^106^, with maximum solvent-accessible surface area (SASA) values from Tien et al.^107^. A residue was classified as ‘exposed’ if its RSA exceeded a threshold of 0.3 in any of the nucleosomal chains.

Candidate PTMs for RDHs were retrieved from PTMcosmos (https://ptmcosmos.wustl.edu/) using human RDH gene symbols. PTMs were grouped into six classes (phosphorylation, acetylation, methylation, ubiquitination, glycosylation, and sumoylation) and for each core RDH family, sites were collapsed to consensus positions using the same approach applied to variants during HistMTR calculation.

### Phenotypic consequence of core RDH mutations

To evaluate the functional impact of histone mutations, we leveraged a humanized yeast library described by Bagert et al.^12^, in which native yeast histones were fully replaced with human histones harboring specific variants. This system was repurposed to quantify the effect of mutations on cell viability. From the original dataset of 130 amino acid mutations, we analyzed the 102 mutations that could be caused by a single human SNV. These variants were categorized into three phenotypic classes: ‘no phenotype’, ‘impaired growth’, and ‘no growth’. For subsequent binary classification, ‘no phenotype’ was treated as a negative label, while ‘impaired growth’ and ‘no growth’ were combined as a positive label.

We benchmarked the predictive performance of the HistMTR metric against seven other variant effect prediction scores on core RDH genes. AlphaMissense and REVEL scores were annotated using Ensembl VEP (release 110)^108^. The data of RGC-MTR^42^ and MPC^46^ were obtained from public sources and integrated via in-house scripts. For a given amino acid change, the family-level consensus score for each of the above metric was calculated as the unweighted mean of scores of all possible substitutions; a mutation-rate weighted mean was also tested but yielded inferior performance. ESM scores^44^ represent the averaged negative log-likelihood ratios of the mutated versus reference amino acid from five ESM-1v models, using the family consensus sequence as input. ProSST scores^47^, which incorporate structural context, were generated by first predicting the monomeric structure of each core histone consensus sequence using the AlphaFold3 server; the top-ranked result for each sequence was then used as input. Like ESM, ProSST assigns each mutation a negative log-likelihood ratio comparing the mutated and reference amino acid. Attempts to use a whole nucleosome complex (without DNA) as input for ProSST resulted in poorer performance, likely because the model was primarily trained on monomeric structures. A final set of 99 mutations, for which all eight scores were available, was used for benchmarking. Performance was assessed by AUROC and AUPRC.

### Analyzing variant data from DD and ASD cohorts

45,221 DNMs from 31,058 DD patients were obtained from the DeNovoWEST repository (input files accompanying Kaplanis et al.^49^). Records were converted to the VCF format, lifted over to GRCh38 using CrossMap, and annotated using ANNOVAR. From the 285 DD-related genes reported by DeNovoWEST, we filtered to retain protein-coding genes with canonical transcripts on autosomes (GENCODE V44), leaving 249 DD-related genes for the final comparison. In the absence of public large-scale unaffected trio-sequencing datasets, population variants from non-Finnish European (NFE) subset of gnomAD (34,029 individuals, comparable in scale to the DD cohort) were used as controls. ORs, 95% CIs, and *P* values were calculated using two-sided Fisher’s exact test implemented in R ‘fisher.test()’.

Rare coding variants from ASD case and control cohorts were extracted from the supplementary table of Fu et al.^51^. Like that in Fu et al., missense variants were stratified by MPC score into three tiers: MisB (MPC ≥ 2), MisA (1 ≤ MPC < 2), and other missense (MPC < 1), and were counted per gene. Because their data used another histone nomenclature style, we defined RDH genes as genes whose gene symbol begins with ‘HIST’; entries containing ‘H1’ in the gene symbol were further filtered to define the core RDH gene subset (n = 45). For comparison, 72 ASD-related genes defined by the TADA framework^51^ were grouped as a category. Enrichment of missense variants in cases versus controls for each MPC tier was assessed using two-sided Fisher’s exact tests, reporting ORs, 95% CIs, and *P* values for each gene group.

### Association analysis

#### Phenotype definition

We defined six phenotypes for association testing, categorized into two groups: three related to cognitive ability and three to reproductive capacity. The phenotypic data for analysis were from UK Biobank.

### Cognitive phenotypes

Fluid intelligence score (UK Biobank Data-Field 20016), qualifications (Data-Field 6138) and reaction time (Data-Field 20023) were obtained from the initial assessment visit. Reaction time values were inverse-normal transformed prior to analysis. The highest qualification reported for each individual was mapped to the International Standard Classification for Education (ISCED) coding for years of education as described by Fridman et al.^52^.

### Reproductive phenotypes

Spontaneous abortion was defined by ICD-10 code O03. Male infertility and female infertility were defined based on a combination of ICD-9, ICD-10, Read Codes, and self-reported records, as previously described by Venkatesh et al.^67^.

#### Rare variant burden calculation

Coding variants in core RDH genes were classified as rare if the cohort allele frequency ≤ 0.1%. Missense variants were considered putatively pathogenic if the HistMTR score was ≤ 0.6303 This threshold corresponded to the upper bound of the first two bins defined by HistMTR (that is, the lowest 40% of sites), in which missense DNMs from individuals with DD were enriched (**Fig. 4a**). Putative loss-of-function (pLoF) variants included those annotated by ANNOVAR as stop-gain, stop-loss, frameshift insertion, or frameshift deletion. Synonymous SNVs within coding regions served as a negative-control variant class in association models.

Regulatory variants located in the stem-loop or HDE regions of RDH genes were defined as rare if the cohort allele frequency ≤ 0.1%. In the absence of a dedicated pathogenicity predictor for non-coding variants, we stratified RDH genes by gene-level MTR to form two groups (lower-MTR versus higher-MTR). Lower-MTR RDH genes were defined as the top 50% of genes with the lowest gene-level MTR within each RDH family.

For each burden category (e.g., putatively pathogenic missense), individuals were coded as binary carriers (1, individuals carrying ≥ 1 rare variants; 0, non-carrier).

#### Association testing

Analyses were restricted to unrelated participants of White British ancestry. Binary traits were tested using logistic regression, and continuous traits using ordinary least-squares regression. All models were adjusted for age, age^2^, sex, and the first ten genetic principal components.

PheWAS was performed with PheTK^109^ using phecodeX mappings of ICD-10 codes. We applied the same covariates, required a minimum of 50 cases per phecode (‘--min_case 50’), set the minimum phecode count for case ascertainment to 1 (‘--min_phecode_count 1’), and corrected for multiple testing across phecodes using the Bonferroni method.

To minimize potential spurious associations driven by linkage disequilibrium (LD), all associations remaining significant after Bonferroni correction in PheWAS were subjected to further LD filtering. For each significant phecode string, the most relevant disease phenotypes and corresponding disease IDs were retrieved via the Open Targets^110^, prioritizing exact name matches or, if unavailable, the top five ranked related diseases. For these diseases, if the disease-associated genes reported by Open Targets are located within the histone gene cluster region (chr6: 23,000,000–30,000,000 in hg38) but are not RDH genes, the corresponding associations with RDH genes identified in our PheWAS analysis were considered potentially attributable to LD with these disease-associated genes. Given the high gene density and complex LD architecture in this region, no single LD threshold can reliably distinguish RDH-specific signals from those driven by neighboring disease-associated genes. We therefore conservatively excluded all such RDH-disease associations from downstream analyses.

### Transmission ratio distortion analysis

#### Data sources and preprocessing

Transmission analysis was performed using trio sequencing data from three sources. First, genomic data for 599 trios (1790 unique individuals) from the 1KG project were obtained from gnomAD v3.1.2 (HGDP+1KG subset). Second, we reanalyzed WES data for 420 trios (866 unique individuals) from an ASD-related cohort published by Sanders et al.^111^, with proband ages ranging from 4 to 18 years. The reads were aligned to the hg38 reference using BWA-MEM2 (v2.2.1), sorted with SAMtools (v1.21)^112^, and deduplicated using GATK (v4.6.1.0) MarkDuplicates. Variants were called with GATK HaplotypeCaller, followed by joint genotyping across samples using GATK CombineGVCFs and GenotypeGVCFs. Resulting SNVs were filtered using GATK SelectVariants and VariantFiltration with QUAL<30, QD<2, FS>60, MQ<40, MQRankSum<-12.5, Read PosRankSum<-8 or SOR>3. Third, whole-exome data from 528 trios (1584 unique individuals, unpublished) from the First Affiliated Hospital of Sun Yat-sen University (FAH-SYSU) were processed identically to the Sanders et al. dataset and included as an independent validation cohort. Most offspring samples (n = 479) of the FAH-SYSU cohort were derived from fetuses through prenatal genetic testing.

#### Transmission ratio calculation

Variant consequences (synonymous or missense) were annotated with ANNOVAR, and only variants in regions passing coverage and mappability filters were retained. Variants with a call rate < 95% were excluded. For each retained variant, we iterated over all trios and applied the following quality controls before counting transmission events:

1. Trios with any missing genotype in the child, father, or mother were excluded.
2. Genotypes with quality (GQ) < 20 or mapping depth (DP) < 20 were excluded.
3. For heterozygous genotypes, the allele balance (alt / [ref + alt]) within an individual was required to be between 0.2 and 0.8.
4. Only trios in which one parent was heterozygous and the other homozygous were considered informative for transmission analysis (**Fig. 6c**).

For each variant, we first calculated the sum of the reference and alternate allele frequencies in gnomAD, excluding sites where this sum was below 0.99 to avoid potential artifacts from multi-allelic loci. For retained variants, we analyzed the transmission of the minor allele based on gnomAD AF values. Within a specific trio dataset described above, the transmission ratio per variant was calculated as *N*_transmited_/(*N*_transmited_ + *N*_untransmited_). Group-level transmission ratios were obtained by averaging across all variants in that group.

To assess statistical significance, we simulated a null distribution by performing 1,000 iterations in which each transmission event was assigned with probability 0.5. The empirical *P* value was defined as the proportion of simulations in which the simulated transmission ratio fell below the observed ratio.

### Distribution of fitness effects analysis

#### Variant data used

For the gnomAD dataset, we focused on variants from the non-Finnish European (NFE) subset in gnomAD v4, selecting records using the INFO fields ‘AC_nfe’ and ‘AN_nfe’. For the UKB exome, we randomly selected 350 unrelated White British individuals from each age bin, yielding 2,100 individuals in total. For the 1KG, we included 493 unrelated individuals with super-population code ‘EUR’.

We retained only SNPs and applied coverage and mappability filters to define high-confidence regions. Coding consequences were annotated using ANNOVAR. Folded synonymous and nonsynonymous site frequency spectra (SFS) were generated for the whole coding sequence and for each gene group separately. To account for sample size differences and missing data, we projected the SFS from the full dataset to a standardized sample size using ‘dadi.Spectrum.from_data_dict’ implemented in dadi^113^ (**Supplementary Table 21**).

#### Inference of demographic model

We assumed a demographic model including an out-of-Africa bottleneck, subsequent recovery phase, and recent exponential growth (**Supplementary Fig. 32**). This model was fitted to the synonymous SFS of the whole coding sequence using ‘dadi.Inference.opt’. Following Kim et al.^114^, we modified the internal ‘timescale_factor’ parameter in dadi to 10^−6^ to improve numerical accuracy for growth-related parameters. For each dataset, we performed 100 random perturbations of initial parameter values using ‘dadi.Misc.perturb_params’ and retained the maximum-likelihood estimate (**Supplementary Table 21**).

#### Calculation of the ancestral population size

For the whole coding sequence, we estimated the synonymous population mutation parameter (*θ*_syn_) under the inferred demographic model using the synonymous SFS and ‘dadi.Inference.optimal_sfs_scaling’ in dadi. To obtain *θ*_nonsyn_, we assumed *θ*_nonsyn_/*θ*_syn_ = 2.5, consistent with the ratio estimated from MuRaL-predicted mutation rates (2.485). We assumed a human exome mutation rate (*μ*) of 1.38 × 10^−8^ per site per generation, as reported for healthy probands^115^ and consistent with our estimate from non-histone genes (1.374 × 10^−8^, **Supplementary Fig. 4**).

After applying coverage and mappability filters, the total coding sequence length (*L*) was 30,197,545 bp, with a synonymous sequence length of 8,627,870 bp (*L*/(1 + 2.5)). The ancestral population size (*N*_*a*_) was calculated from the relationship *θ* = 4*N*_*a*_ ∗ μ ∗ *L* (**Supplementary Table 21**).

#### Inference of distribution of fitness effects

We precomputed SFSs over a grid of scaled selection coefficients (γ = 2*N*_*a*_*s*) using ‘dadi.DFE.Cache1D’ with parameters ‘gamma_bounds=[1e-5, 500]’ and ‘gamma_pts=300’. We assumed a gamma DFE for *γ* (‘DFE.PDFs.gammà in dadi), parameterized by a shape parameter (*α*) and a scale parameter (*β*).

Inferences of the DFE require an estimate of *θ*_nonsyn_. For the whole coding sequence, *θ*_nonsyn_ was set to 2.5 ∗ *θ*_syn_. Because sequence length and mutation rate differ across gene groups, we estimated *θ*_syn_ separately for each group using its synonymous SFS. The expected *θ*_nonsyn_/*θ*_syn_ ratio for each group was derived from MuRaL-predicted mutation rates (core RDH: 2.427; DD-related: 2.487; ASD-related: 2.556; random 200 genes: 2.504), yielding group-specific *θ*_nonsyn_ estimates. These values and nonsynonymous SFSs were used as inputs for DFE inference with ‘dadi.Inference.opt’.

For each dataset and gene group, we performed 100 random perturbations of initial parameter values and retained the maximum-likelihood estimate (**Supplementary Table 16**). Based on the best-fitting model, *γ* was converted to *s* using the estimated *N*_*a*_, and the proportions of variants within predefined selection coefficient bins were calculated.

## Data availability

All data analysed are included in the main text and its supplementary materials of this study. All the analyses were based on published data except for the FAH-SYSU exome data. The FAH-SYSU exome data are available upon reasonable request, subject to protections for participant privacy.

## Code availability

The code for the performed analyses in this study is available on GitHub at https://github.com/JuseTiZ/RDH_mutation_analysis.

## Acknowledgements

We thank the members of the Li lab for helpful discussion throughout the project. This research has been conducted using the UK Biobank Resource under Application Number 54148. This study makes use of data generated by the DECIPHER community. A full list of centres who contributed to the generation of the data is available from https://deciphergenomics.org/about/stats and via email from. DECIPHER is hosted by EMBL-EBI and funding for the DECIPHER project was provided by the Wellcome Trust [grant number WT223718/Z/21/Z]. This work was supported by National Natural Science Foundation of China (32470690).

## Author contributions

C.L. conceived and supervised the project. Z.J., J.X., Y.T., M.L., S.L., and C.L. performed the analyses. Z.J., J.X., Y.T. and C.L. wrote the initial draft of the manuscript. All authors provided critical feedback and contributed to revising the manuscript.

## Supplementary Figures

**Supplementary Fig. 1:**
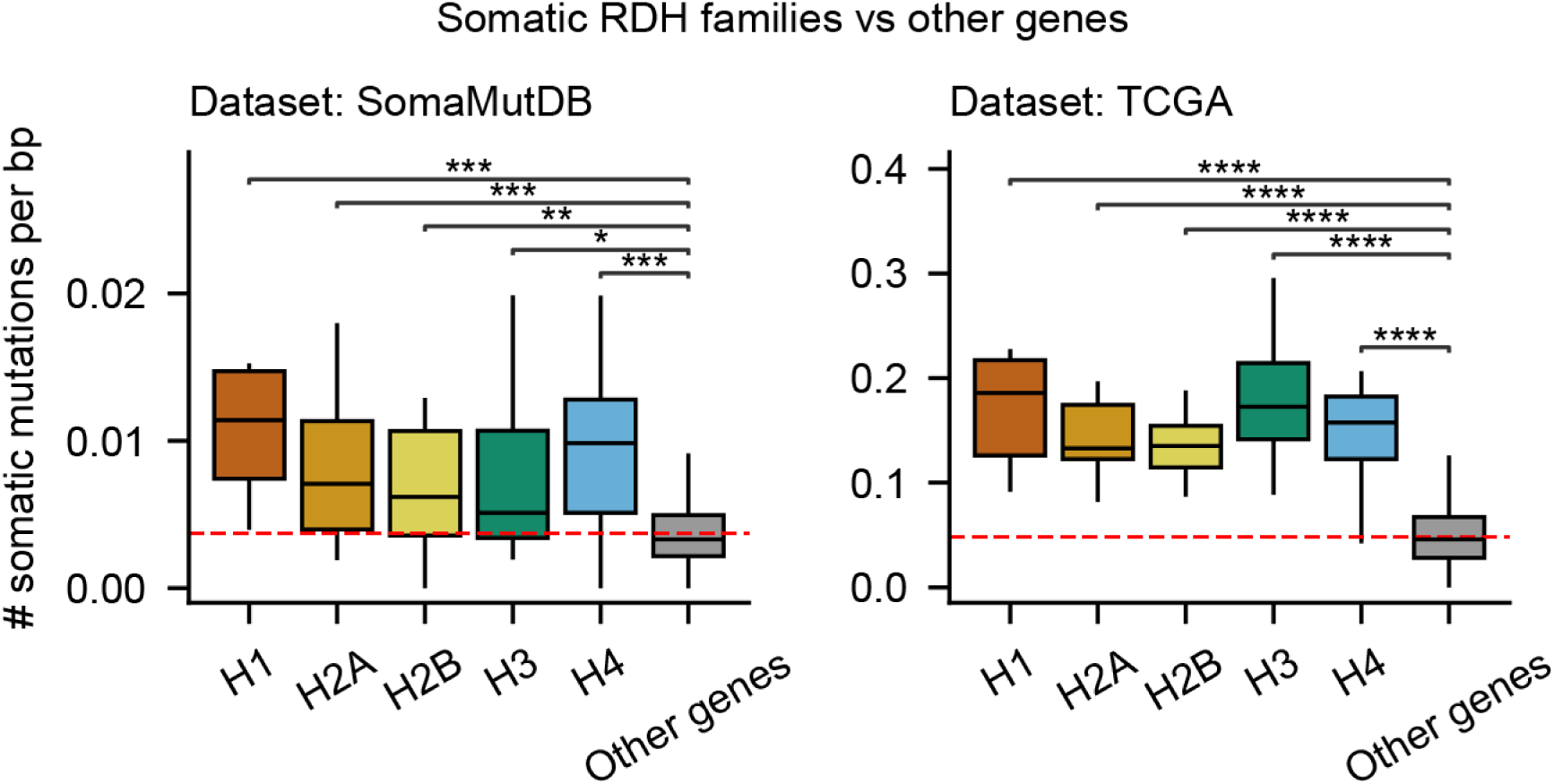
Elevated somatic mutability across different RDH gene families. Somatic mutation density comparisons in normal somatic tissues (from SomaMutDB) and primary tumors (from TCGA). The red lines indicate the mean somatic mutation density across all genes (SomaMutDB: 0.0037; TCGA: 0.0483). Benjamini-Hochberg corrected *P* values from two-sided Mann–Whitney U tests are shown; all comparisons were significant (*: 0.01 < *P* ≤ 0.05; **: 0.001 < *P* ≤ 0.01; ***: 0.0001 < *P* ≤ 0.001; ****: *P* ≤ 0.0001).

**Supplementary Fig. 2:**
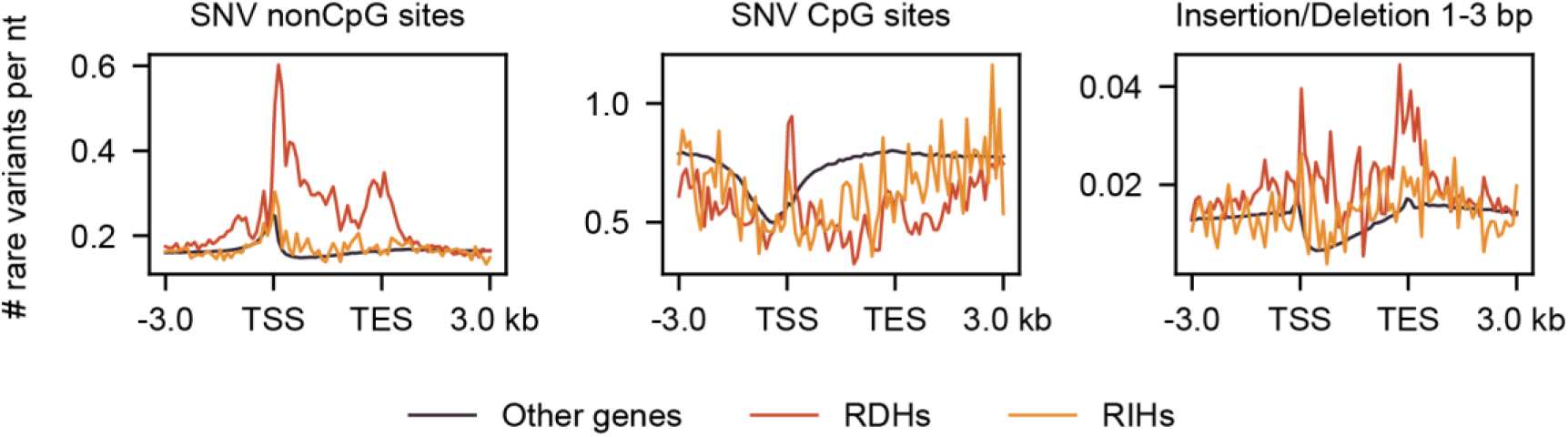
Comparison of rare variant density profiles across genic regions among different gene groups. Meta-gene profiles of rare variant densities in 100-bp bins around gene bodies (scaled to 3 kb) and 3 kb flanks. Lines depict mean rare variant densities for non-CpG SNVs, CpG SNVs, and short indels (1–3 bp) across RDH genes, RIH genes, and other coding genes in the human genome. TSS, transcription start site; TES, transcription end site.

**Supplementary Fig. 3:**
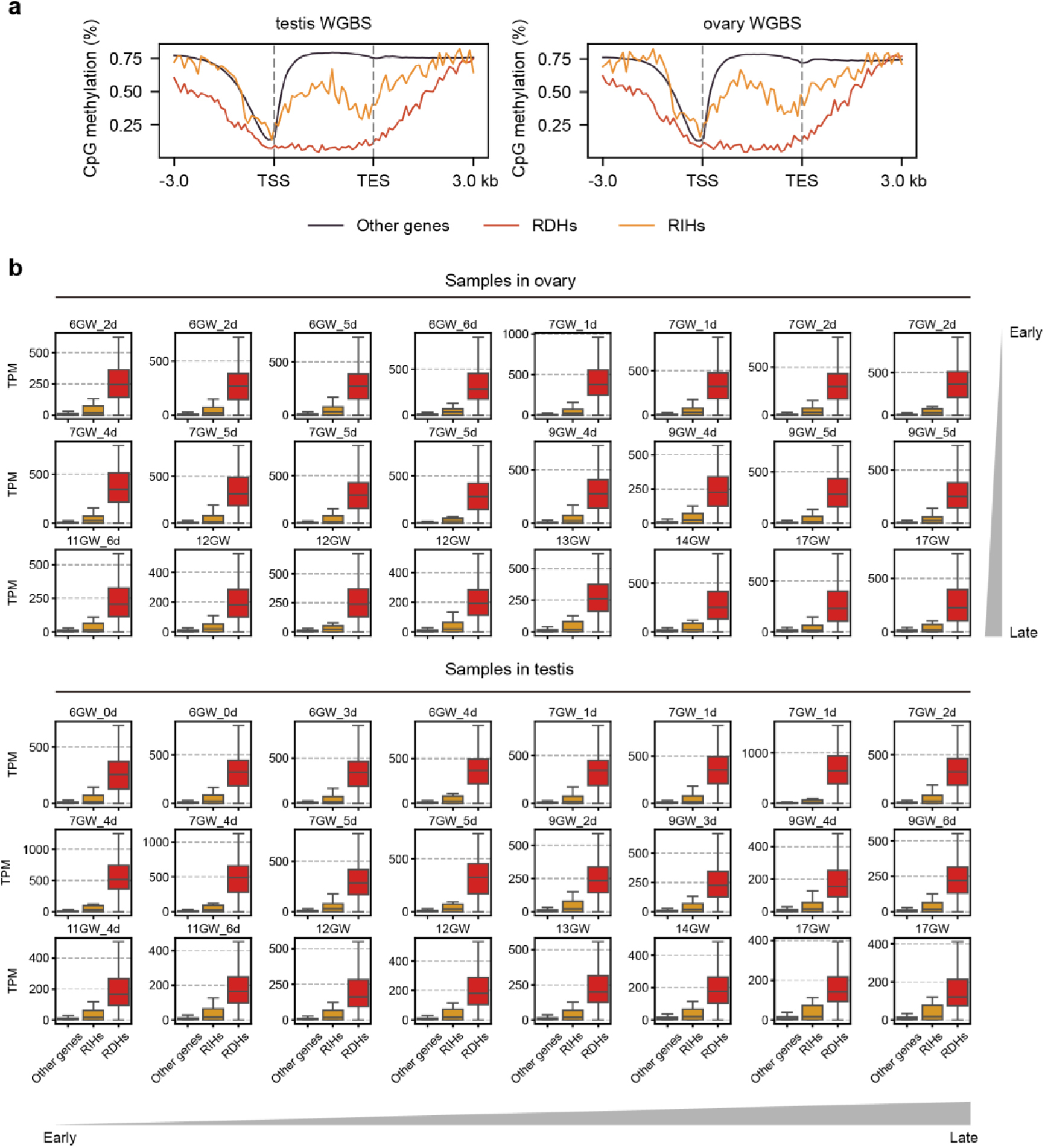
Methylation landscape and germline expression of histone genes. **a,** DNA methylation meta-gene profiles of RDH genes, RIH genes, and other genes in testis (left; ENCODE: ENCSR806NNG) and ovary (right; ENCODE: ENCSR803SIO). **b,** Germline expression levels of gene groups in human fetal ovary and testis, based on RNA-seq data from Lecluze et al.^77^. Samples are split by tissue (ovary or testis) and ordered chronologically from early to late gestational age. Expression (TPM, transcripts per million) is shown as box plots for other genes, RIH genes, and RDH genes.

**Supplementary Fig. 4:**
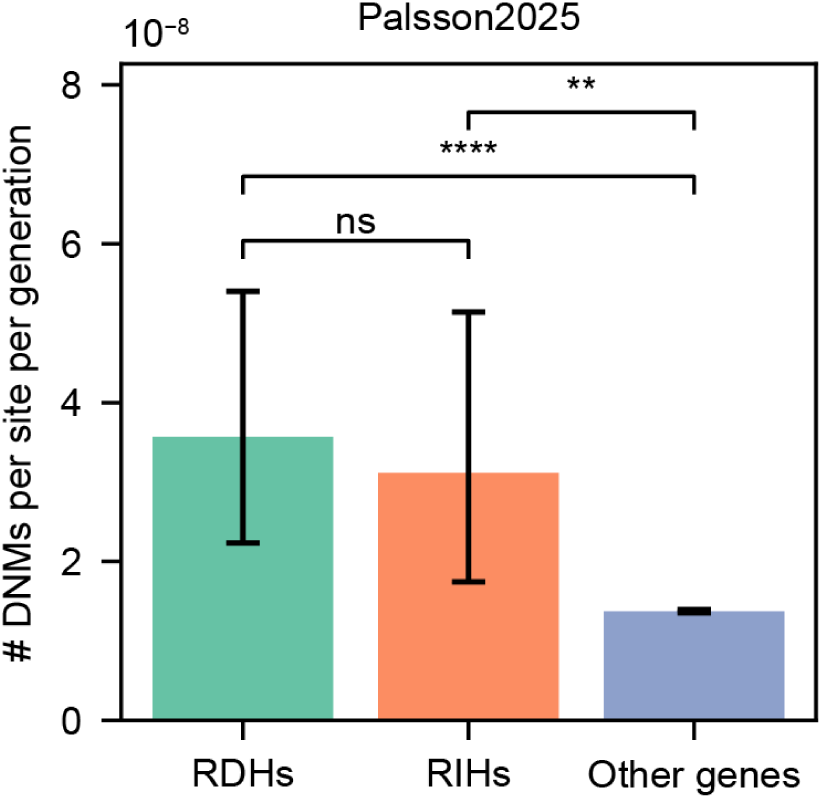
*De novo* mutation rates across different gene groups. *De novo* mutation rates of RDH genes, RIH genes and other genes. *De novo* mutation counts were modeled as Poisson-distributed events proportional to the genomic region length. Mutation rates were estimated as the number of observed mutations divided by callable bases. Exact 95% confidence intervals were calculated using the Garwood method for Poisson counts. Differences between regions were assessed using a two-sided score test for equality of independent Poisson rates. All *P* values are given above the bars (*: 0.01 < *P* ≤ 0.05; **: 0.001 < *P* ≤ 0.01; ***: 0.0001 < *P* ≤ 0.001; ****: *P* ≤ 0.0001).

**Supplementary Fig. 5:**
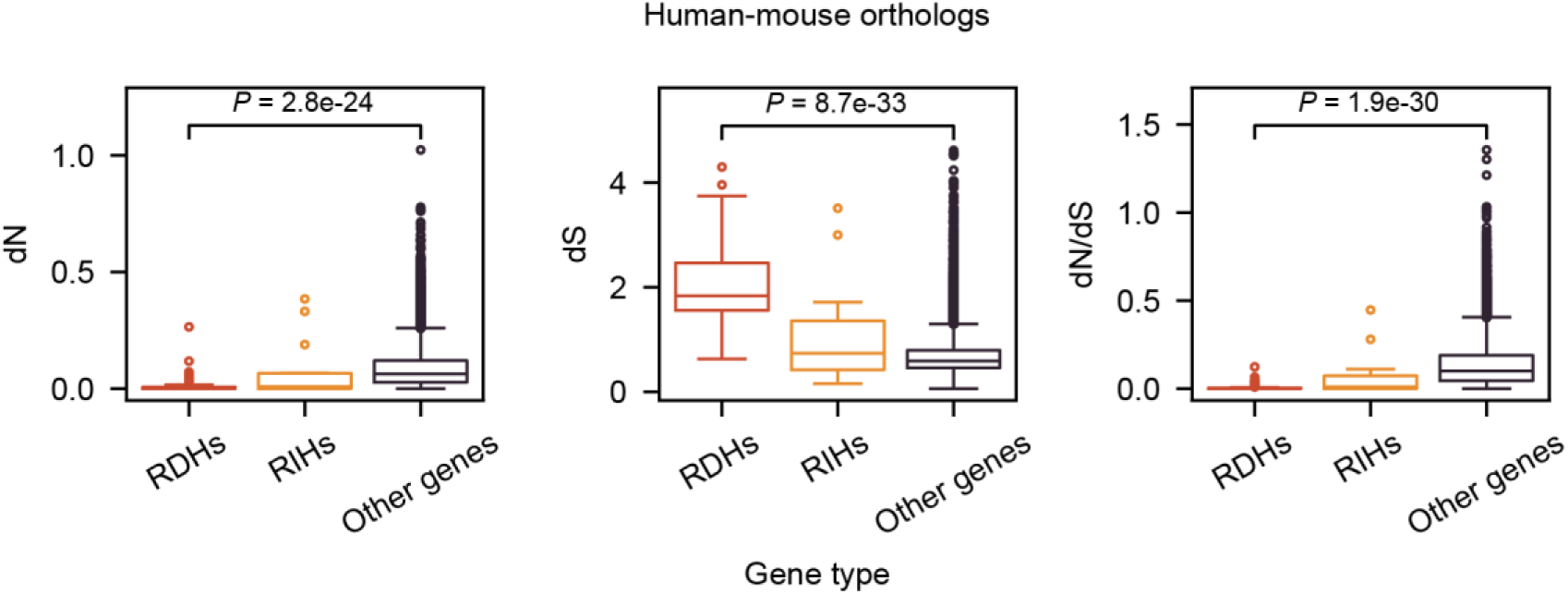
Evolutionary rates across gene groups for human-mouse orthologs. Distribution of nonsynonymous rate (dN, left), synonymous rate (dS, middle) and dN/dS ratio (right) across gene categories. RDH genes show significantly lower dN but higher dS values compared to other gene groups (two-sided Mann-Whitney U test). Higher dS in RDH genes reflects the elevated mutability during evolution. The low dN/dS reflects strong purifying selection in RDH genes.

**Supplementary Fig. 6:**
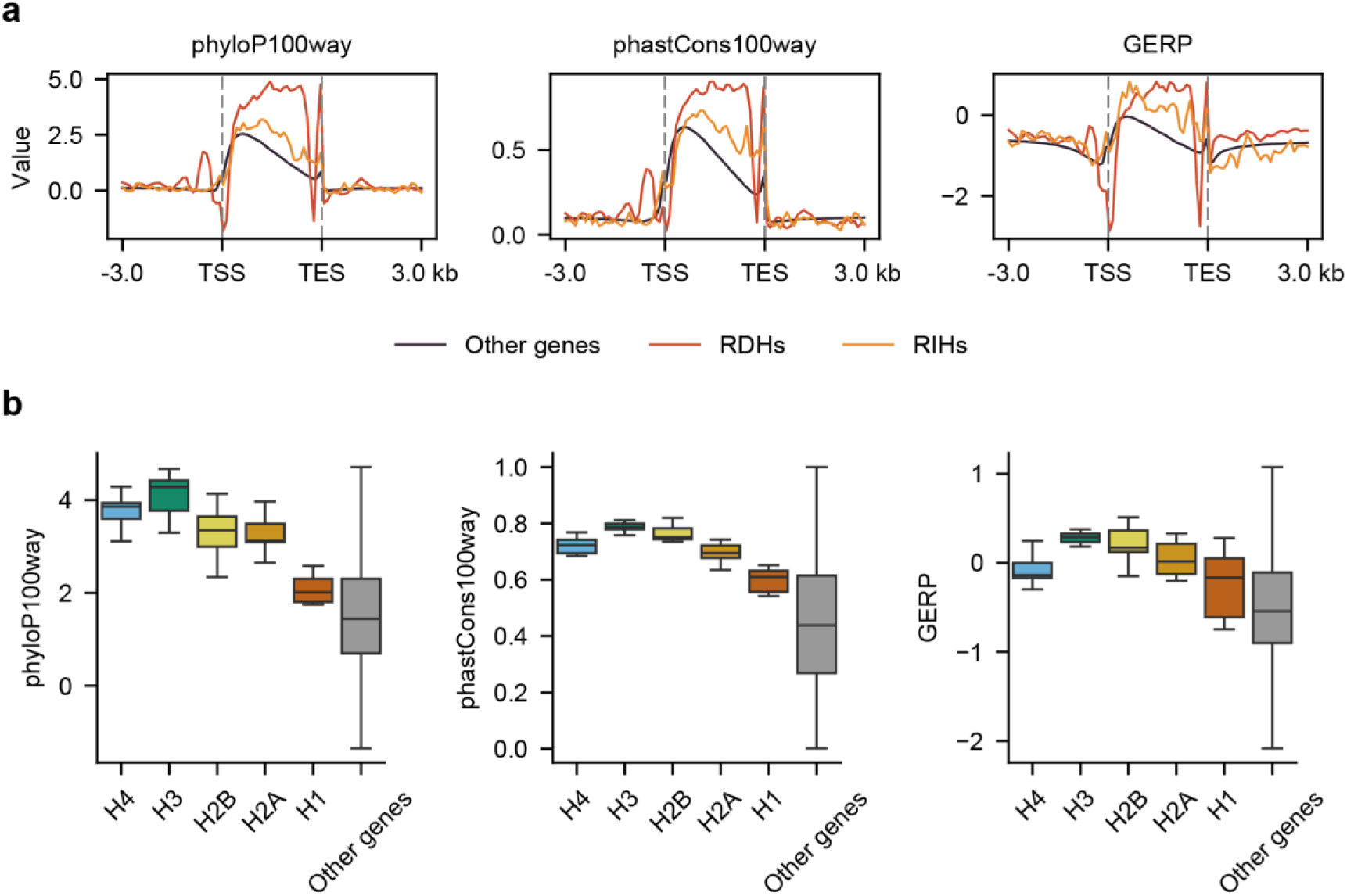
RDH genes are highly conserved across multiple evolutionary metrics. **a,** Conservation profiles of RDH genes, RIH genes, and other genes based on three metrics—phyloP100way, phastCons100way, and GERP—plotted in the same format as Fig. 1a. All three conservation scores indicate higher evolutionary conservation for RDH genes compared with other genes. **b,** Distribution of conservation scores for RDH families (H1, H2A, H2B, H3, and H4) and for non-histone genes. All histone families show elevated conservation relative to other genes, with H1 being the most similar to the genome-wide average.

**Supplementary Fig. 7:**
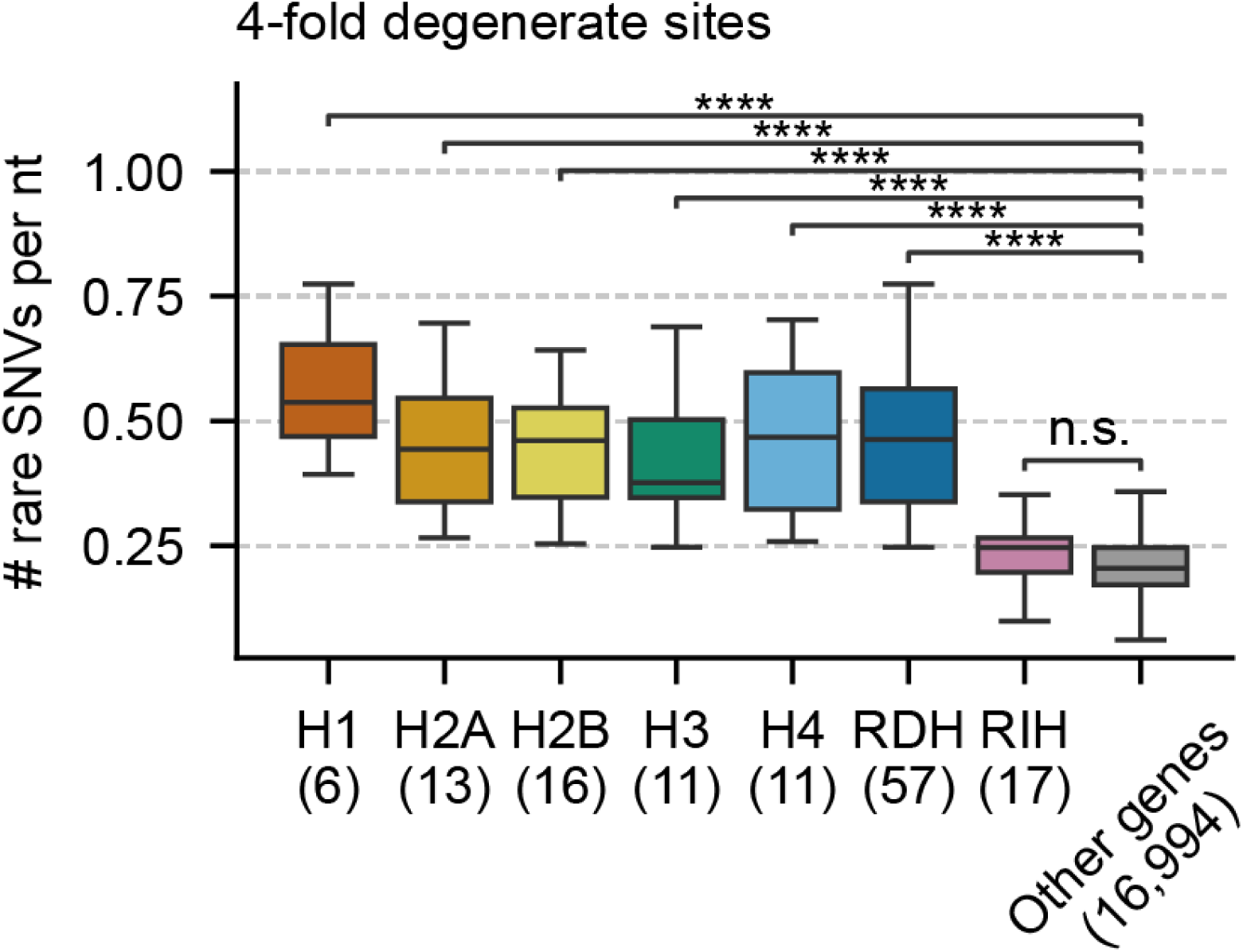
Rare variant density across gene sets at 4-fold degenerate sites. Bars represent the mean rare variant density per nucleotide at 4-fold degenerate sites for each histone family, the aggregated RDHs group (H1–H4), RIHs, and other genes. RDH denotes the combined set of canonical replication-dependent histone families (H1, H2A, H2B, H3, and H4). The number of genes in each set is indicated in brackets. *P* values were obtained by two-sided Mann–Whitney U tests (*: 0.01 < *P* ≤ 0.05; **: 0.001 < *P* ≤ 0.01; ***: 0.0001 < *P* ≤ 0.001; ****: *P* ≤ 0.0001).

**Supplementary Fig. 8:**
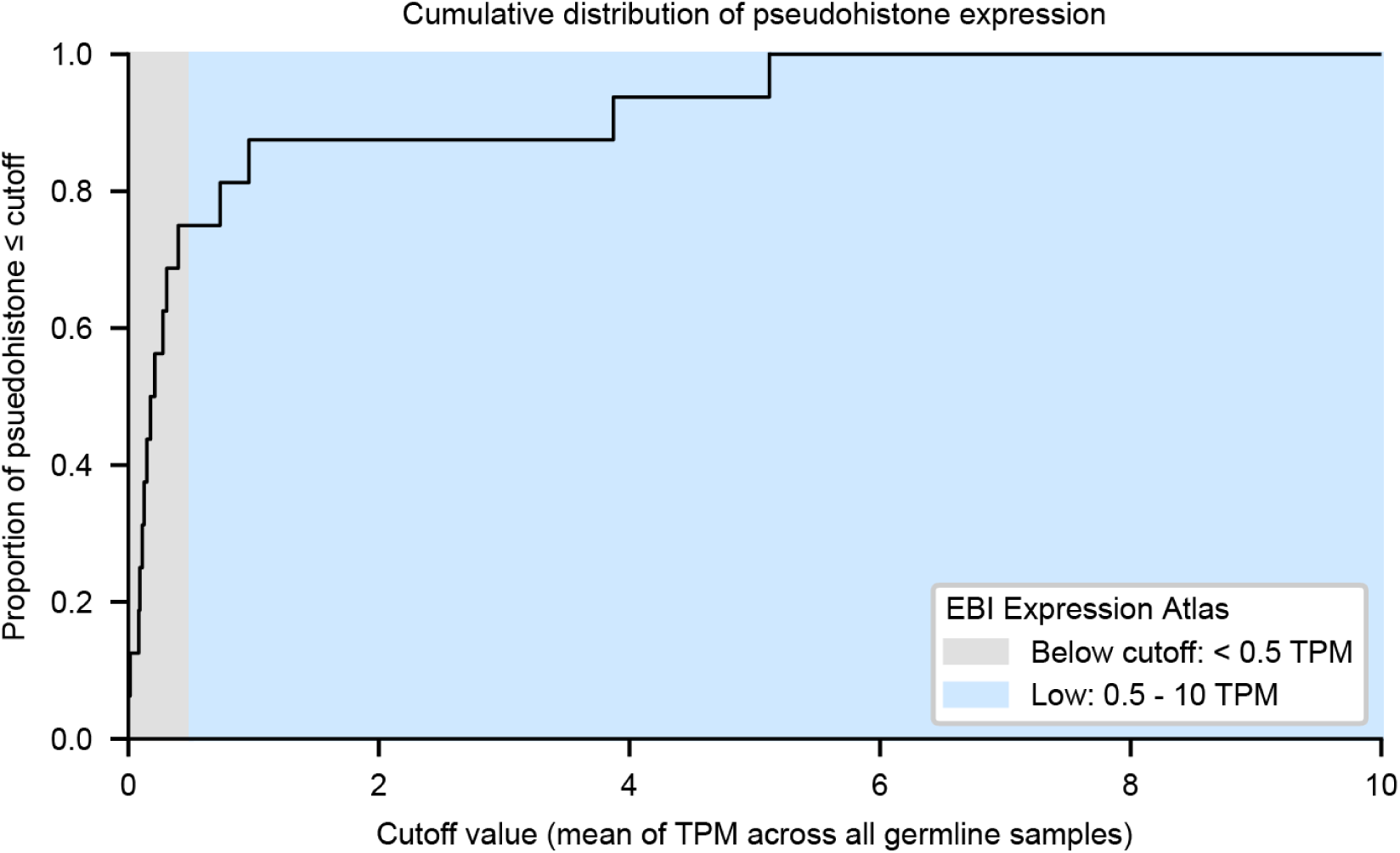
Germline expression profiles of pseudohistone genes. Cumulative distribution of pseudohistone gene expression levels in germline tissues. Most pseudogenes (12 of 16, 75%) show mean expression below 0.5 TPM, which falls below the EBI Expression Atlas threshold for detectable expression. The remaining four pseudogenes exhibit detectable but low expression (0.5-10 TPM). This overall minimal expression supports their classification as pseudogenes and suggests limited transcriptional activity in germline contexts.

**Supplementary Fig. 9:**
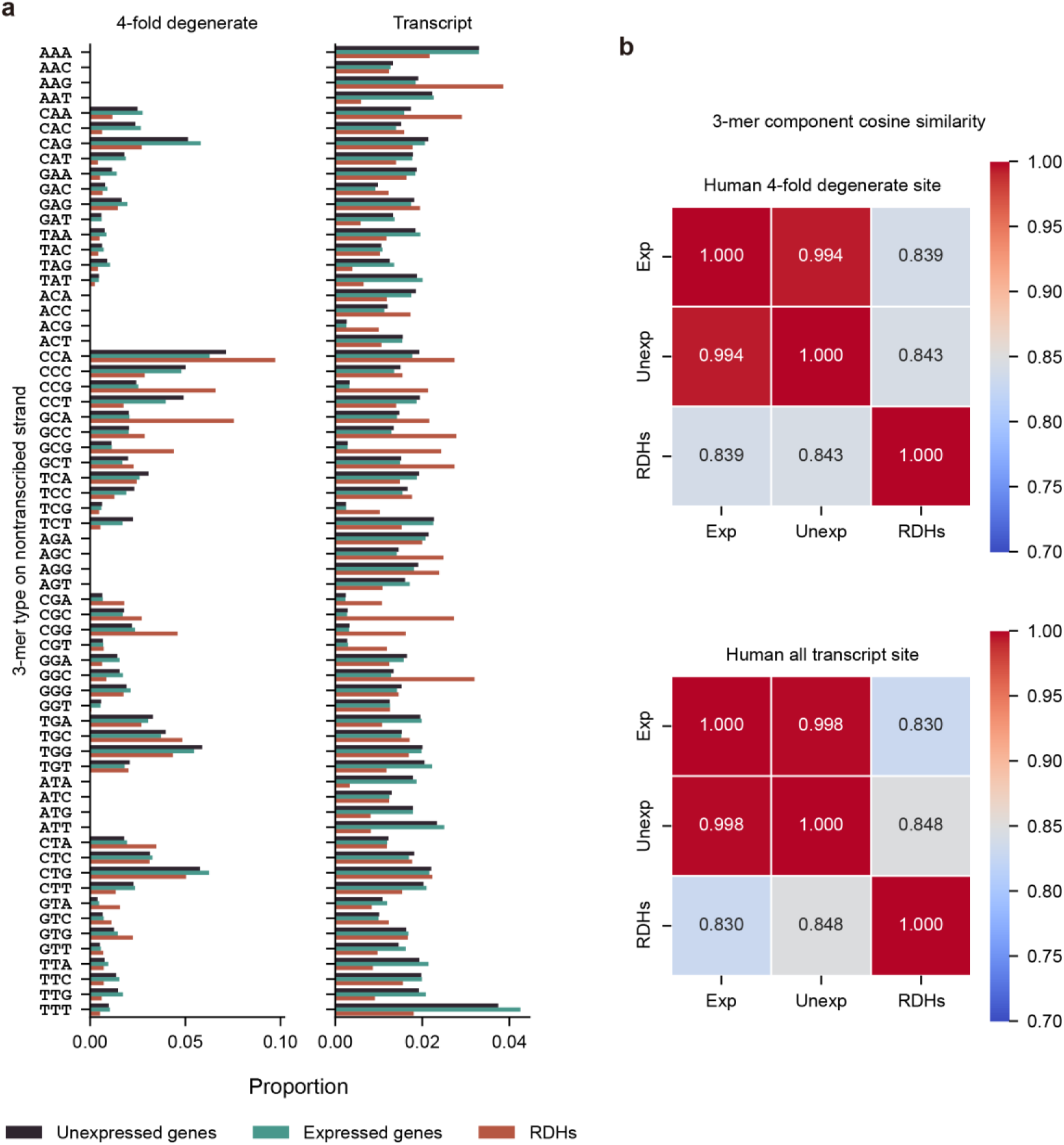
Sequence composition differs across gene groups. **a,** 3-mer sequence composition at 4-fold degenerate sites (left) and across full transcript sequences (right) for each gene group. **b,** Cosine similarity of 3-mer composition among gene groups. RDH genes show a distinct composition compared to the highly similar expressed and unexpressed gene groups.

**Supplementary Fig. 10:**
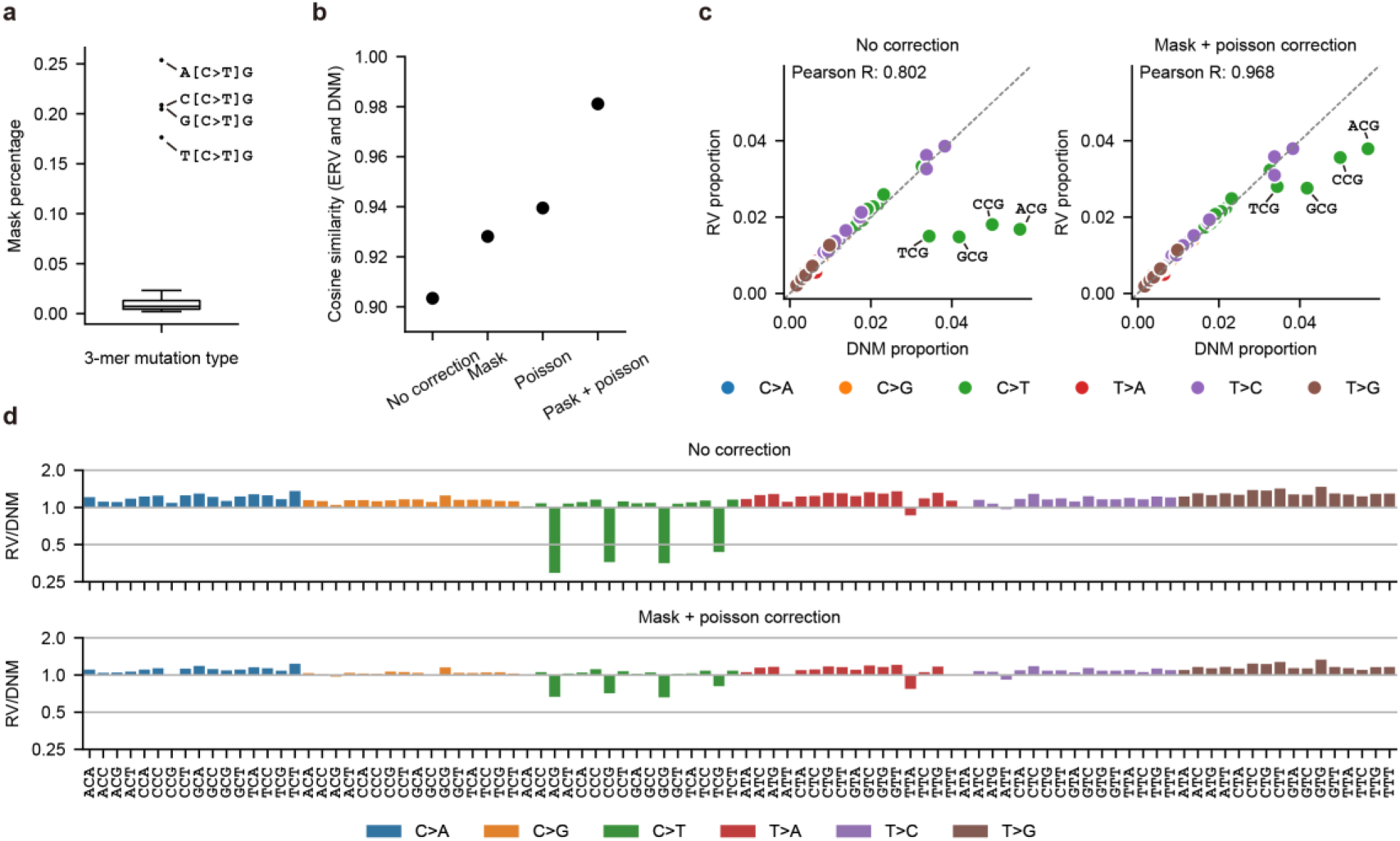
Calibration of rare variant spectrum against *de novo* mutations in humans. **a,** Proportion of masked sites for different 3-mer mutation contexts. The highest masking rates occurred at CpG sites (A[C>T]G, C[C>T]G, G[C>T]G, T[C>T]G), indicating prevalent mutation recurrence**. b,** Cosine similarity between rare variant (RV) and *de novo* mutation (DNM) spectra under different correction methods. Combined masking and Poisson correction achieved the highest similarity. **c,** Scatter plots of RV versus DNM mutation proportions before and after correction. Calibration improved concordance, though CpG variants in RVs remain slightly underestimated due to masking of recent mutations. **d,** Ratio of RV to DNM counts across 3-mer contexts. The combined correction method effectively minimized systematic bias from mutation saturation.

**Supplementary Fig. 11:**
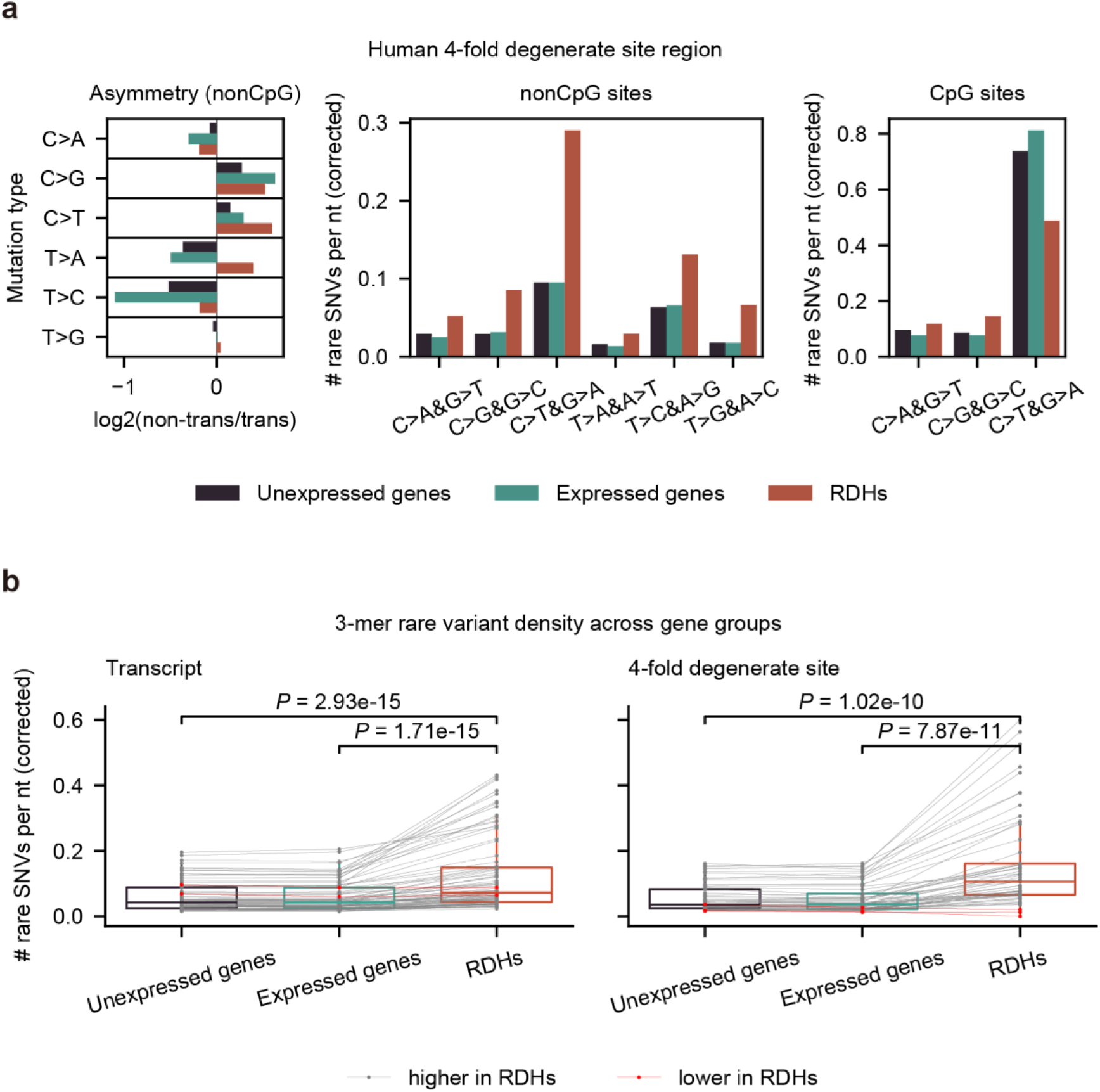
Comparison of rare variant densities among different 1-mer and 3-mer contexts at 4-fold degenerate sites. **a,** Strand asymmetry (at non-CpG sites) and calibrated spectrum of rare variants at 4-fold degenerate sites. 3-mer compositions were rescaled to match that of expressed genes (transcript level). **b,** Corrected rare SNV density for 3-mer contexts (restricted to non-CpG-context 3-mers with >50 occurrences in RDH genes). *P* values were obtained using two-sided Wilcoxon signed-ranks tests. Each line corresponds to a single 3-mer substitution type; red lines indicate lower rare SNV density in RDH genes relative to other groups, whereas grey lines indicate higher density in RDH genes.

**Supplementary Fig. 12:**
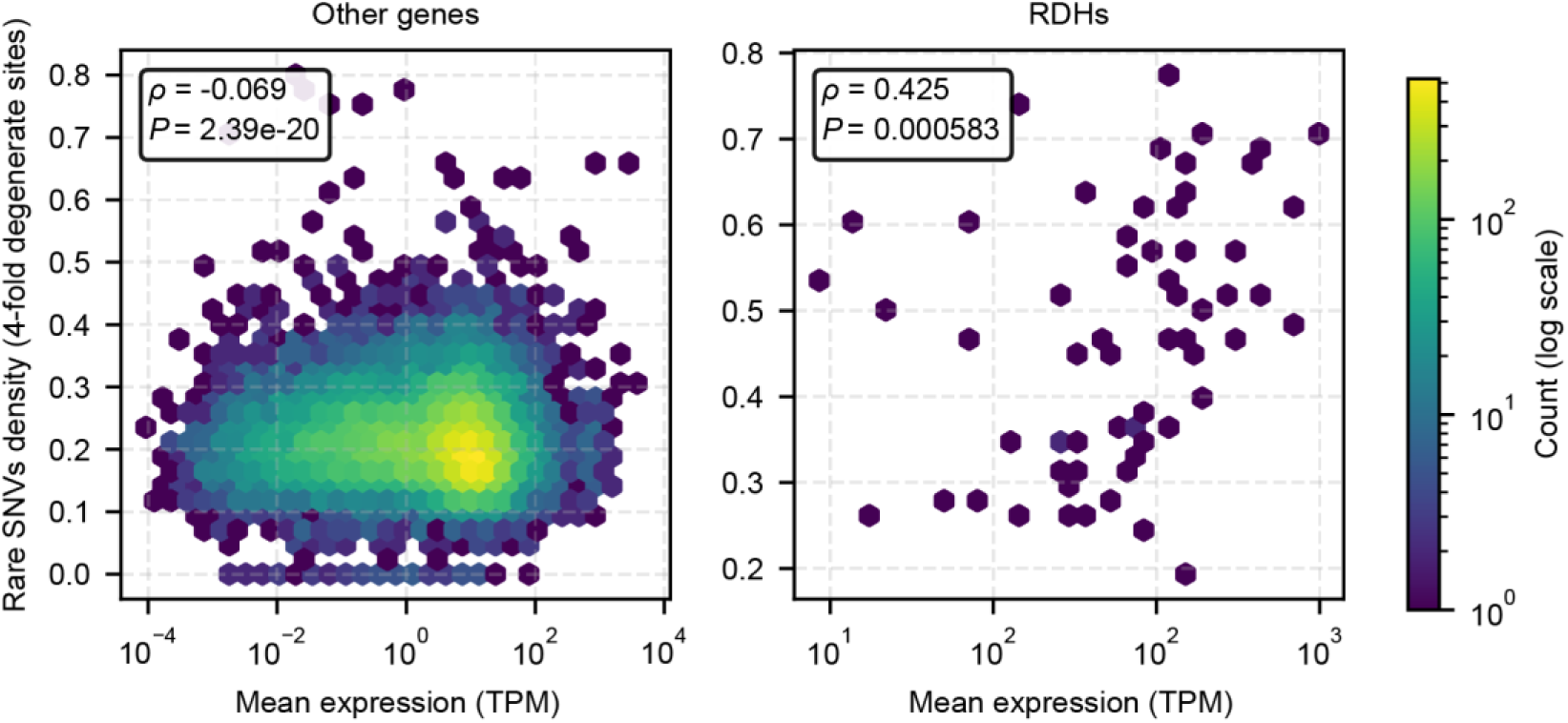
Correlation between germline expression level and rare SNV density. Scatterplot of germ cell expression versus rare SNV density at 4-fold degenerate sites for individual genes. Correlation coefficients and *P* values were calculated using Spearman’s rank correlation. RDH genes show a strong positive correlation between expression and variant density (ρ = 0.425, *P* = 0.0006). In contrast, only a weak negative correlation was observed for other genes (ρ = −0.069, *P* = 2.39 × 10^-20^).

**Supplementary Fig. 13:**
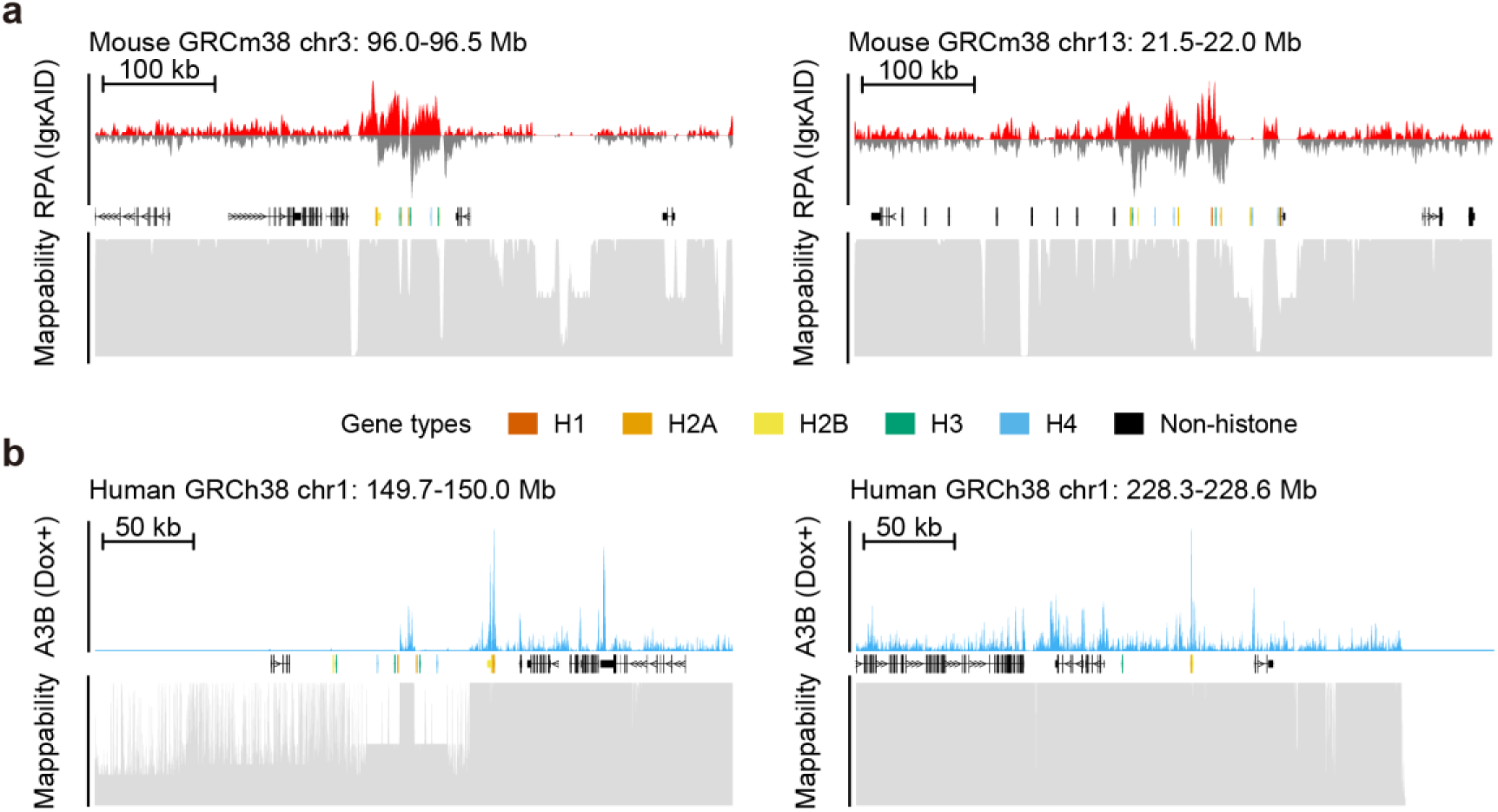
AID/APOBEC signal in other histone gene clusters. **a,** RPA ChIP-seq signal profile (top, RPKM values in AID-transgenic conditions) across two additional histone gene clusters in the mouse genome, with corresponding mappability scores (bottom, 0-1 scale calculated by GenMap). Some regions lacked mapped reads because of low mappability. **b,** A3B ChIP-seq signal profile (top, −log_10_(*P* value) under doxycycline-induced conditions) across two additional histone gene clusters in the human genome, with corresponding mappability scores.

**Supplementary Fig. 14:**
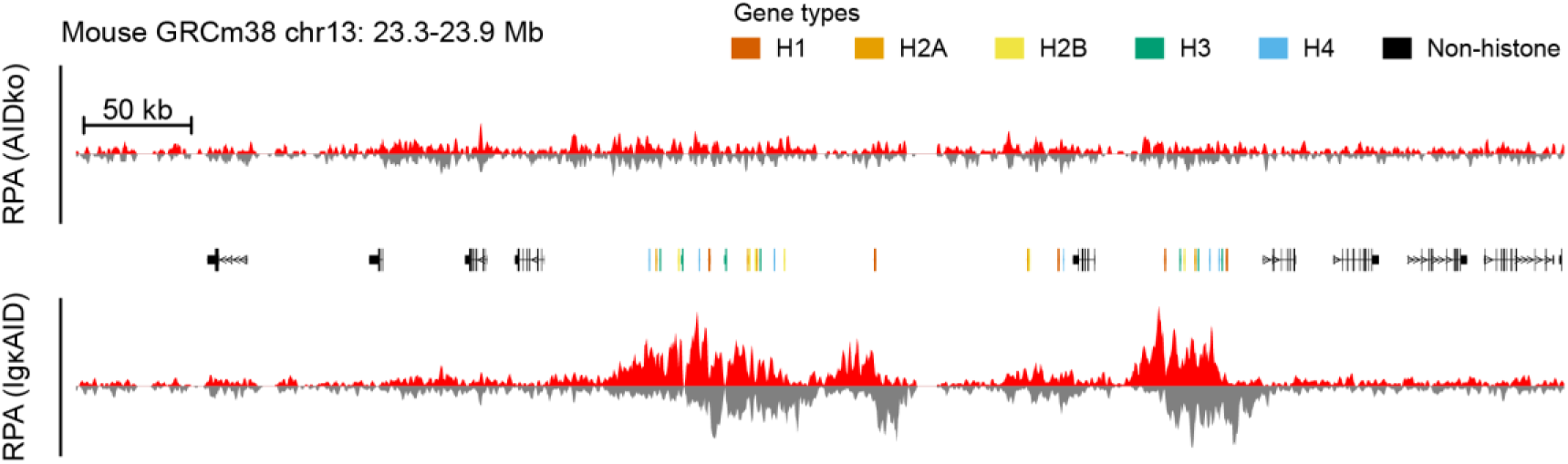
AID-dependent strand-asymmetric accumulation of RPA at histone genes. Strand-specific RPA ChIP–seq signal (RPKM) in AID-knockout (top) and AID-transgenic (bottom) cells. Strand-asymmetric RPA accumulation around histone genes is observed only in AID-transgenic cells, indicating that RPA loading at these loci is AID dependent.

**Supplementary Fig. 15:**
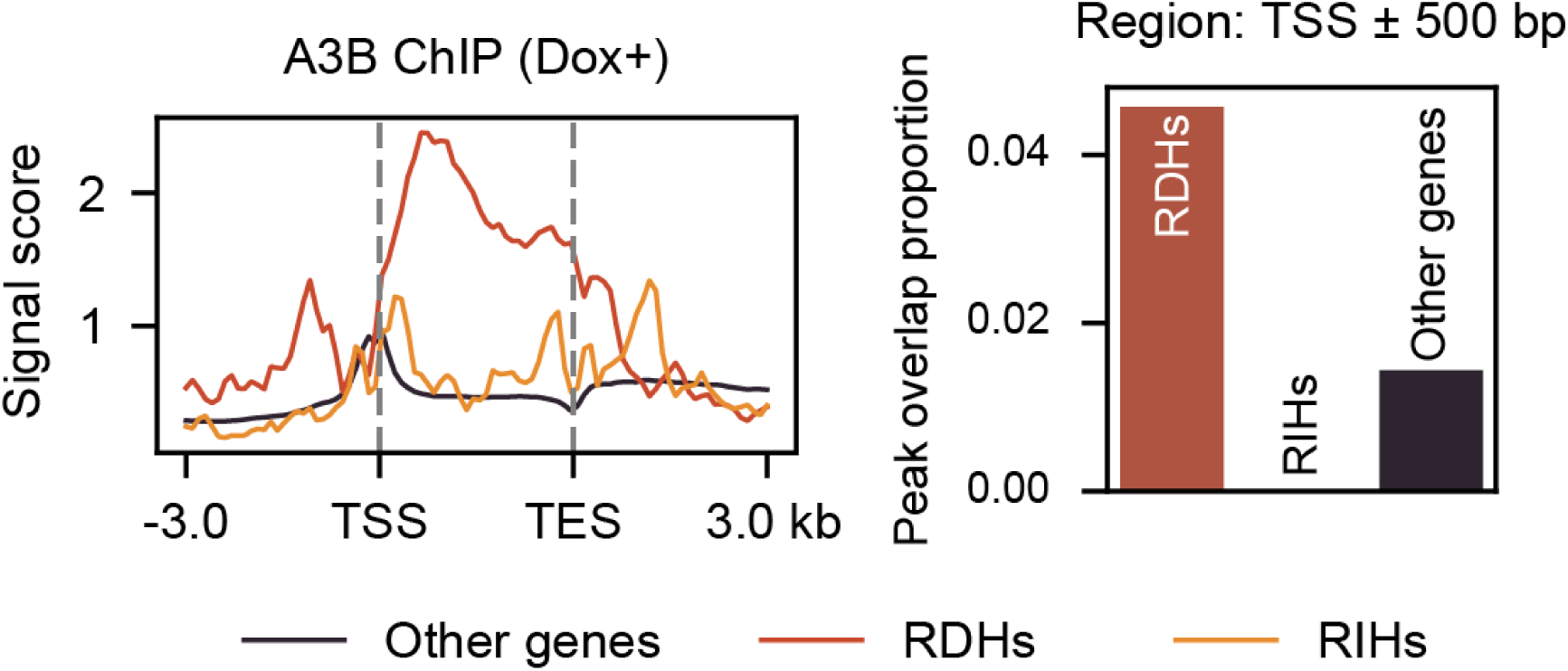
APOBEC3B peaks colocalize with RDH genes. A3B ChIP-seq in doxycycline-induced cells showing the −log_10_(*P*) signal profile (left) and the proportion of TSS regions (TSS ± 500 bp) overlapping A3B peaks across gene groups (right). RDH genes exhibit a markedly higher overlap with A3B peaks compared with other genes, and Fisher’s exact test confirms a significant enrichment (*P* < 2.2 × 10^-16^).

**Supplementary Fig. 16:**
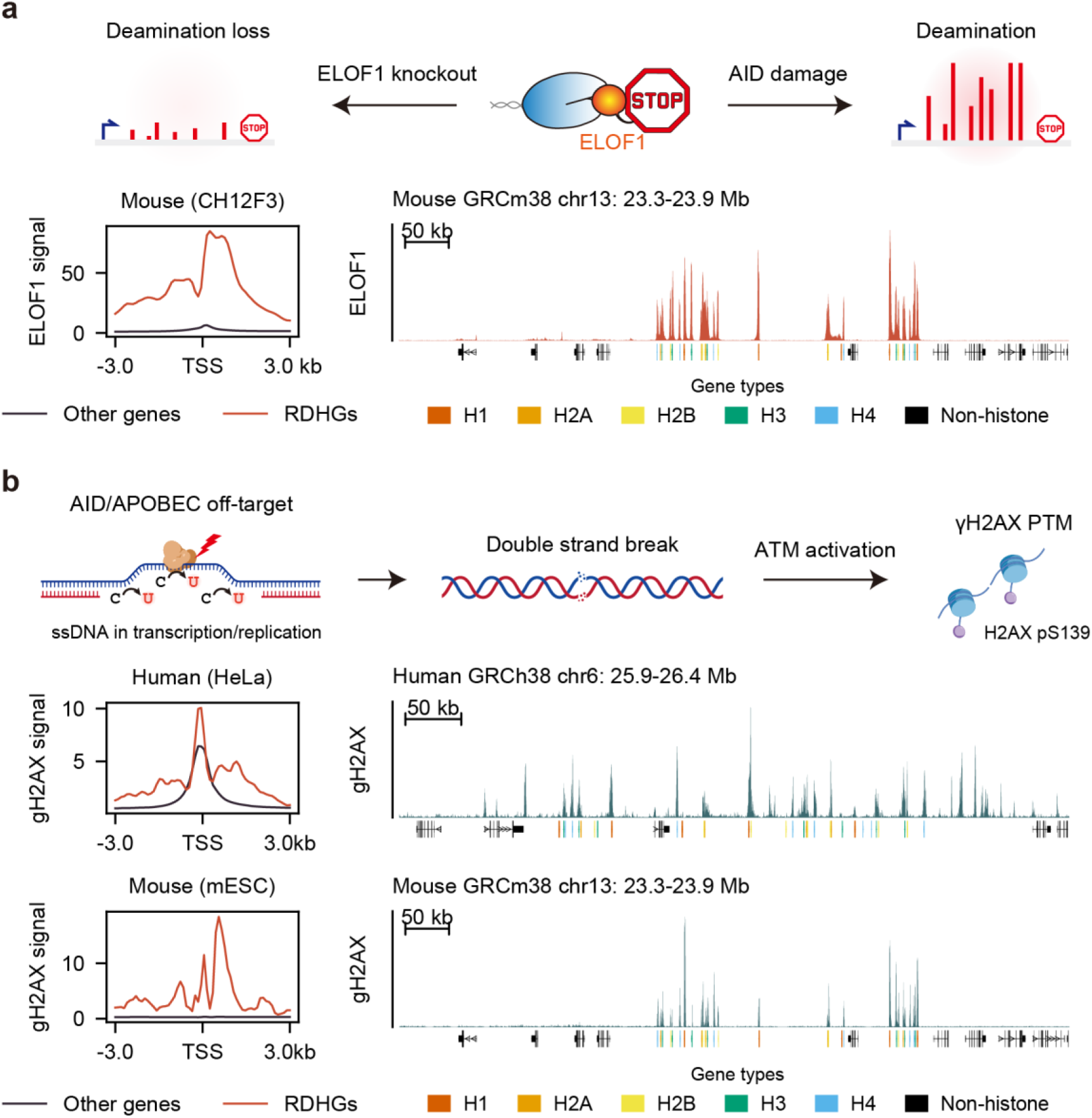
Indirect evidence supports AID/APOBEC activity at RDH genes. **a,** ELOF1, a factor required for AID targeting^28^, serves as additional indirect evidence (top). Comparison of ELOF1 ChIP-seq signal profile across gene groups (bottom left) and the genomic track within the major mouse RDH gene cluster (bottom right) are shown. **b**, AID/APOBEC-induced double-strand breaks lead to γH2AX modification^24,29,30^ (top). Comparison of signal profiles of γH2AX ChIP-seq (bottom left) and genomic tracks within the major RDH gene cluster in human and mouse (bottom right) are shown. Cell line names are given in the brackets.

**Supplementary Fig. 17:**
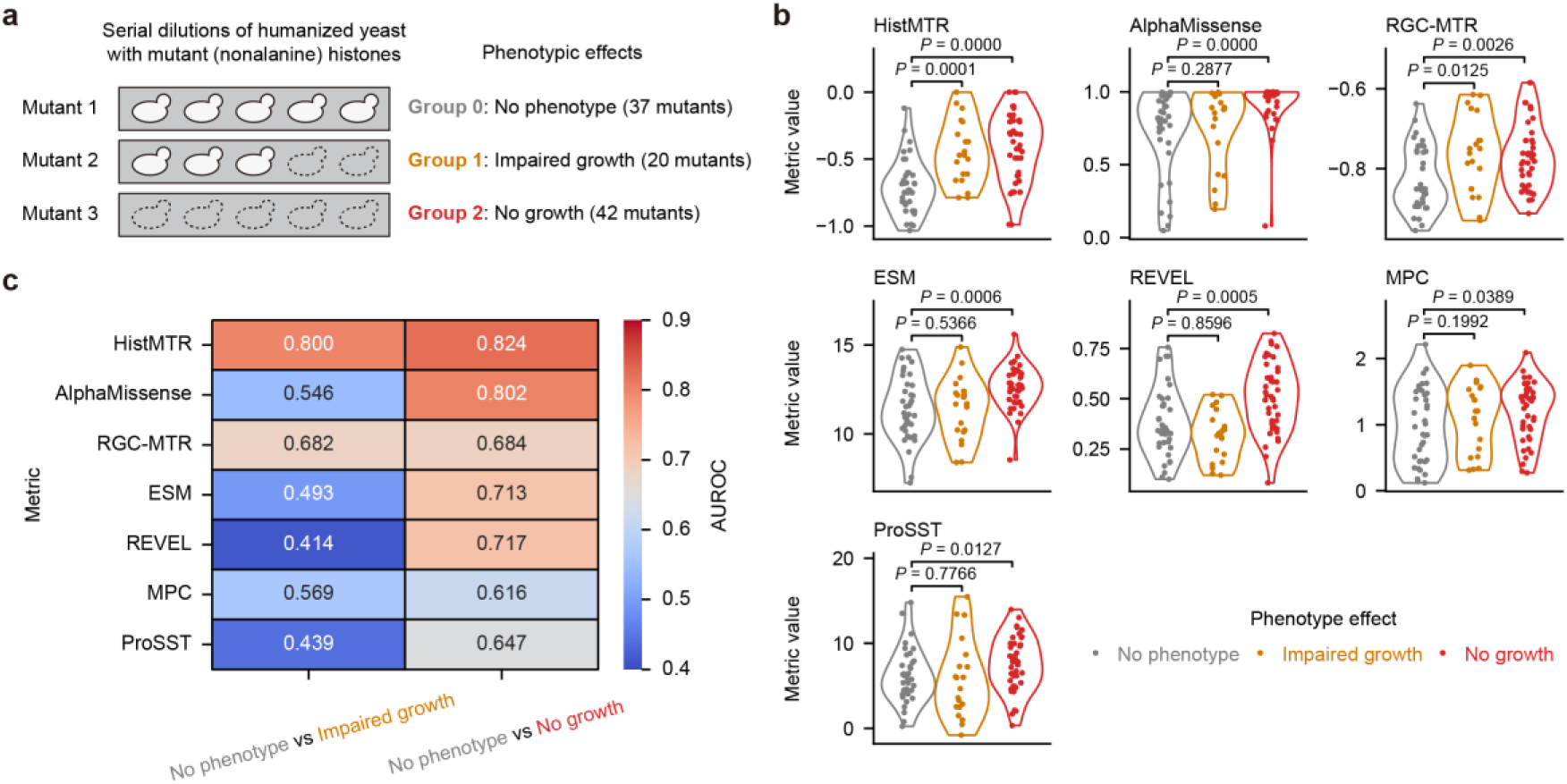
Performance of predictors across phenotypic severity classes. **a,** Illustration of phenotypic consequence categories (‘No phenotype’, ‘Impaired growth’, and ‘No growth’) for RDH mutations as defined by Bagert et al.^12^. Numbers of mutants used for benchmarking are indicated for each category. **b,** Distributions of predictor scores for RDH mutations in each phenotypic class. Scores were direction-harmonized (inverted where necessary) so that larger values indicate greater predicted deleteriousness. *P* values were computed with one-sided Mann–Whitney U tests under the alternative hypothesis that scores for ‘no phenotype’ variants are lower than those for the other two classes. Only HistMTR and RGC-MTR exhibited significant difference (*P* < 0.05) between ‘no phenotype’ and ‘impaired growth’. **c,** Heat map of AUROC values for pairwise classification between phenotypic classes. Variant-centric predictors (HistMTR, RGC-MTR, MPC: leveraging missense variation and/or sequence constraint) performed better at distinguishing intermediate-effect variants; notably, HistMTR achieved comparable AUROCs in both ‘no phenotype vs impaired growth’ and ‘no phenotype vs no growth’ comparisons.

**Supplementary Fig. 18:**
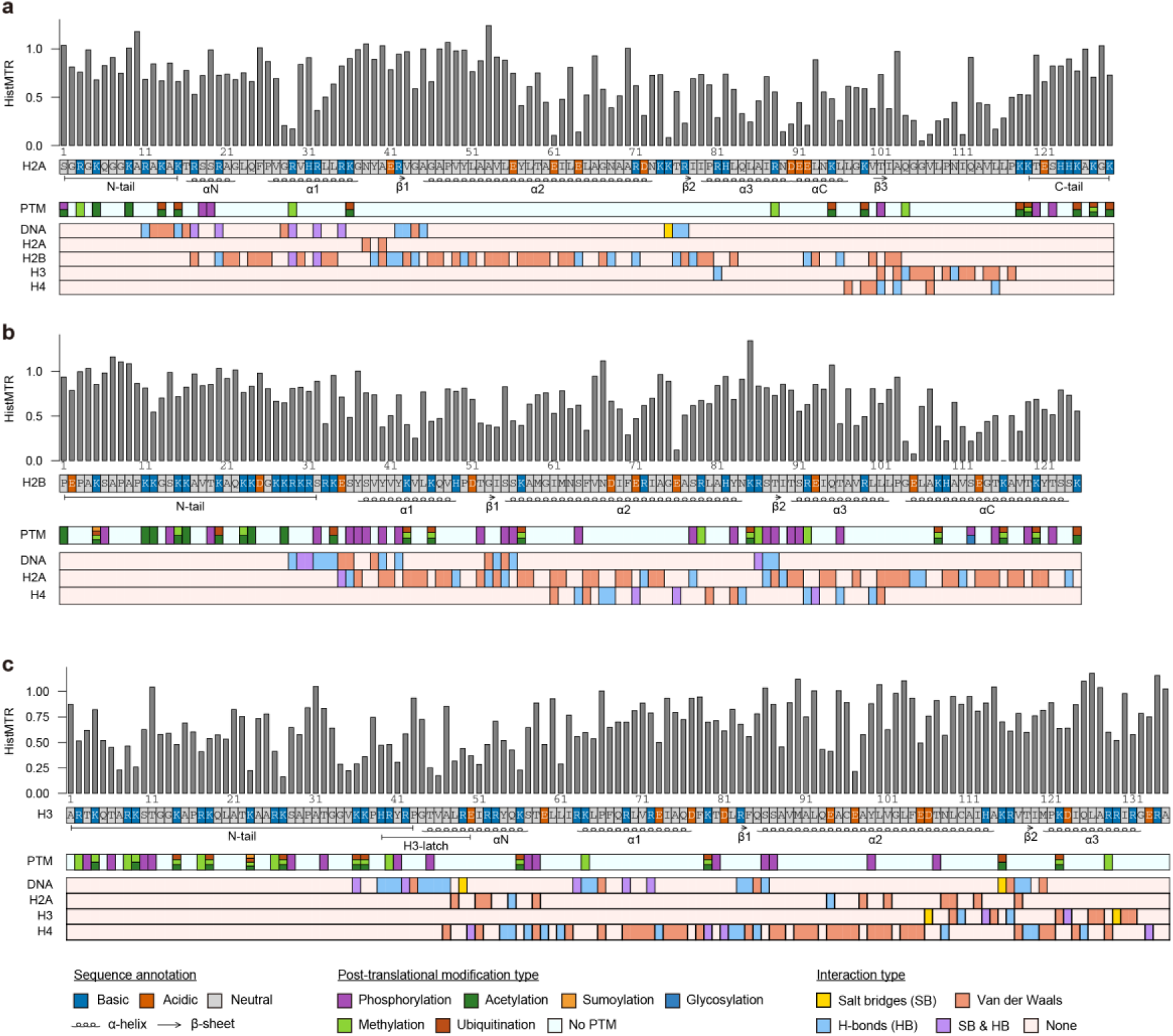
HistMTR profiles of H2A, H2B and H3 RDH families. HistMTR profiles for core RDH gene families **(a)** H2A, **(b)** H2B, and **(c)** H3. Other properties of each amino acid are given below the HistMTR profile. The presentation format and related annotations are consistent with Fig. 3c.

**Supplementary Fig. 19:**
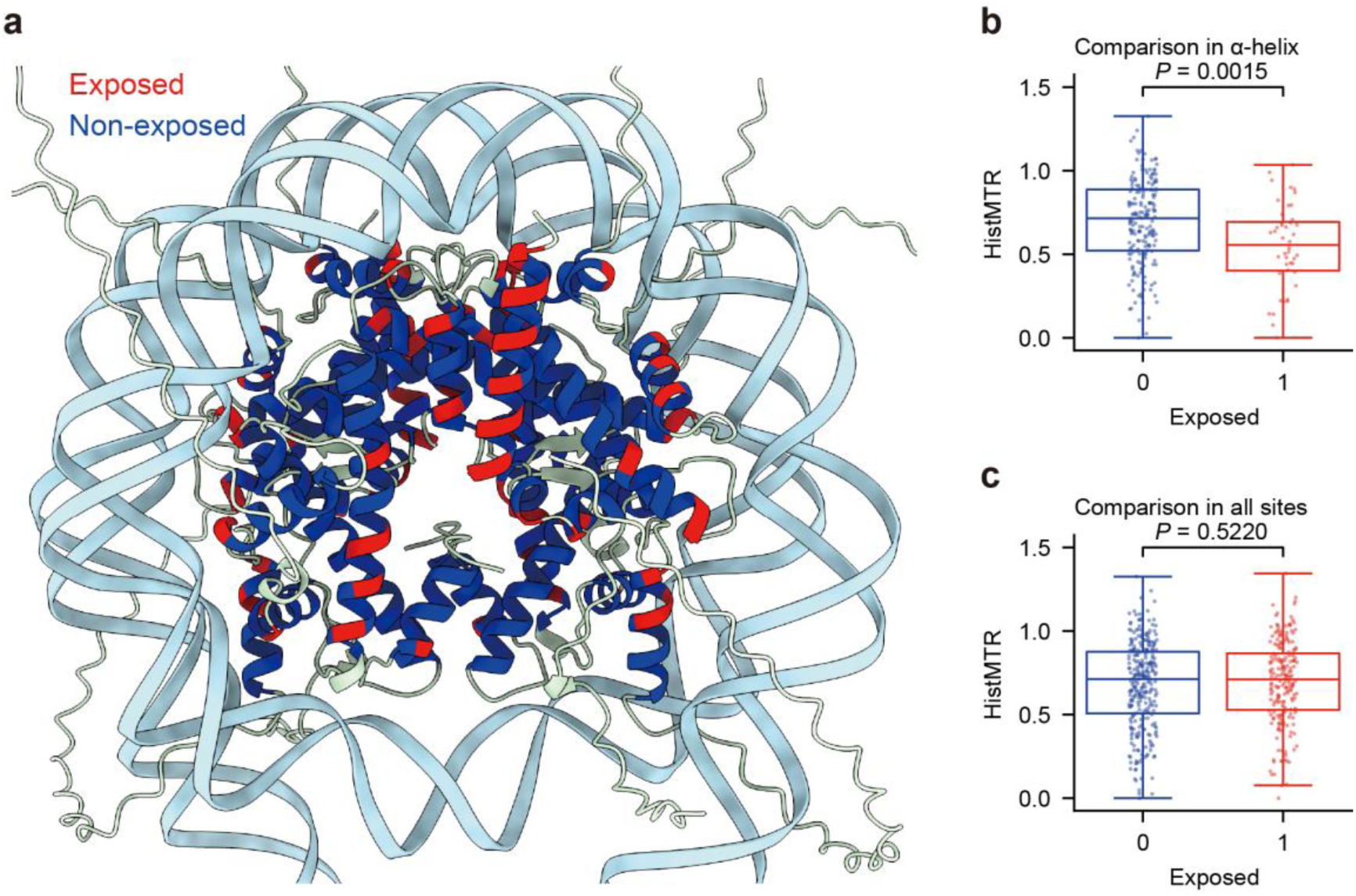
Functional constraint on surface-exposed helical residues. **a,** Structure of the nucleosome (DNA in light blue; histone H1 omitted) with helical residues colored by exposure status (red, exposed; blue, non-exposed; see Methods). **b,** Distribution of HistMTR scores for exposed versus non-exposed residues located on alpha-helices. Exposed residues show significantly lower HistMTR (greater constraint; *P* value from a two-sided Mann-Whitney U test). **c,** The same comparison for all residues in the nucleosome, which shows no significant difference.

**Supplementary Fig. 20:**
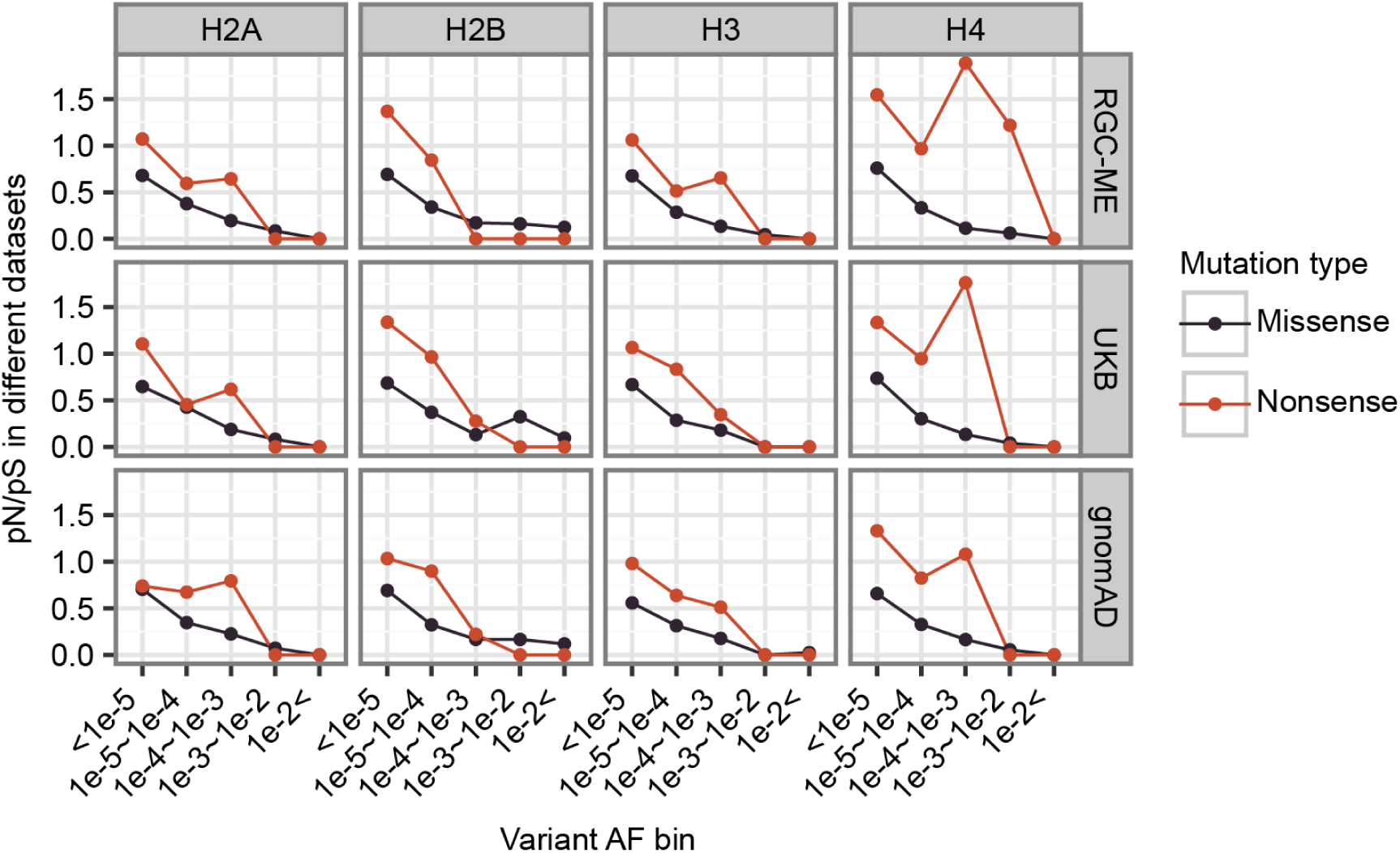
Comparative constraint analysis across RDH gene families. Comparison of missense and nonsense pN/pS ratios for individual RDH gene families.

**Supplementary Fig. 21:**
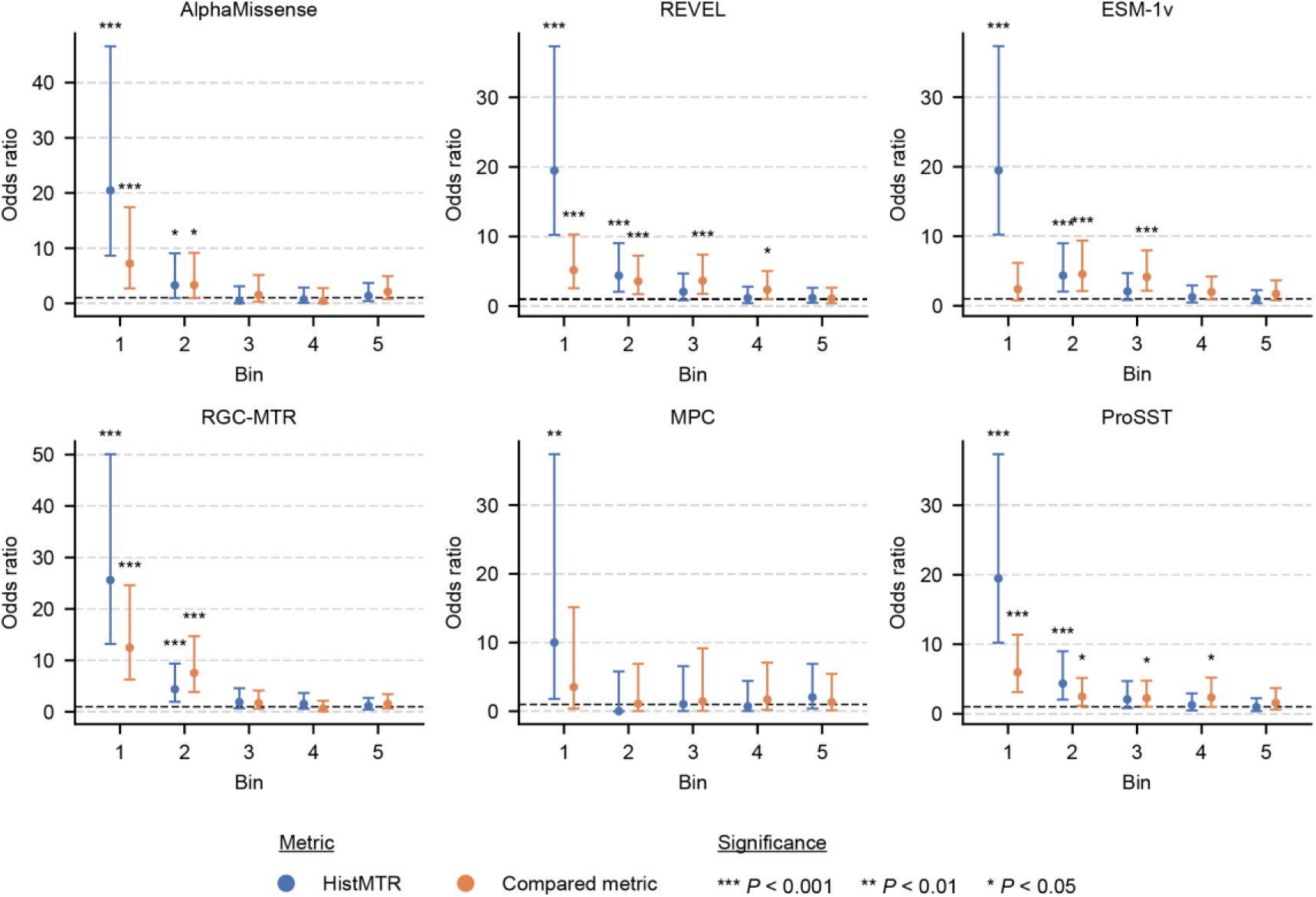
Enrichment of missense DNMs in DD patients across RDH missense sites stratified by deleteriousness predictors. Enrichment of missense DNMs in DD patients across core RDH missense sites stratified by deleteriousness predictors. Because the number of missense sites with annotations differed across predictors, we restricted each comparison to missense sites annotated for the corresponding metric. Missense sites were then divided into five bins of equal size according to HistMTR or the corresponding predictor. ORs and *P* values were calculated for each bin. Across predictors, the first bin defined by HistMTR consistently showed the highest OR, supporting its improved ability to prioritize DD risk sites. Odds ratios and *P* values were calculated using Fisher’s exact test.

**Supplementary Fig. 22:**
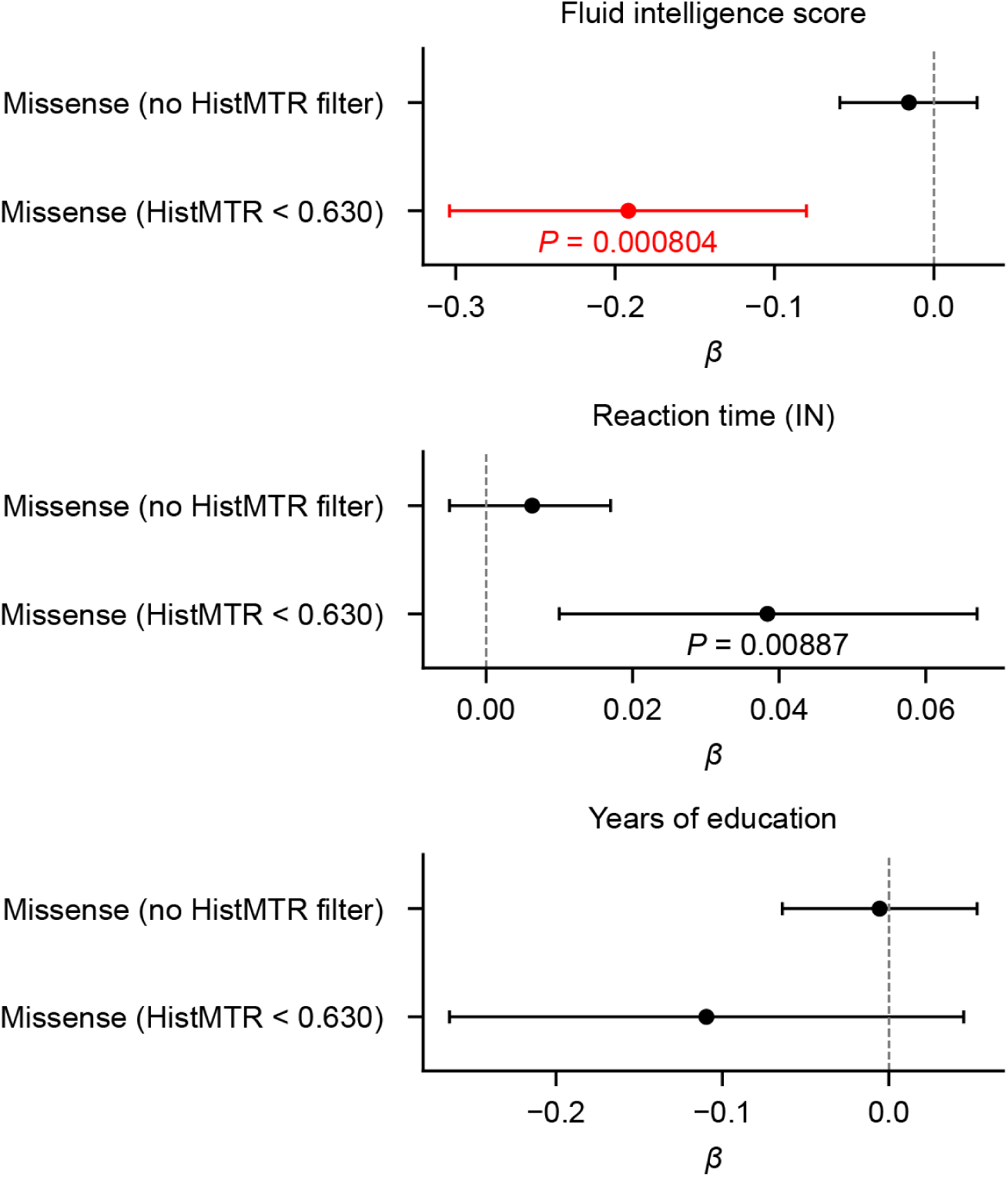
Association tests between cognitive phenotypes and rare missense burden. Association tests between cognitive phenotypes and rare missense burden were performed with and without the HistMTR filter. For fluid intelligence, *β* values represent effect sizes on standardized scores (range, 1–13); for reaction time (inverse-normal transformed), *β* values represent effect sizes in standard-deviation units; and for years of education, *β* values represent effect sizes on years of education attained. Error bars represent 95% confidence intervals; bars are colored red if the association remained significant after Bonferroni correction (adjusted *P* < 0.05). Raw *P* values < 0.05 are shown. Without HistMTR-based classification of putatively pathogenic missense variants, effect sizes decreased and *P* values increased, indicating that the HistMTR filter improved association signals.

**Supplementary Fig. 23:**
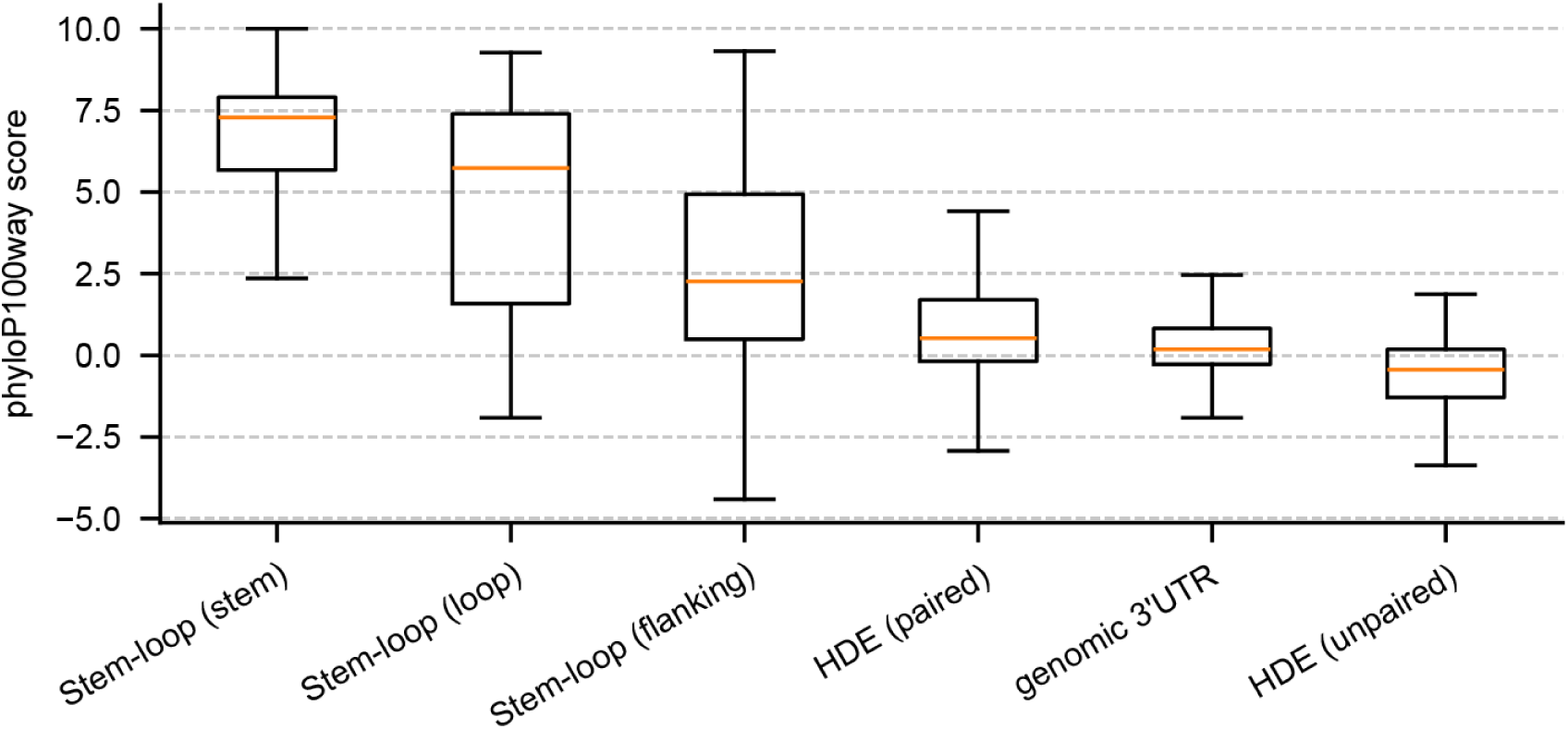
Evolutionary conservation of RDH gene regulatory elements (stem-loops and HDEs). Boxplots showing phyloP100way conservation scores for different non-coding regions in RDH genes. Regions are ordered by median conservation score. Within stem-loops, the stem and loop regions show stronger evolutionary constraint than the flanking sequences. Similarly, paired positions within HDEs are more conserved than unpaired regions.

**Supplementary Fig. 24:**
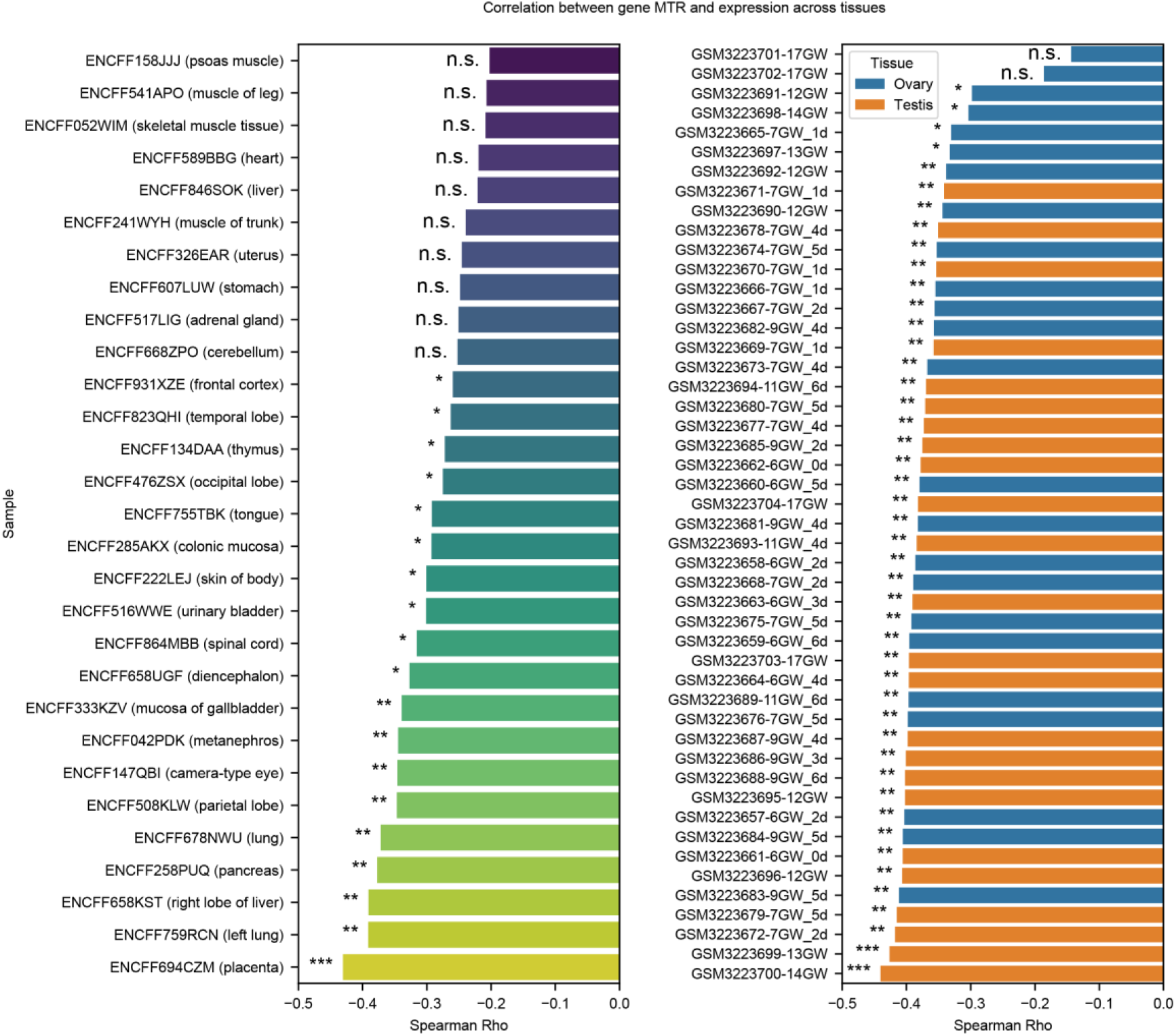
Correlation between constraint and expression in core RDH genes across tissues. Left: Expression data of somatic tissues from ENCODE (Methods). Right: Expression data from germline tissues (Methods). Spearman’s rank correlation coefficients and *P* values were calculated using gene-level MTR and TPM values for core RDH genes. *: 0.01 ≤ *P* < 0.05; **: 0.001 ≤ *P* < 0.01; ***: *P* < 0.001.

**Supplementary Fig. 25:**
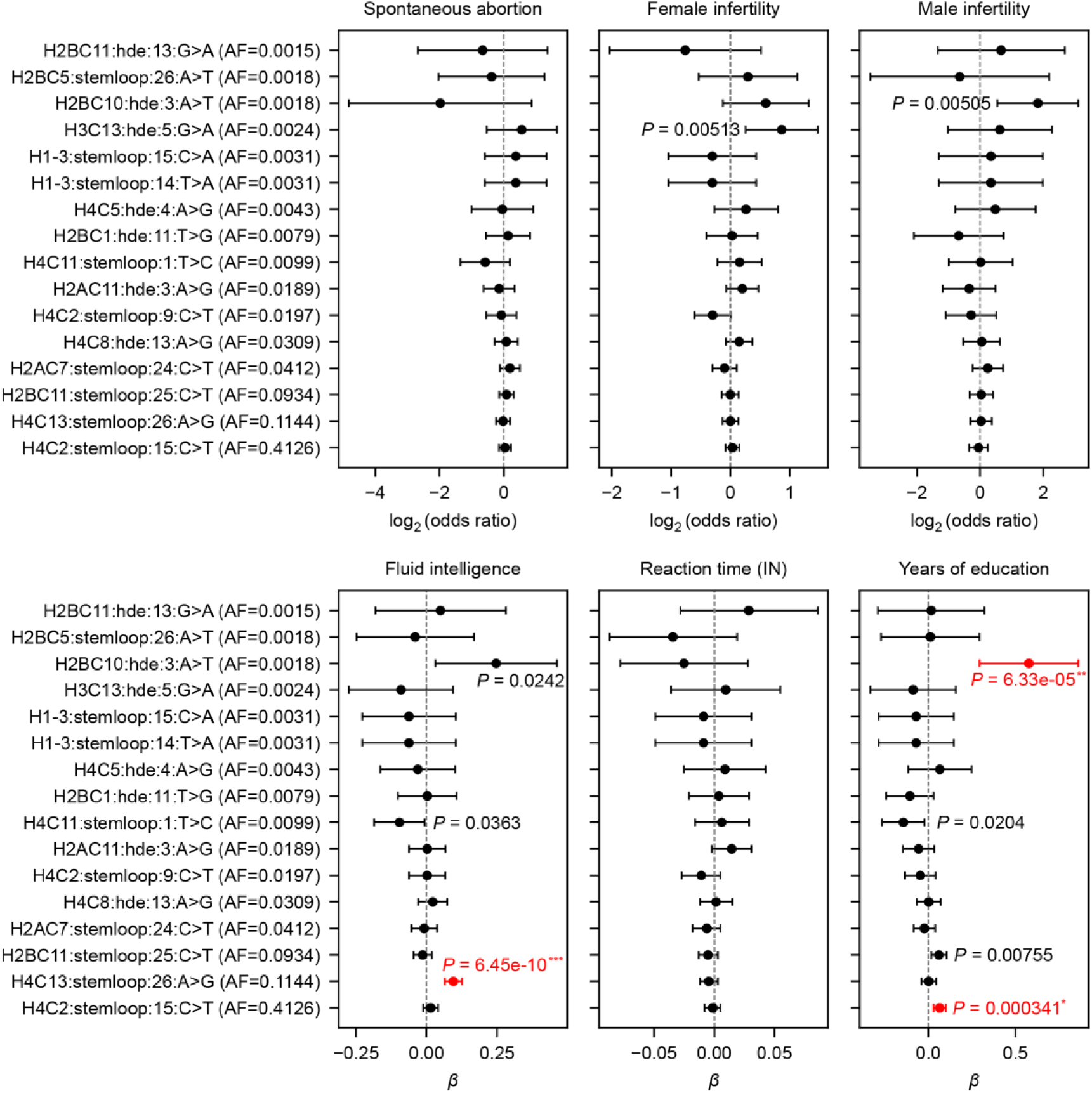
Association of common variants in stem-loops and HDEs of RDH genes with cognitive and reproductive phenotypes. The figure presents association results for six phenotypes: the top three panels show reproductive phenotypes, and the bottom three show cognitive phenotypes. Within each panel, variants are ordered by their allele frequency in the UK Biobank. Error bars represent 95% confidence intervals; bars are colored red if the association remained significant after Bonferroni correction (adjusted *P* < 0.05). Original *P* values are shown and annotated with asterisks based on the corrected significance thresholds (***: adjusted *P* < 0.001; **: 0.001 ≤ adjusted *P* < 0.01; *: 0.01 ≤ adjusted *P* < 0.05).

**Supplementary Fig. 26:**
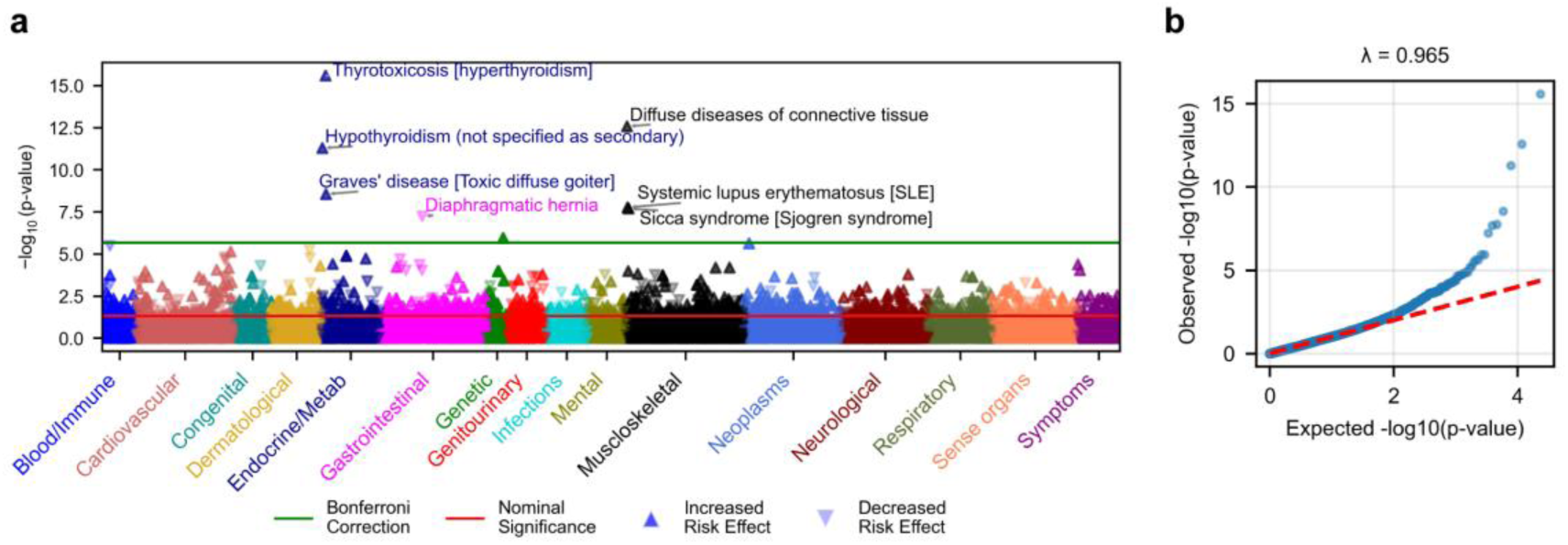
Phenome-wide association scan for common variants in stem-loops and HDEs of RDH genes. **a.** Manhattan plot displaying results from the PheWAS analysis of common regulatory variants (stem-loop or HDE variants) in RDH genes. Points represent phecodes, colored by phenotypic category. Phecodes that possess known association with other non-histone genes in adjacent regions were masked (Methods). **b.** Quantile–quantile plot of association *P* values. The genomic inflation factor (λ_GC_ = 0.960) indicates well-calibrated test statistics with minimal inflation.

**Supplementary Fig. 27:**
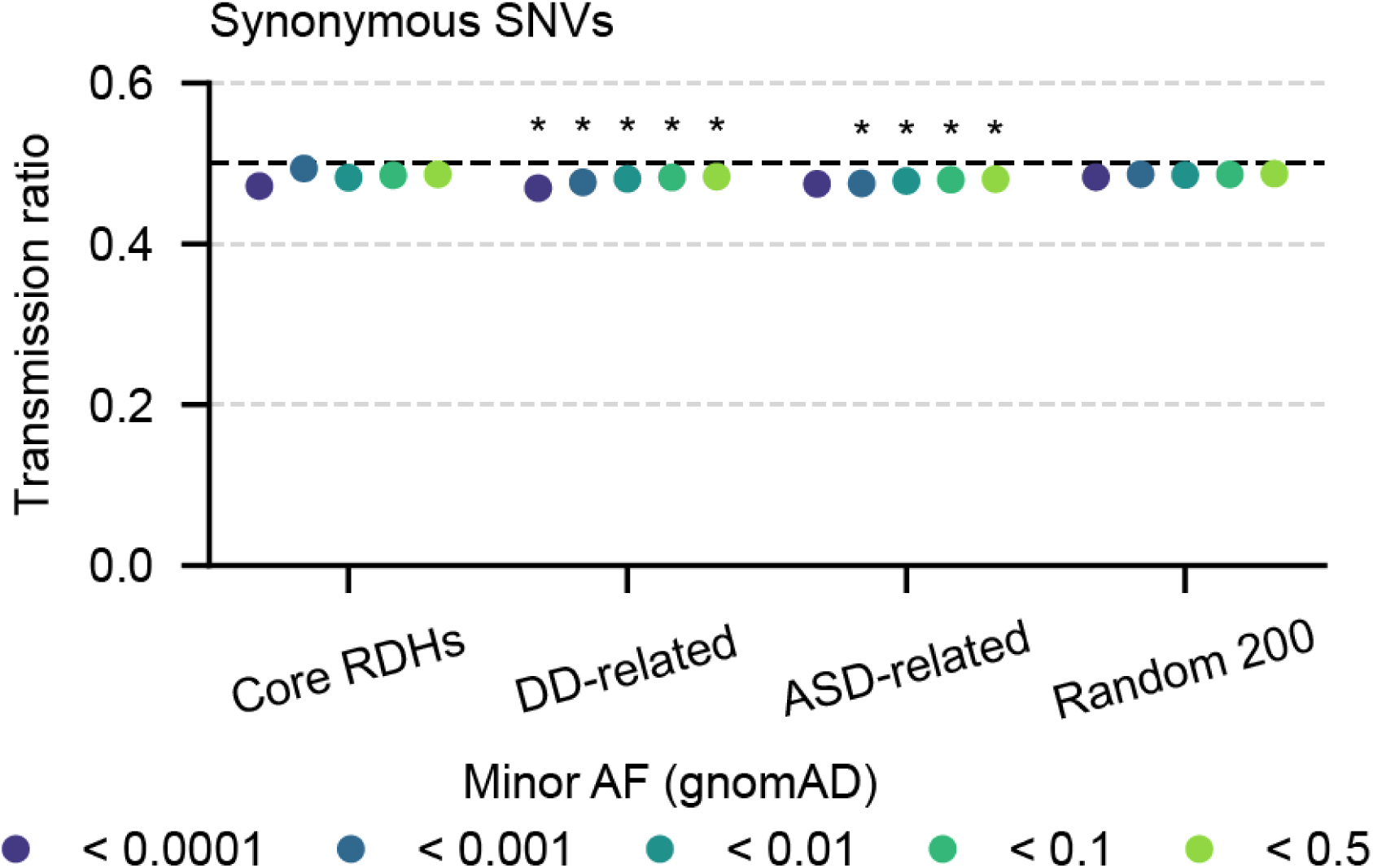
Transmission ratios of synonymous variants across gene groups. Transmission ratios of synonymous variants across gnomAD AF bins for different gene groups. The dashed line (ratio = 0.5) indicates the neutral expectation. Asterisks denote significance after Benjamini–Hochberg correction (adjusted *P* < 0.05) based on simulation analysis.

**Supplementary Fig. 28:**
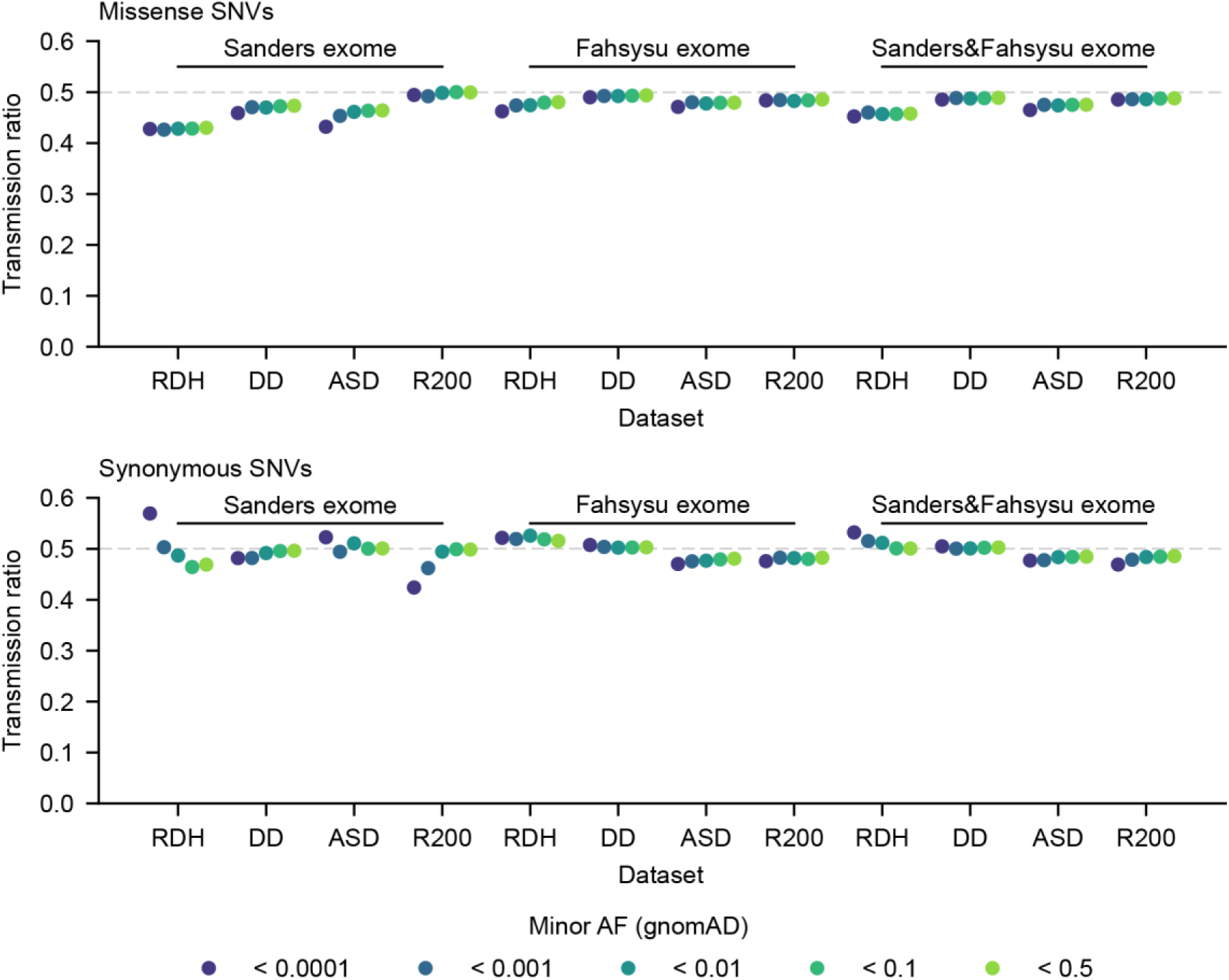
Transmission ratio distortion in independent exome trio datasets. Analysis of missense (top) and synonymous (bottom) variant transmission across gnomAD AF bins for different gene groups in two independent exome datasets (‘Sanders’ exome and ‘Fahsysu’ exome). Two datasets were also merged to increase power (‘Sanders&Fahsysu’ exome). These independent cohorts recapitulate the pattern observed in the 1KG data: RDH missense variants show lower transmission ratios than other gene groups, while their synonymous variants do not show no reduced transmission. The TRD in RHDs did not reach statistical significance in the exome datasets (**Supplementary Table 14**), likely due to the smaller number of informative variants present in these datasets. Moreover, the weaker TRD patterns observed in the two exome datasets compared to the 1000 Genomes Project data likely reflected a shorter duration of exposure to selection and/or ascertainment bias toward individuals with disease phenotypes.

**Supplementary Fig. 29:**
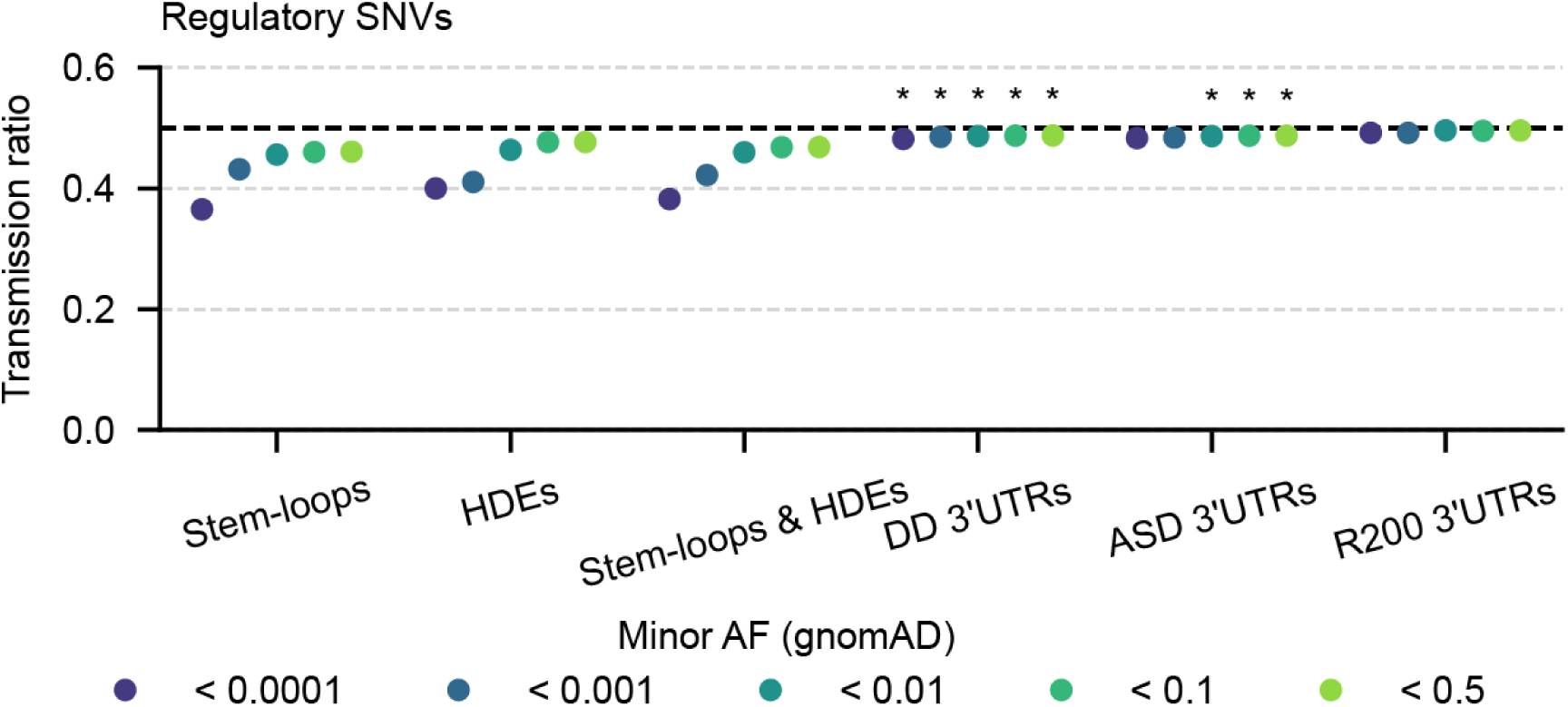
Transmission ratio distortion in regulatory elements. Analysis of regulatory variant transmission across gnomAD AF bins for different regulatory elements in 1KG trio data. Derived variants in RDH regulatory elements (stem-loops and HDEs) show lower transmission ratios than other gene groups. However, the effect did not reach statistical significance (**Supplementary Table 15**), likely due to the smaller number of informative variants present in stem-loops and HDEs. Asterisks denote significance after Benjamini–Hochberg correction (*P* < 0.05) based on simulation analysis

**Supplementary Fig. 30:**
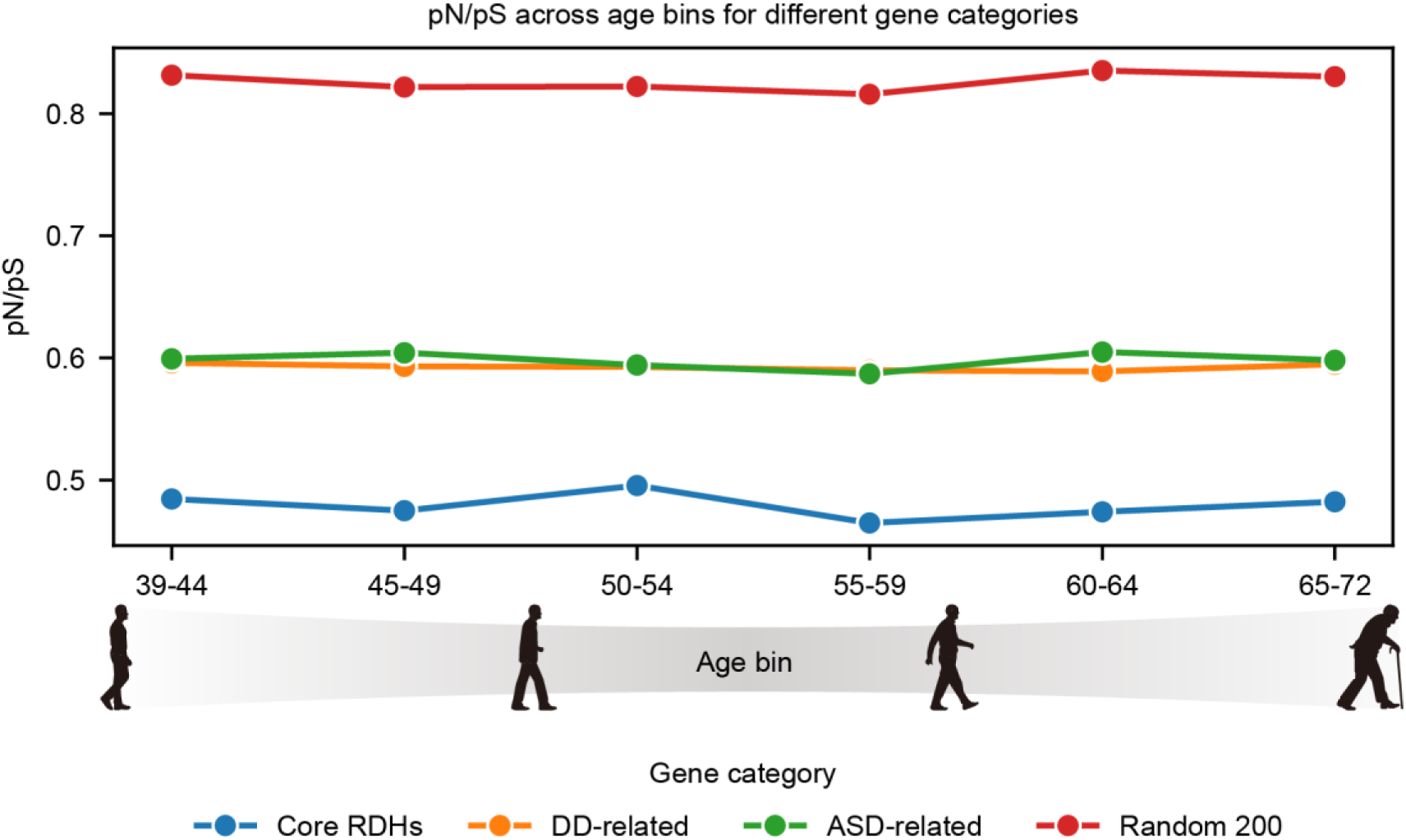
Missense pN/pS ratios remain stable across age bins. Analysis of missense pN/pS across age bins with UKB data reveals no significant age-dependent trends in any gene category (pairwise Fisher’s exact tests). For each age bin, 30,000 unrelated White British individuals were randomly sampled to control for sample size differences across bins. Note that the sequenced individuals in UKB are relatively old (>39 years old) and pN/pS ratios may exhibit age dependence at younger ages.

**Supplementary Fig. 31:**
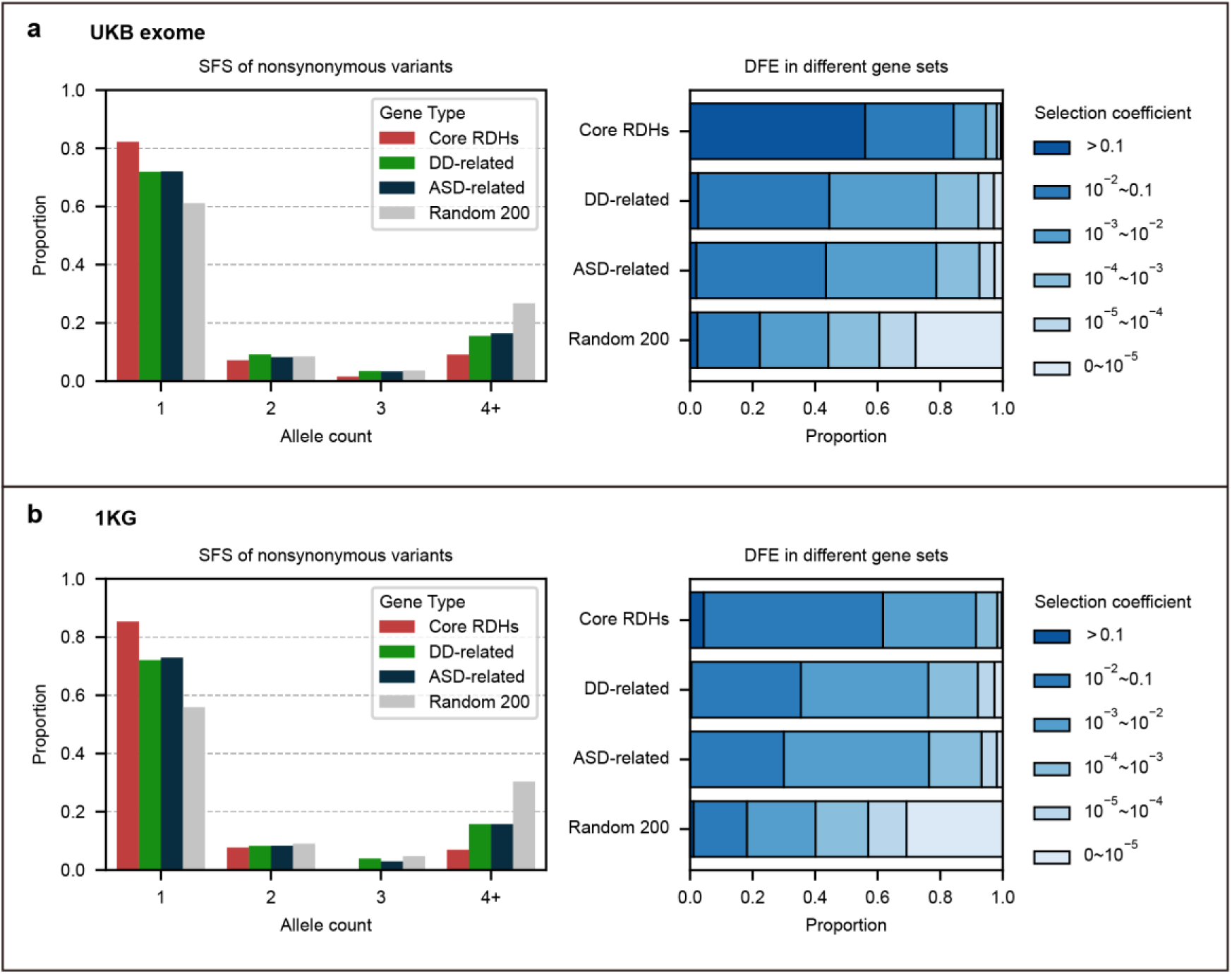
DFEs in UKB exome and 1KG datasets. **a.** UKB exome data: we randomly selected 350 unrelated White British individuals per age bin (2,100 total) and downsampled to 2,000 chromosomes. **b.** 1KG European data: we used 493 unrelated individuals labeled ‘EUR’ and downsampled to 900 chromosomes. For each dataset, the site frequency spectrum and inferred DFE are shown for nonsynonymous variants across gene categories. Although the proportion of the most deleterious class (|s| > 0.1) appears lower for core RDHs in 1KG, core RDHs consistently show a higher fraction of strongly deleterious variants (|s| > 10^-2^) in both datasets.

**Supplementary Fig. 32:**
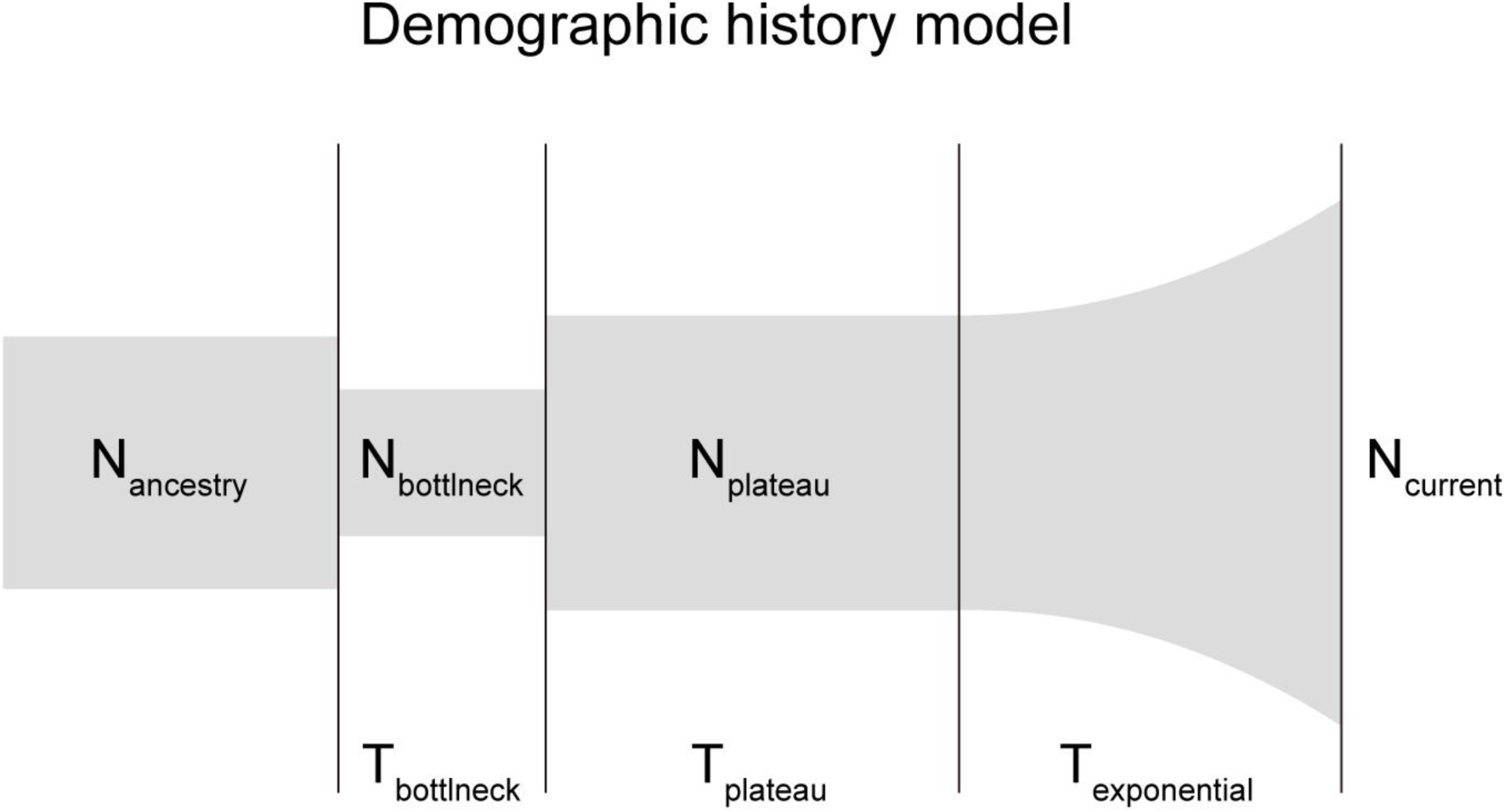
Demographic model for distribution of fitness effects analysis. The population history model includes an out-of-Africa bottleneck, subsequent recovery phase, and recent exponential growth (adapted from Kim et al.^114^). Parameter estimates for this model are provided in **Supplementary Table 21**.

## Supplementary Tables

Titles of Supplementary Tables are shown below. See details in the associated Excel file.

*Supplementary Table 1. Mutation signature analysis under different variant filtering and processing strategies*.

*Supplementary Table 2. HistMTR values for all possible missense mutations in core RDH genes*.

*Supplementary Table 3. Phenotypic effects for a subset of amino acid mutations in RDHs from a previous reported yeast library (Bagert et al. 2021) and associated deleteriousness metrics by different predictors*.

*Supplementary Table 4. Comparison of AUROC and AUPRC values of predictors distinguishing pathogenicity of RDH missense mutations in the previously reported yeast library (Bagert et al 2021)*.

*Supplementary Table 5. HistMTR values of all amino acid sites for each core RDH family*.

*Supplementary Table 6. Enrichment of missense DNMs in core RDH genes in DD patients*.

*Supplementary Table 7. Reported associations between core RDH genes and DD from DECIPHER and related studies*.

*Supplementary Table 8. Occurrence of missense DNMs within core RDH genes in DD patients (denovoWEST) and related annotations*.

*Supplementary Table 9. Association results for RDH rare coding variant burden with phenotypes*.

*Supplementary Table 10. PheWAS results of the burden analysis for putatively pathogenic rare missense variants in core RDH genes*.

*Supplementary Table 11. Association analysis of RDH rare regulatory variant burden with phenotypes*.

*Supplementary Table 12. Association results for common RDH regulatory variants with phenotypes*.

*Supplementary Table 13. PheWAS result for common regulatory variants in RDH genes*.

*Supplementary Table 14. Results of transmission ratio distortion (TRD) analysis for coding variants across gene groups and datasets*.

*Supplementary Table 15. Results of transmission ratio distortion (TRD) analysis for regulatory variants across regulatory element groups*.

*Supplementary Table 16. Distributions of fitness effects (DFEs) for different gene groups in different datasets*.

*Supplementary Table 17. List of species, genome assemblies, and population datasets used for analysis of RDH genes*.

*Supplementary Table 18. Genome coordinates (hg38) of stem-loops for human RDH genes*.

*Supplementary Table 19. Genome coordinates (hg38) of HDEs for human RDH genes*.

*Supplementary Table 20. ChIP-seq data sources and processing*.

*Supplementary Table 21. Inferred demographic model parameters in different datasets*.

## References

1. Seplyarskiy, V.B. & Sunyaev, S. The origin of human mutation in light of genomic data. Nat Rev Genet 22, 672–686 (2021).

2. Segurel, L., Wyman, M.J. & Przeworski, M. Determinants of mutation rate variation in the human germline. Annu Rev Genomics Hum Genet 15, 47–70 (2014).

3. Gonzalez-Perez, A., Sabarinathan, R. & Lopez-Bigas, N. Local determinants of the mutational landscape of the human genome. Cell 177, 101–114 (2019).

4. Carlson, J. et al. Extremely rare variants reveal patterns of germline mutation rate heterogeneity in humans. Nat Commun 9, 3753 (2018).

5. Fang, Y., Deng, S. & Li, C. A generalizable deep learning framework for inferring fine-scale germline mutation rate maps. Nature Machine Intelligence 4, 1209–1223 (2022).

6. Seplyarskiy, V. et al. A mutation rate model at the basepair resolution identifies the mutagenic effect of polymerase III transcription. Nat Genet 55, 2235–2242 (2023).

7. Neville, M.D.C. et al. Sperm sequencing reveals extensive positive selection in the male germline. Nature (2025).

8. Seplyarskiy, V. et al. Hotspots of human mutation point to clonal expansions in spermatogonia. Nature 647, 429–435 (2025).

9. Grau-Bove, X. et al. A phylogenetic and proteomic reconstruction of eukaryotic chromatin evolution. Nat Ecol Evol 6, 1007–1023 (2022).

10. Malik, H.S. & Henikoff, S. Phylogenomics of the nucleosome. Nat Struct Biol 10, 882–91 (2003).

11. Dai, J. et al. Probing nucleosome function: a highly versatile library of synthetic histone H3 and H4 mutants. Cell 134, 1066–78 (2008).

12. Bagert, J.D. et al. Oncohistone mutations enhance chromatin remodeling and alter cell fates. Nat Chem Biol 17, 403–411 (2021).

13. Jena, S.G. et al. Engineered histones reshape chromatin in human cells. bioRxiv (2025).

14. Jinks-Robertson, S. & Bhagwat, A.S. Transcription-associated mutagenesis. Annu Rev Genet 48, 341–59 (2014).

15. Thornlow, B.P. et al. Transfer RNA genes experience exceptionally elevated mutation rates. Proc Natl Acad Sci U S A 115, 8996–9001 (2018).

16. Marzluff, W.F., Wagner, E.J. & Duronio, R.J. Metabolism and regulation of canonical histone mRNAs: life without a poly(A) tail. Nat Rev Genet 9, 843–54 (2008).

17. Nacev, B.A. et al. The expanding landscape of ‘oncohistone’ mutations in human cancers. Nature 567, 473–478 (2019).

18. Knapp, K., Naik, N., Ray, S., van Haaften, G. & Bicknell, L.S. Histones: coming of age in Mendelian genetic disorders. J Med Genet 60, 213–222 (2023).

19. Lubin, E.E. et al. Coupling deep phenotypic quantification with next-generation phenotyping for 192 individuals with germline histonopathies. HGG Adv 6, 100440 (2025).

20. Al Ojaimi, M. et al. Molecular and clinical aspects of histone-related disorders. Hum Genomics 19, 47 (2025).

21. Chen, S. et al. A genomic mutational constraint map using variation in 76,156 human genomes. Nature 625, 92–100 (2024).

22. Eirín-López, J.M., González-Romero, R., Dryhurst, D., Méndez, J. & Ausió, J. Long-term evolution of histone families: Old notions and new insights into their mechanisms of diversification across eukaryotes. in Evolutionary Biology: Concept, Modeling, and Application (ed. Pontarotti, P.) 139–162 (Springer Berlin Heidelberg, Berlin, Heidelberg, 2009).

23. Yamane, A. et al. RPA accumulation during class switch recombination represents 5’-3’ DNA-end resection during the S-G2/M phase of the cell cycle. Cell Rep 3, 138–47 (2013).

24. Qian, J. et al. B cell super-enhancers and regulatory clusters recruit AID tumorigenic activity. Cell 159, 1524–37 (2014).

25. McCann, J.L. et al. APOBEC3B regulates R-loops and promotes transcription-associated mutagenesis in cancer. Nat Genet 55, 1721–1734 (2023).

26. Petljak, M. et al. Mechanisms of APOBEC3 mutagenesis in human cancer cells. Nature 607, 799–807 (2022).

27. Seplyarskiy, V.B., Andrianova, M.A. & Bazykin, G.A. APOBEC3A/B-induced mutagenesis is responsible for 20% of heritable mutations in the TpCpW context. Genome Res 27, 175–184 (2017).

28. Dai, P. et al. Transcription-coupled AID deamination damage depends on ELOF1-associated RNA polymerase II. Mol Cell 85, 1280–1295 e9 (2025).

29. Landry, S., Narvaiza, I., Linfesty, D.C. & Weitzman, M.D. APOBEC3A can activate the DNA damage response and cause cell-cycle arrest. EMBO Rep 12, 444–50 (2011).

30. Yang, Y., Liu, N. & Gong, L. An overview of the functions and mechanisms of APOBEC3A in tumorigenesis. Acta Pharm Sin B 14, 4637–4648 (2024).

31. Jin, H. et al. Accurate and sensitive mutational signature analysis with MuSiCal. Nat Genet 56, 541–552 (2024).

32. Moore, L. et al. The mutational landscape of human somatic and germline cells. Nature 597, 381–386 (2021).

33. Spisak, N., de Manuel, M., Milligan, W., Sella, G. & Przeworski, M. The clock-like accumulation of germline and somatic mutations can arise from the interplay of DNA damage and repair. PLoS Biol 22, e3002678 (2024).

34. Spisak, N., de Manuel, M. & Przeworski, M. Collateral mutagenesis funnels multiple sources of DNA damage into a ubiquitous mutational signature. bioRxiv (2025).

35. Bennett, R.L. et al. A mutation in histone H2B represents a new class of oncogenic driver. Cancer Discov 9, 1438–1451 (2019).

36. Weinberg, D.N., Allis, C.D. & Lu, C. Oncogenic mechanisms of histone H3 mutations. Cold Spring Harb Perspect Med 7, a026443 (2017).

37. Tessadori, F. et al. Recurrent *de novo* missense variants across multiple histone H4 genes underlie a neurodevelopmental syndrome. Am J Hum Genet 109, 750–758 (2022).

38. Bryant, L. et al. Histone H3.3 beyond cancer: Germline mutations in Histone 3 Family 3A and 3B cause a previously unidentified neurodegenerative disorder in 46 patients. Sci Adv 6, eabc9207 (2020).

39. Tessadori, F. et al. Germline mutations affecting the histone H4 core cause a developmental syndrome by altering DNA damage response and cell cycle control. Nat Genet 49, 1642–1646 (2017).

40. Zeng, T., Spence, J.P., Mostafavi, H. & Pritchard, J.K. Bayesian estimation of gene constraint from an evolutionary model with gene features. Nat Genet 56, 1632–1643 (2024).

41. Traynelis, J. et al. Optimizing genomic medicine in epilepsy through a gene-customized approach to missense variant interpretation. Genome Res 27, 1715–1729 (2017).

42. Sun, K.Y. et al. A deep catalogue of protein-coding variation in 983,578 individuals. Nature 631, 583–592 (2024).

43. Cheng, J. et al. Accurate proteome-wide missense variant effect prediction with AlphaMissense. Science 381, eadg7492 (2023).

44. Meier, J. et al. Language models enable zero-shot prediction of the effects of mutations on protein function. Advances in Neural Information Processing Systems 34 (Neurips 2021) 34(2021).

45. Ioannidis, N.M. et al. REVEL: An ensemble method for predicting the pathogenicity of rare missense variants. Am J Hum Genet 99, 877–885 (2016).

46. Chao, K.R., et al. The landscape of regional missense mutational intolerance quantified from 125,748 exomes. bioRxiv (2024).

47. Li, M., et al. ProSST: Protein language modeling with quantized structure and disentangled attention. bioRxiv (2024).

48. Hu, Q. et al. Mechanisms of BRCA1-BARD1 nucleosome recognition and ubiquitylation. Nature 596, 438–443 (2021).

49. Kaplanis, J. et al. Evidence for 28 genetic disorders discovered by combining healthcare and research data. Nature 586, 757–762 (2020).

50. Foreman, J. et al. DECIPHER: Supporting the interpretation and sharing of rare disease phenotype-linked variant data to advance diagnosis and research. Hum Mutat 43, 682–697 (2022).

51. Fu, J.M. et al. Rare coding variation provides insight into the genetic architecture and phenotypic context of autism. Nat Genet 54, 1320–1331 (2022).

52. Fridman, H., Khazeeva, G., Levy-Lahad, E., Gilissen, C. & Brunner, H.G. Reproductive and cognitive phenotypes in carriers of recessive pathogenic variants. Nat Hum Behav 9, 1726–1736 (2025).

53. van Alten, S., Domingue, B.W., Faul, J., Galama, T. & Marees, A.T. Reweighting UK Biobank corrects for pervasive selection bias due to volunteering. Int J Epidemiol 53, dyae054 (2024).

54. Fry, A. et al. Comparison of sociodemographic and health-related characteristics of UK Biobank participants with those of the general population. Am J Epidemiol 186, 1026–1034 (2017).

55. Brayne, C. & Moffitt, T.E. The limitations of large-scale volunteer databases to address inequalities and global challenges in health and aging. Nat Aging 2, 775–783 (2022).

56. Pollard, K.S., Hubisz, M.J., Rosenbloom, K.R. & Siepel, A. Detection of nonneutral substitution rates on mammalian phylogenies. Genome Res 20, 110–21 (2010).

57. Siepel, A. et al. Evolutionarily conserved elements in vertebrate, insect, worm, and yeast genomes. Genome Res 15, 1034–50 (2005).

58. Davydov, E.V. et al. Identifying a high fraction of the human genome to be under selective constraint using GERP++. PLoS Comput Biol 6, e1001025 (2010).

59. Endler, J.A. Natural Selection in the Wild, (Princeton University Press, Princeton, NJ, 1986).

60. Jiang, S. et al. Generic Diagramming Platform (GDP): a comprehensive database of high-quality biomedical graphics. Nucleic Acids Res 53, D1670–D1676 (2025).

61. Kimura, M. & Ota, T. The average number of generations until extinction of an individual mutant gene in a finite population. Genetics 63, 701–9 (1969).

62. Wissing, S., Montano, M., Garcia-Perez, J.L., Moran, J.V. & Greene, W.C. Endogenous APOBEC3B restricts LINE-1 retrotransposition in transformed cells and human embryonic stem cells. J Biol Chem 286, 36427–37 (2011).

63. Schreck, S. et al. Activation-induced cytidine deaminase (AID) is expressed in normal spermatogenesis but only infrequently in testicular germ cell tumours. J Pathol 210, 26–31 (2006).

64. Popp, C. et al. Genome-wide erasure of DNA methylation in mouse primordial germ cells is affected by AID deficiency. Nature 463, 1101–5 (2010).

65. Martin, A. & Scharff, M.D. Somatic hypermutation of the AID transgene in B and non-B cells. Proc Natl Acad Sci U S A 99, 12304–8 (2002).

66. Zhang, J. & Qian, W. Functional synonymous mutations and their evolutionary consequences. Nat Rev Genet 26, 789–804 (2025).

67. Venkatesh, S.S. et al. Genome-wide analyses identify 25 infertility loci and relationships with reproductive traits across the allele frequency spectrum. Nat Genet 57, 1107–1118 (2025).

68. Matheson-Grant, T. Investigating a dose-dependent threshold for histone H4 pathogenesis in neurodevelopment. Univ. of Otago (2024).

69. Mudge, J.M. et al. GENCODE 2025: reference gene annotation for human and mouse. Nucleic Acids Res 53, D966–D975 (2025).

70. Seal, R.L. et al. A standardized nomenclature for mammalian histone genes. Epigenetics Chromatin 15, 34 (2022).

71. Camacho, C. et al. BLAST+: architecture and applications. BMC Bioinformatics 10, 421 (2009).

72. Nawrocki, E.P. & Eddy, S.R. Infernal 1.1: 100-fold faster RNA homology searches. Bioinformatics 29, 2933–5 (2013).

73. Katoh, K., Misawa, K., Kuma, K. & Miyata, T. MAFFT: a novel method for rapid multiple sequence alignment based on fast Fourier transform. Nucleic Acids Res 30, 3059–66 (2002).

74. Busch, A., Richter, A.S. & Backofen, R. IntaRNA: efficient prediction of bacterial sRNA targets incorporating target site accessibility and seed regions. Bioinformatics 24, 2849–56 (2008).

75. Pockrandt, C., Alzamel, M., Iliopoulos, C.S. & Reinert, K. GenMap: ultra-fast computation of genome mappability. Bioinformatics 36, 3687–3692 (2020).

76. Ramirez, F., Dundar, F., Diehl, S., Gruning, B.A. & Manke, T. deepTools: a flexible platform for exploring deep-sequencing data. Nucleic Acids Res 42, W187–91 (2014).

77. Lecluze, E. et al. Dynamics of the transcriptional landscape during human fetal testis and ovary development. Hum Reprod 35, 1099–1119 (2020).

78. Patro, R., Duggal, G., Love, M.I., Irizarry, R.A. & Kingsford, C. Salmon provides fast and bias-aware quantification of transcript expression. Nat Methods 14, 417–419 (2017).

79. Robinson, M.D., McCarthy, D.J. & Smyth, G.K. edgeR: a Bioconductor package for differential expression analysis of digital gene expression data. Bioinformatics 26, 139–40 (2010).

80. Luo, Y. et al. New developments on the Encyclopedia of DNA Elements (ENCODE) data portal. Nucleic Acids Res 48, D882–D889 (2020).

81. Chen, S. Ultrafast one-pass FASTQ data preprocessing, quality control, and deduplication using fastp. Imeta 2, e107 (2023).

82. Vasimuddin, M., Misra, S., Li, H. & Aluru, S. Efficient architecture-aware acceleration of BWA-MEM for multicore systems. in 2019 IEEE International Parallel and Distributed Processing Symposium (IPDPS) 314–324 (2019).

83. McKenna, A. et al. The Genome Analysis Toolkit: a MapReduce framework for analyzing next-generation DNA sequencing data. Genome Res 20, 1297–303 (2010).

84. Pedersen, B.S. & Quinlan, A.R. Mosdepth: quick coverage calculation for genomes and exomes. Bioinformatics 34, 867–868 (2018).

85. Garcia, M. et al. Sarek: A portable workflow for whole-genome sequencing analysis of germline and somatic variants. F1000Res 9, 63 (2020).

86. Conomos, M.P., Reiner, A.P., Weir, B.S. & Thornton, T.A. Model-free estimation of recent genetic relatedness. Am J Hum Genet 98, 127–48 (2016).

87. Palsson, G. et al. Complete human recombination maps. Nature 639, 700–707 (2025).

88. Wang, K., Li, M. & Hakonarson, H. ANNOVAR: functional annotation of genetic variants from high-throughput sequencing data. Nucleic Acids Res 38, e164 (2010).

89. Sun, S., Wang, Y., Maslov, A.Y., Dong, X. & Vijg, J. SomaMutDB: a database of somatic mutations in normal human tissues. Nucleic Acids Res 50, D1100–D1108 (2022).

90. Emms, D.M. & Kelly, S. OrthoFinder: phylogenetic orthology inference for comparative genomics. Genome Biol 20, 238 (2019).

91. Zhang, Z. et al. ParaAT: a parallel tool for constructing multiple protein-coding DNA alignments. Biochem Biophys Res Commun 419, 779–81 (2012).

92. Wang, D., Zhang, Y., Zhang, Z., Zhu, J. & Yu, J. KaKs_Calculator 2.0: a toolkit incorporating gamma-series methods and sliding window strategies. Genomics Proteomics Bioinformatics 8, 77–80 (2010).

93. Kokic, G., Wagner, F.R., Chernev, A., Urlaub, H. & Cramer, P. Structural basis of human transcription-DNA repair coupling. Nature 598, 368–372 (2021).

94. Alexandrov, L.B. et al. The repertoire of mutational signatures in human cancer. Nature 578, 94–101 (2020).

95. Liu, M.H. et al. DNA mismatch and damage patterns revealed by single-molecule sequencing. Nature 630, 752–761 (2024).

96. Mathieson, I. & Reich, D. Differences in the rare variant spectrum among human populations. PLoS Genet 13, e1006581 (2017).

97. Langmead, B. & Salzberg, S.L. Fast gapped-read alignment with Bowtie 2. Nat Methods 9, 357–9 (2012).

98. Tarasov, A., Vilella, A.J., Cuppen, E., Nijman, I.J. & Prins, P. Sambamba: fast processing of NGS alignment formats. Bioinformatics 31, 2032–4 (2015).

99. Zhang, Y. et al. Model-based analysis of ChIP-Seq (MACS). Genome Biol 9, R137 (2008).

100. Zhao, H. et al. CrossMap: a versatile tool for coordinate conversion between genome assemblies. Bioinformatics 30, 1006–7 (2014).

101. Abramson, J. et al. Accurate structure prediction of biomolecular interactions with AlphaFold 3. Nature 630, 493–500 (2024).

102. Dombrowski, M., Engeholm, M., Dienemann, C., Dodonova, S. & Cramer, P. Histone H1 binding to nucleosome arrays depends on linker DNA length and trajectory. Nat Struct Mol Biol 29, 493–501 (2022).

103. Pettersen, E.F. et al. UCSF ChimeraX: Structure visualization for researchers, educators, and developers. Protein Sci 30, 70–82 (2021).

104. Jubb, H.C. et al. Arpeggio: A web server for calculating and visualising interatomic interactions in protein structures. J Mol Biol 429, 365–371 (2017).

105. Armeev, G.A., Kniazeva, A.S., Komarova, G.A., Kirpichnikov, M.P. & Shaytan, A.K. Histone dynamics mediate DNA unwrapping and sliding in nucleosomes. Nat Commun 12, 2387 (2021).

106. Mitternacht, S. FreeSASA: An open source C library for solvent accessible surface area calculations. F1000Res 5, 189 (2016).

107. Tien, M.Z., Meyer, A.G., Sydykova, D.K., Spielman, S.J. & Wilke, C.O. Maximum allowed solvent accessibilites of residues in proteins. PLoS One 8, e80635 (2013).

108. McLaren, W. et al. The Ensembl Variant Effect Predictor. Genome Biol 17, 122 (2016).

109. Tran, T.C. et al. PheWAS analysis on large-scale biobank data with PheTK. Bioinformatics 41, btae719 (2024).

110. Koscielny, G. et al. Open Targets: a platform for therapeutic target identification and validation. Nucleic Acids Res 45, D985–D994 (2017).

111. Sanders, S.J., et al. *De novo* mutations revealed by whole-exome sequencing are strongly associated with autism. Nature 485, 237–41 (2012).

112. Danecek, P. et al. Twelve years of SAMtools and BCFtools. Gigascience 10(2021).

113. Gutenkunst, R.N., Hernandez, R.D., Williamson, S.H. & Bustamante, C.D. Inferring the joint demographic history of multiple populations from multidimensional SNP frequency data. PLoS Genet 5, e1000695 (2009).

114. Kim, B.Y., Huber, C.D. & Lohmueller, K.E. Inference of the distribution of selection coefficients for new nonsynonymous mutations using large samples. Genetics 206, 345–361 (2017).

115. Rodriguez-Galindo, M., Casillas, S., Weghorn, D. & Barbadilla, A. Germline *de novo* mutation rates on exons versus introns in humans. Nat Commun 11, 3304 (2020).

